# Influence of ionic liquids on enzymatic asymmetric carboligations

**DOI:** 10.1101/2025.05.21.655364

**Authors:** Tina Gerhards, Till El Harrar, Andreas Sebastian Klein, Eric von Lieres, Caroline E. Paul, Iván Lavandera, Vicente Gotor-Fernández, Jörg Pietruszka, Martina Pohl, Holger Gohlke, Dörte Rother

## Abstract

The asymmetric mixed carboligation of aldehydes catalyzed by thiamine diphosphate (ThDP)-dependent enzymes provides a sensitive system for monitoring changes in activity, chemo-, and enantioselectivity. While previous studies have shown that organic cosolvents influence these parameters, we now demonstrate that similar effects occur upon addition of water-miscible ionic liquids (ILs). In this study, six ThDP-dependent enzymes were analyzed in the presence of 14 ILs under comparable conditions to assess their influence on enzymatic carboligation reactions yielding 2-hydroxy ketones. ILs exerted a moderate to strong influence on activity and, more notably, altered enantioselectivity. (*R*)-selective reactions were generally stable upon IL addition, while (*S*)-selective reactions frequently showed reduced selectivity or even inversion to the (*R*)-enantiomer. The most significant change was observed for the *Ap*PDC_E469G variant of pyruvate decarboxylase from *Acetobacter pasteurianus*, where the enantiomeric excess shifted from 86% (*S*) to 60% (*R*) in the presence of 9% (w/v) Ammoeng 102. Control experiments indicated that this shift was primarily due to the Ammoeng cation rather than the anion. To explore the molecular basis of this phenomenon, all-atom molecular dynamics (MD) simulations were performed on wild-type *Ap*PDC and the E469G variant in Ammoeng 101 and Ammoeng 102. The simulations revealed that hydrophobic and hydrophilic regions of the Ammoeng cations interact with the (*S*)-selective binding pocket, thereby favoring formation of the (*R*)-product. These results highlight the potential of solvent engineering for modulating enzyme selectivity and demonstrate that MD simulations can capture functionally relevant enzyme–solvent interactions at the atomic level.

## 1. Introduction

Thiamine diphosphate (ThDP)-dependent enzymes catalyze the C-C coupling of two aldehydes to form 2-hydroxy ketones. In the bimolecular mixed carboligation of benzaldehyde and acetaldehyde, four pairs of enantiomeric products are possible, depending on which aldehyde acts as a donor or acceptor [1,2]. The chemoselectivity of ThDP-dependent enzymes is determined by the binding sequence of the two aldehyde substrates to the cofactor ThDP, where the first substrate to bind is called the donor and the second substrate is called the acceptor [1,3] (**Figure 1**).

**Figure 1:**
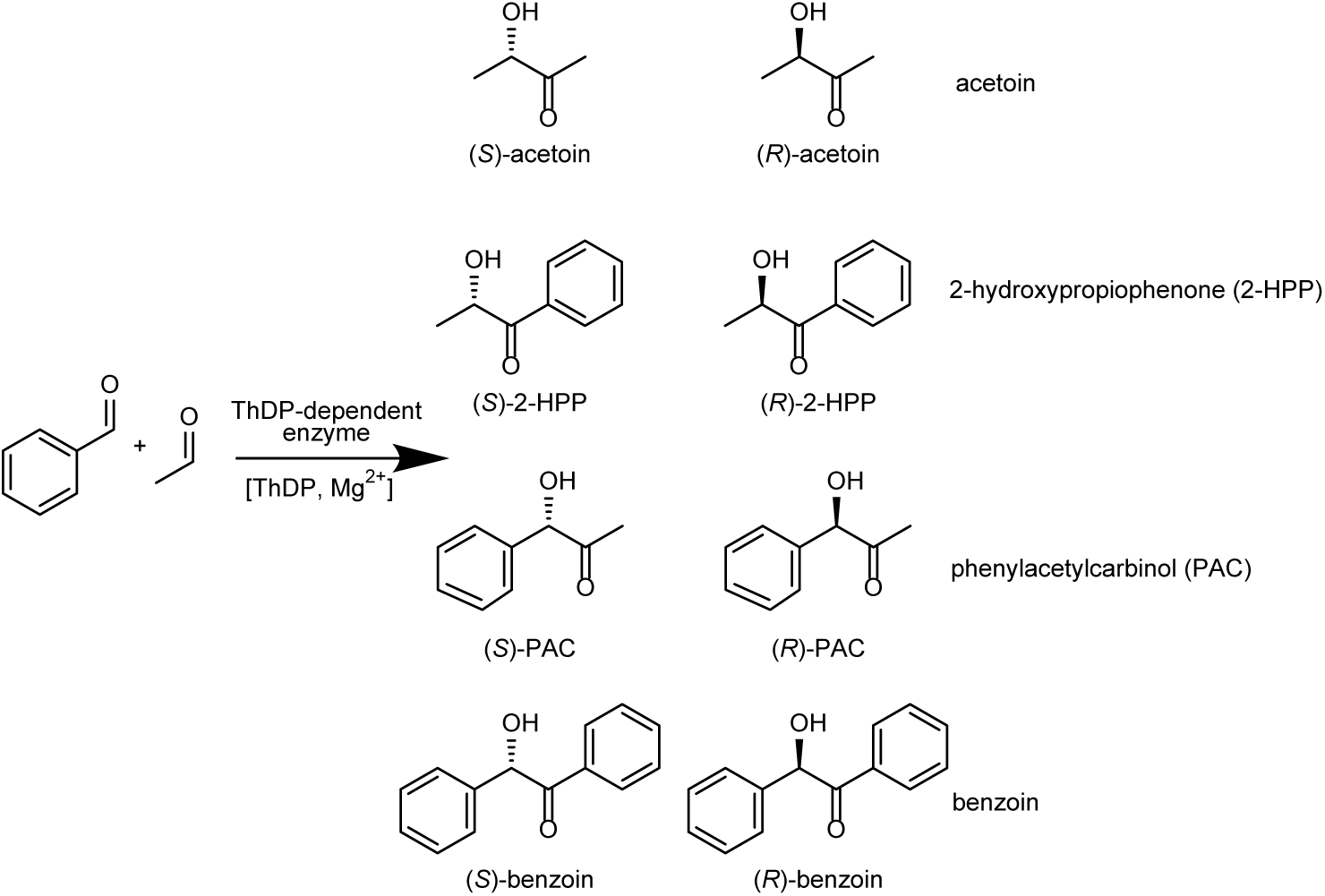
The four possible pairs of enantiomeric products derived from the carboligation of acetaldehyde and benzaldehyde catalyzed by ThDP-dependent enzymes.

The relative orientation of the two substrates prior to C-C-coupling determines the enantioselectivity of these enzymes. The parallel orientation of the two aldehydes with respect to the carbonyl groups and side chains prior to C-C-bond formation allows the formation of the (*R*)-enantiomer. In contrast, the formation of the (*S*)-enantiomer requires the antiparallel orientation of the two aldehydes, which is only possible in ThDP-dependent enzymes that have a special feature in their active site, the so-called (*S*)-pocket [4–7]. High chemo-as well as stereoselectivities have been achieved in several wild-type enzymes and variants by optimal stabilization of both substrates in the active site [1,7–9].

Although the overall three-dimensional structure of ThDP-dependent enzymes is highly conserved, the different spatial conditions in the donor and acceptor binding sites lead to different products with predominantly high enantioselectivity [1,10]. Because the structure-function relationship is well understood, including the orientation of the two substrates to achieve high enantioselectivity and chemoselectivity, ThDP-dependent enzymes are a reliable and sensitive system to study the effects of non-conventional media. Even small differences in the three-dimensional structure of the active sites lead to pronounced shifts in the chemoselectivity and enantioselectivity of this family of biocatalysts.

Previously, we have extensively investigated the influence of organic cosolvents on the activity and selectivity of ThDP-dependent enzymes [11]. The presence of different organic cosolvents in concentrations up to 30% (v/v) often shifted the chemoselectivity towards the smaller product (e.g. 2-hydroxypropiophenone, 2-HPP, instead of benzoin, Figure 1). We interpreted the results in terms of (i) flexibility shifts, (ii) changes in substrate and product solubility (and hence kinetic effects), and (iii) selective blockage by direct interactions of solvent molecules with the active site. These effects of (i) to (iii) varied depending on the biocatalyst studied. In terms of chemoselectivity, biotransformations that were rather chemoselective in buffer without solvent addition were less susceptible to alteration in the presence of organic solvents than non-selective biotransformations. A similar effect was observed for enantioselectivity: No observable effect was detected when the selectivity in buffer was already exceptionally high (>99% enantiomeric excess, *ee*). We found the most pronounced shift in enantioselectivity for a variant of pyruvate decarboxylase from *Acetobacter pasteurianus, Ap*PDC_E469G, which catalyzes the synthesis of (*S*)-phenylacetylcarbinol [(*S*)-PAC] with an *ee* of 89% (*S*) in aqueous buffer. The addition of 0.5% (v/v) chloroform resulted in a shift towards the (*R*)-enantiomer (49% *ee*). However, it was not possible to further improve the (*S*)-selectivity by employing other organic cosolvents. We found a clear correlation between the observed shifts in enantioselectivity with the size and polarity of the cosolvent, which could be explained by the selective blocking of the (*S*)-pocket.

In this study, we now demonstrate the influence of ionic liquids (ILs) on the same enzymatic system and show similarly pronounced effects on the activity and selectivity of the bimolecular carboligation reaction of benzaldehyde and acetaldehyde. The enzymes investigated in this study are well described: Benzaldehyde lyase from *Pseudomonas fluorescens* (*Pf*BAL [12,13]), benzoylformate decarboxylase from *Pseudomonas putida* (*Pp*BFD [14]), a variant of this enzyme (*Pp*BFD_H281A [15]), branched-chain 2-ketoacid decarboxylase from *Lactococcus lactis* (*Ll*KdcA [16,17]), pyruvate decarboxylase from *Acetobacter pasteurianus* (*Ap*PDC [18]), and an (*S*)-selective variant thereof (*Ap*PDC_E469G [7]).

In the past decades, ILs were considered as promising green alternative to traditional organic solvents due to their lack of vapor pressure, non-flammability, and excellent chemical stability [19,20]. Later, however, the greenness of ILs was questioned, taking into account the entire life cycle and partially unknown effects on the ecosystem [19,21,22]. Since several preparation steps are required, the purity of the preparations often varies greatly, which can be challenging when reproducing syntheses. In addition, such impurities can affect enzymatic reactions. However, if some general guidelines are followed as comprehensively summarized by Bubalo *et al*. [19] (e.g., prepared from naturally derived materials, low toxicity), the potential of ILs is enormous, considering the unlimited possible combinations of cations and anions.

Regarding the use of ILs in biotransformations [23–25], it has been claimed for years that they do not inactivate enzymes as many organic solvents do [26]. As more and more applications have become known, studies have also been performed showing inactivation and subsequent destabilization of the selected biocatalysts. This was initially attributed to strong enzyme-IL-ion interactions of individual cation [27] or anion [28] species with the respective oppositely charged enzyme residues, which were suggested to induce deactivating structural changes as determined by structure-activity data or MD simulations [29]. While in the case of chymotrypsin, papain, and lipases, stability could be increased by charge modification, recent results indicate that especially less selected but energetically advantageous interactions of IL-ions – often involving uncharged enzyme residues [30] – are equally important in destabilizing the local enzyme structure [31]. In addition, cooperative and competitive ion-ion or enzyme-ion effects can significantly influence enzyme-IL interactions [30,32,33], while the effects on enzyme-residue pair interactions can be interaction-, configuration-, and solvent-specific [34].

ILs have already been implemented as solvents and cosolvents in several biotransformations [35–37], but mostly on a selective basis [20]. Comprehensive data concerning their influence on whole enzyme families is missing. Most studies deal with the addition of ILs in lipase-catalyzed biotransformations. For example, in some studies, the enantioselectivity of lipases was increased by the addition of IL [38,39], even more than with the addition of organic solvents [40,41]. A very detailed study concerning the influence of ILs with respect to the position of the amino acids of the enzyme was published by Frauenkron-Machedjou *et al.* [42]. Saturation mutagenesis at each position of the gene encoding *Bacillus subtilis* lipase A revealed that improvement in resistance could be achieved at 50–69 % of all amino acid positions in the presence of four different ILs. This dataset was used to extend our knowledge of guidelines for protein engineering following a data-driven approach [43] and to critically assess structure-based approaches to improve the protein’s resistance toward aqueous ILs [31].

First effects of ILs on ThDP-dependent enzymes have already been investigated by Pohl and coworkers [44]. In this study, the enzymes *Pf*BAL and the *Pp*BFD variant H281A were tested. The initial reaction rate of *Pf*BAL for the benzoin ligation was improved by the addition of Ecoeng 21M and Ecoeng 111P. Furthermore, the stereoselective formation of (*R*)-2-hydroxypropiophenone [(*R*)-2-HPP] catalyzed by both enzymes was more selective in the presence of 50% (v/v) (for *Pf*BAL) and 40% (v/v) (for *Pp*BFD_H281A) IL [44]. However, these initial results are not sufficient to understand the general effects of ILs on carboligation reactions and to extrapolate general trends to other biotransformations using this group of enzymes. Therefore, we present here a more comprehensive study involving six ThDP-dependent enzymes (differing in substrate preference and enantioselectivity) in the presence of fourteen different chiral and achiral water-miscible ILs (Figure 2) diluted in aqueous buffer at three different concentrations. From the wide range of data on IL-induced shifts in chemo- and enantioselectivity, general trends were identified and interpreted with the help of all-atom molecular dynamics (MD) simulations in terms of structure-function relationships and structural dynamics.

**Figure 2:**
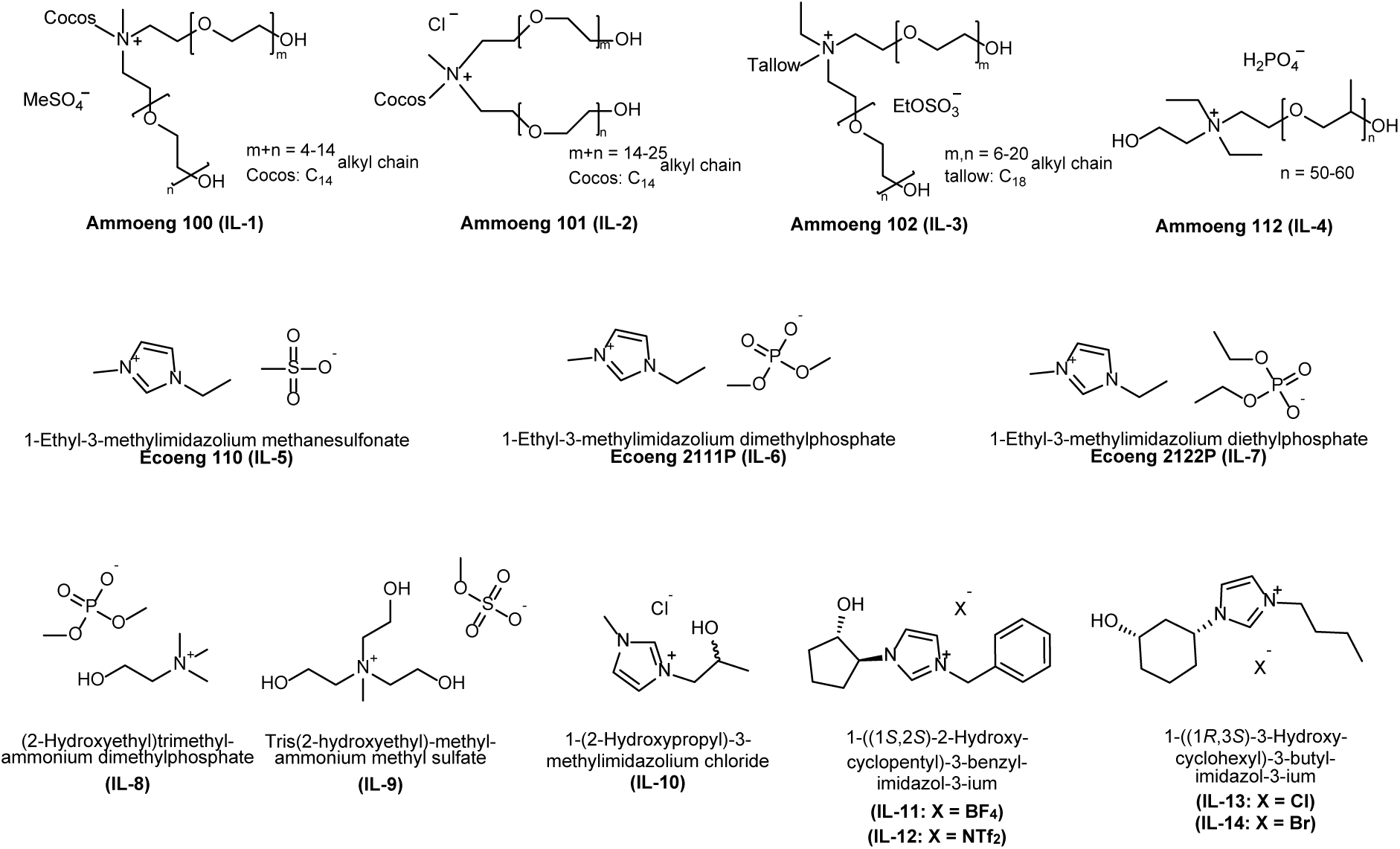
ILs tested as additives in this study.

## 2. Materials, Methods, and Computational Details

### 2.1 Enzymes

All enzymes were produced and purified as described elsewhere (*Pf*BAL [45], *Pp*BFD (and *Pp*BFD_H281A) [14,15], *Ll*KdcA [16], *Ap*PDC (and *Ap*PDC_E469G) [7].

### 2.2 Reactions

All reactions were performed at an 800 µL scale. The substrates benzaldehyde (18 mM final concentration) and acetaldehyde (18 mM (*Ap*PDC, *Ap*PDC_E469G, *Pf*BAL) / 180 mM (*Ll*KdcA, *Pp*BFD, *Pp*BFD_H281A) final concentration) were dissolved in triethanolamine (TEA) buffer (50 mM, 0.1 mM ThDP, 2.5 mM MgSO_4_), pH 7.5 (for all enzymes except *Pf*BAL) or pH 8.0 (for *Pf*BAL). 0.02 mg/mL (*Pf*BAL) / 0.1 mg/mL (for all other enzymes) was used. If the productivity was so low that the enantioselectivity could not be determined reliably, the approach was repeated with 0.4 mg/mL enzyme. The catalyst was dissolved in buffer and finally added to the reaction mixture. Reactions were performed in a thermo mixer (Eppendorf, Germany) for 24 h at 20°C. The reactions were performed in one to four replicates (pronounced effects were detected several times to get solid data). The error bars are given in the plots in the supporting information. Reaction batches containing ILs were prepared as described above, but the volume of the buffer was reduced according to the volume of the respective solvent.

### 2.3 Sample preparation and analysis

The samples were prepared and measured as described before [11,46]. All changes in reactions with IL were detected relative to a reaction with the same reaction conditions but without IL addition.

### 2.4 Ionic liquids

Figure 2 shows the ILs used. IL-1 and IL-3 were supplied by IoLiTec Ionic Liquids Technologies GmbH, IL-2, IL-5, IL-6, and IL-7 were from Solvent Innovation (now Merck Chemicals), IL-9 was obtained from Sigma-Aldrich and IL-4 was kindly provided by Dr. Schwab from Evonik Goldschmidt GmbH. Based on previous studies [45,46], they were added to the biotransformations with a concentration of 9%, 2.5%, and 1% (w/v). To obtain a single-phasic system, the solubility was tested beforehand. Only IL-12 was not well soluble and therefore tested in concentrations of 0.1%, 0.5%, and 0.9% (w/v).

The ionic liquids IL-10 to IL-14 were prepared as follows:

**IL-10** was prepared as described in the literature [49]. Light yellow oil. **^1^H-NMR** (300 MHz, D_2_O) δ 8.64 (s, 1H), 7.40 (d, *J* = 1.7 Hz, 1H), 7.36 (d, *J* = 1.9 Hz, 1H), 4.23 (dd, *J* = 13.6, 2.7 Hz, 1H), 4.13-4.03 (m, 1H), 4.00 (dd, *J* = 13.6, 7.0 Hz, 1H), 3.81 (s, 3H), 1.14 (d, *J* = 6.2 Hz, 3H); **^13^C-NMR** (75 MHz, D_2_O) δ 137.4, 123.3, 122.8, 65.8, 55.6, 35.6, 18.9; **MS** (ESI^+^, *m*/*z*): 142 [(M+H)^+^, 10%], 141 (M^+^, 100%).

**IL-11** was prepared as described in the literature [50]. Light yellow oil. **^1^H-NMR** (300 MHz, CDCl_3_): δ 1.66-1.95 (m, 4H), 2.04-2.11 (m, 1H), 2.26-2.34 (m, 1H), 3.80 (s, 1H), 4.34 (q, *J* =7.9 Hz, 1H), 4.54 (q, *J* = 7.9 Hz, 1H), 5.40 (s, 2H), 7.25 (s, 1H), 7.35-7.43 (m, 6H), 9.49 (s, 1H); **^13^C-NMR** (75 MHz, CDCl_3_): δ 18.4, 27.8, 30.3, 52.5, 66.3, 76.5, 120.2, 121.0, 129.5, 128.2, 128.5, 131.9, 134.5; **MS** (ESI^+^, *m/z*): 244 [(M+H)^+^, 20%], 243 [M^+^, 100%]; (ESI^-^, m/z): 87 [BF_4_^-^, 100%]; **[α]_D_^20^** = +14.9 (c = 1.0, CHCl_3_) for > 99% *ee*.

**IL-12** was prepared as described in the literature [50]. Light yellow oil. **^1^H-NMR** (300 MHz, CDCl_3_): δ 1.62-1.75 (m, 1H), 1.83-1.99 (m, 3H), 2.07-2.19 (m, 1H), 2.34-2.41 (m, 1H), 4.27 (q, *J* = 7.9 Hz, 1H), 4.39 (q, *J* = 7.9 Hz, 1H), 5.31 (s, 2H), 7.18 (s, 1H), 7.26-7.40 (m, 6H), 8.80 (s, 1H); **^13^C-NMR** (75 MHz, CDCl_3_): δ 19.0, 28.2, 31.1, 53.6, 67.1, 77.2, 120.9, 122.0, 129.5, 128.9, 129.7, 131.9, 134.3; **MS** (ESI^+^, *m/z*): 244 [(M+H)^+^, 18%], 243 [M^+^, 100%]; (ESI^-^, m/z): 280 [NTf_2_^-^, 100%]; **[α]_D_^20^** = +16.5 (c = 1.0, CHCl_3_) for > 99% *ee*.

**IL-13** was prepared as described in the literature [51]. Colorless oil. **^1^H-NMR** (300 MHz, CD_3_OD) δ 9.14 (s, 1H), 7.77 (s, 1H), 7.68 (s, 1H), 4.39 (ap tt, *J* = 12.2, 3.9 Hz, 1H), 4.23 (t, *J* = 7.4 Hz, 3H), 3.73 (ap tt, *J* = 10.9, 4.2 Hz, 1H), 2.44 (ap ddq, *J* = 9.6, 3.9, 2.0 Hz, 1H), 2.15 (ap d, *J* = 11.9 Hz, 1H), 2.05-1.84 (m, 4H), 1.76-1.62 (m, 2H), 1.58-1.44 (m, 1H), 1.42-1.19 (m, 3H), 0.99 (t, *J* = 7.4 Hz, 3H); **^13^C-NMR** (75 MHz, CD_3_OD) δ 136.0, 123.8, 122.2, 69.3, 59.3, 50.7, 42.7, 35.0, 33.3, 33.1, 22.7, 20.5, 13.7; **MS** (ESI^+^, *m/z*): 223 [M^+^, 100%], 224 [(M+H)^+^, 15%]; **[α]_D_^20^** = −7.2 (c = 1.0, MeOH) for > 99% *ee*.

**IL-14** was prepared as described in the literature [51]. White powder. **^1^H-NMR** (300 MHz, CD_3_OD) δ 9.16 (s, 1H), 7.78 (s, 1H), 7.68 (s, 1H), 4.41 (ap tt, *J* = 12.2, 3.8 Hz, 1H), 4.24 (t, *J* = 7.4 Hz, 2H), 3.73 (ap tt, *J* = 10.9, 4.1 Hz, 1H), 2.44 (ap d, *J* = 11.4 Hz, 1H), 2.16 (ap d, *J* = 12.0 Hz, 1H), 2.04-1.84 (m, 4H), 1.77-1.63 (m, 2H), 1.59-1.44 (m, 1H), 1.43-1.24 (m, 3H), 0.99 (t, *J* = 7.4 Hz, 3H); **^13^C-NMR** (75 MHz, CD_3_OD) δ 136.0, 123.7, 122.2, 69.3, 59.3, 50.8, 42.8, 35.0, 33.3, 33.1, 22.7, 20.5, 13.8; **MS** (ESI^+^, *m/z*): 223 [M^+^, 100%], 224 [(M+H)^+^, 20%]; **[α]_D_^20^** = −7.8 (c = 1.0, MeOH) for > 99% *ee*.

### 2.5 Instrumental analysis of IL-3

IL-3 was analyzed using GC/MS, ^1^H-NMR, ^13^C-NMR and IR. For details see SI (**Figures S14-S16**).

### 2.6 Molecular dynamics simulations

Three sets of unbiased molecular dynamics (MD) simulations of the (*R*)-selective *Ap*PDC wild type (*Ap*PDC_WT) and the (*S*)-selective *Ap*PDC_E469G variant were performed in water and up to two ILs, in addition to one set of steered MD simulations.

First, unbiased MD simulations of homotetramers based on a crystal structure (PDB-ID: 2VBI) of both enzymes were performed in water and 0.08 M Ammoeng 102 (IL-3) with ions placed randomly in the simulation box, as this IL showed the highest influence on carboligation enantioselectivity. To screen for the influence of distinct structural elements of the IL ion, we additionally simulated *Ap*PDC_E469G in the structurally similar IL Ammoeng 101 (IL-2) at the same concentration, which showed a lower effect on enantioselectivity in *Ap*PDC_E469G. Note that we used the same concentrations in our simulations for IL-2 and IL-3 in contrast to experimentally determined values [46] to maintain a consistent simulation setup. For each system, 8 (*Ap*PDC_WT) or 12 (*Ap*PDC_E469G) independent replica simulations were conducted, each of 8 µs length, resulting in a total of 416 µs of simulation time. For trajectory analyses, the first 1000 ns were discarded as equilibration time.

Second, unbiased MD simulations were performed based on a snapshot obtained in the previous simulations of *Ap*PDC_E469G in IL-3 for tetramers of *Ap*PDC_WT and *Ap*PDC_E469G with the hydrophobic moiety of the IL-3 cation present in the substrate tunnels of the first of the two active sites of each dimer and without an IL-3 cation present in the substrate tunnels. Additionally, the substrates benzaldehyde (BA) and acetaldehyde (AA) were placed in the catalytic site following the arrangement during catalysis oriented in the respective (*R*)-or (*S*)-selective state according to Ref. [7]. The cation and substrate structures were selectively minimized using the function integrated into Schrödinger’s Maestro program suite [52] after positioning them in the substrate tunnel and active sites, respectively, to eliminate steric clashes. For each system, we conducted 8 independent replica simulations, each of 4 µs length, resulting in a total of 128 µs of simulation time. For trajectory analyses, the first 1000 ns were discarded as equilibration time.

Third, as in the previous set of simulations the IL-3 cation showed a repositioning towards the (*S*)-pocket during the end of the relaxation phase due to reduced positional restraints, i.e., resulting in a biased starting configuration, MD simulations were performed starting from the same input structure as for set two but keeping the IL cation restraint until the end of the relaxation phase for dimers of *Ap*PDC_WT and *Ap*PDC_E469G. Additionally, ThDP was modeled as acetaldehyde-ThDP-conjugated iminopyrimidine (IP) state following Paulikat *et al*. [53] with benzaldehyde again being oriented in the respective (*R*)- or (*S*)-selective state to reduce the high mobility of benzaldehyde and acetaldehyde, as benzaldehyde and acetaldehyde displayed high mobility during the previous set of simulations and sometimes left the active sites. The cation and substrate structures were minimized using the function integrated into Schrödinger’s Maestro program suite [52] after positioning them in the substrate tunnels and active sites, respectively, to eliminate steric clashes. For each system, we conducted 12 independent replica simulations, each of 750 ns length, resulting in a total of 36 µs of simulation time. For trajectory analyses, the first 500 ns were discarded as equilibration time.

Fourth, we simulated the approach of benzaldehyde to ThDP by performing 100 steered MD simulations of 31 ns length each for dimers of *Ap*PDC_WT and *Ap*PDC_E469G with and without the hydrophobic moiety of the IL-3 cation located in the (*S*)-pocket and benzaldehyde placed at the entry of the substrate tunnel, resulting in a total of 12.4 µs of simulation time. ThDP was modeled as acetaldehyde-ThDP-conjugated IP state following [53]. For trajectory analyses, only the last snapshot was used for binding mode evaluation.

### 2.7 System preparation

The initial coordinates of the tetrameric *Ap*PDC_WT (including the noncovalently bound cofactor thiamine diphosphate (ThDP) and the Mg^2+^ ion) were obtained from the crystal structure (PDB ID: 2VBI), resolved at a resolution of 2.75 Å. Water molecules within the crystal structures were retained. The *Ap*PDC_E469G variant was obtained using the Modeller Software [54] to perform the E469G substitution in each monomer. Protonation states for all titratable amino acids were assigned according to pH 7.5 using the program Epik [55], which is part of the Protein Preparation Wizard [56] included in Schrödinger’s Maestro program suite [52]. All hydrogen atoms of the crystal structure were removed using the REDUCE program and reassigned with the program LEaP [57] according to the Amber ff19SB library [58], both of which are included in the AMBER20 [59] program package. We added an NME cap group at the C-terminal amino acid to avoid an artificially charged terminus. The force field parameters and atomic partial charges for *Ap*PDC_WT and *Ap*PDC_E469G were taken from the AMBER ff19SB force field [58].

As to the IL ions, the cofactor ThDP, and the substrates benzaldehyde and acetaldehyde, the initial 3D structures were prepared by using the LEaP [57] program from AMBER20 [59] or taken from the crystal structure (ThDP). Our investigations revealed that, to our knowledge, available structures for the IL-3 cations (**Figure S18**) are questionable, as evidenced by findings in IR-, ^13^C-NMR-, and mass spectrometry spectra (see **Figures S14-S16** and **Text S5)**. Thus, we deliberately decided to employ – following our structural analyses – a mass-corrected version of the manufacturer structure of the IL-3 cation with the widely-used IL-3 cation core structure in our simulations to maintain consistency with previous studies (**Figure S18B)**. We carefully tested to what extent uncertain structural parts might impact the validity of our results (see **Figure S17** and **Text S6)**. However, the results should largely be unaffected, as the effects observed in the MD simulations are predominantly influenced by structural features located towards the end of the cation side chains, i.e., regions that are far away from the central cationic core. Overall, the conclusions of our computational studies should not be influenced by the uncertainty of the IL-3 cation structure. The structures were subjected to quantum mechanical (QM) geometry optimization using Gaussian 16 [60] at the HF/6-31G* level of theory [61]. The ThDP structure was prepared as aminopyrimidine (AP) state for the first two sets of unbiased MD simulations and as acetaldehyde-ThDP-conjugated iminopyrimidine (IP) state for the third set of unbiased MD simulations and the steered MD simulations, following ref. [53]. Atomic partial charges were derived according to the RESP [62] procedure. The resulting parameters, e.g., the atomic partial charges, were compared with parameters from dedicated IL force fields or similar studies [63–65], if possible, and found to be in good agreement. The force field parameters for ILs were taken from the second generation of the general AMBER force field (GAFF2) [66]. Packmol [67] was initially used to place one enzyme in the center of a cubic simulation box and then to randomly add the needed amount of the respective cations and anions to reflect the desired concentration, with electroneutrality ensured using additional Na^+^ ions. Since periodic boundary conditions were used, a minimal distance of 15 Å from the protein to the box sides was used to prevent self-interaction of the protein across the box borders. The systems were then solvated using the OPC water model [68], also by using Packmol [67]. Two different but equivalent preparation schemes were used for the MD simulations as described in the following. Initially, for the simulations of *Ap*PDC_E469G in water, IL-2, and IL-3, scheme A was used, whereas scheme B, which was adapted from Ref. [69] was used for the remainder of the simulations.

**Scheme A:** Initially, harmonic restraints with a force constant of 5 kcal mol^-1^ Å^-2^ were applied to all protein atoms for 25 000 cycles (500 cycles of steepest descent (SD) followed by 24 500 cycles of conjugate gradient (CG) minimization). Second, the harmonic restraints were reduced to a force constant of 1 kcal mol^-1^ Å^-2^ for 20 000 cycles (2000 cycles SD and 18 000 cycles CG minimization). Third, 10 000 cycles SD and 40 000 cycles CG minimization without any restraints were performed. In the subsequent thermalization, the system was first heated from 0 K to 100 K over 50 ps in a canonical (NVT) MD simulation. Harmonic restraints of 1 kcal mol^-1^ Å^-2^ were applied on protein atoms, and a time step of 2 fs was used. The temperature was then raised from 100 K to 300 K over 100 ps of isobaric-isothermal (NPT) MD simulations, followed by an additional 2.5 ns of NPT-MD simulations. Finally, the harmonic restraints were reduced to 0 kcal mol^-1^ Å^-2^ over the course of six NPT-MD simulations with a length of 50/250/500/500/500/1000 ps with a step size of 1/1/2/2/2/2 fs, respectively.

**Scheme B:** The systems were first subjected to an energy minimization to eliminate steric clashes for 4000 cycles SD followed by 1000 cycles CG minimization while applying harmonic restraints to protein and IL atoms with a force constant of 5 kcal mol^-1^ Å^-2^. In a subsequent thermalization, the systems were heated from 0 K to 100 K to 200 K to 300 K over three steps of each 5 ps while applying harmonic restraints to protein and IL atoms with a force constant of 5 kcal mol^-1^ Å^-2^ in a canonical (NVT) MD simulation, followed by additional 10 ps at 300 K. Subsequently, two additional minimization steps were applied, analogous to the first step, first with reduced harmonic restraints of 0.5 kcal mol^-1^ Å^-2^, then without restraints, respectively. Following, a second thermalization equal to the first one was applied, with harmonic restraints reduced to 1 kcal mol^-1^ Å^-2^ and an extended final phase of 35 ps. Finally, the harmonic restraints were reduced from 1 kcal mol^-1^ Å^-2^ to 0 kcal mol^-1^ Å^-2^ over the course of four NPT-MD simulations with a length of 50/100/200/2000 ps with a step size of 1/2/4/4 fs, respectively.

The production simulations were performed in the NPT ensemble at 300 K all with a time step of 4 fs using the hydrogen mass repartitioning (HMR) method [70]. Coordinates were saved every 2 ns for the first two sets of unbiased MD simulations and 0.2 ns for the third set of unbiased MD simulations and the steered MD simulations, respectively. All computations were performed using the GPU-accelerated version of pmemd [71] from the AMBER20 [59] program suite.

In all MD simulations, the particle mesh Ewald (PME [72]) method was used to treat long-range electrostatic interactions. The distance cutoff for short-range non-bonded interactions was set to 9 Å. Langevin dynamics were used with a time constant (*τ*) of 0.5 ps for heat bath-coupling to keep the system temperature at the target temperature of 300 K during the simulations. The SHAKE [73] algorithm was applied to all bonds involving hydrogens. To set up the independent MD production simulations, the simulations were assigned random initial velocities at the start of the production run.

### 2.8 Steered MD simulations

In the steered MD simulations, the approach of benzaldehyde from the entrance of the main substrate tunnel found in the crystal structure to the acetaldehyde-ThDP conjugate was performed by pulling the C-atom of the carbonyl function of benzaldehyde towards the C-atom of the hydroxy function of ThDP using a force constant of 50 kcal mol^-1^ and a pulling rate of 0.1 Å ns^-1^. We used harmonic restraints with a force constant of 20 kcal mol^-1^ to ensure a catalytically plausible binding mode that did not confer biases towards a distinct configuration leading to (*S*)- or (*R*)-selective product formation. This was achieved by enforcing a *Si*-face approach of benzaldehyde to the ThDP-bound acetaldehyde in the first half of the simulations, with the aldehyde function approaching first. In the second half, this restraint was gradually reduced from 20 kcal mol^-1^ to 0 kcal mol^-1^ during the remainder of the simulation, while simultaneously introducing two restraints enforcing coplanar and colinear orientation of benzaldehyde with force constants gradually increasing from 0 kcal mol^-1^ to 20 kcal mol^-1^ during the remainder of the simulation. The simulations were performed for dimeric structures of *Ap*PDC_WT and *Ap*PDC_E469G with and without an IL-3 cation present in the (*S*)-pocket of *Ap*PDC_E469G or the active site in *Ap*PDC_WT, respectively. The last frame of each simulation was used for evaluation of the binding orientation.

### 2.9 Trajectory analysis

The first 500 to 1000 ns of each unbiased MD simulation were discarded as equilibration phase, leaving 7 µs, 3 µs, or 250 ns for the trajectory analysis for the first, second, or third set of unbiased MD simulations, respectively. The analyses were performed with cpptraj from the AmberTools20 package [59] or the CAVER software [74]. The following measures were evaluated:

**I)** the root mean-square deviation (RMSD) as a measure of structural similarity (fitted to the backbone atoms of the crystal structure)
**II)** the root mean-square fluctuation (RMSF) as a measure of mobility (fitted to the backbone atoms of the crystal structure)
**III)** solvent density grids
**IV)** intermolecular distances
**V)** spatial product
**VI)** substrate tunnel radius and dynamics

Results over all independent trajectories of the same system are shown as means ± standard errors of the mean (SEM) and were analyzed with the SciPy [75] and Pandas [76] libraries of the Python language using the two-sided independent Student’s *t*-test. Results with *p*-values ≤ 0.05 were considered significant.

### 2.10 Presence of IL ions in the active site and (*S*)-pocket cavities

To investigate the presence of ions in the active site cavity, we computed the pairwise distance of all IL heavy atoms to the center-of-mass of C_α_-atoms of residues E469/G469 and H113 of the adjacent monomer, which reliably described the center of the active site cavity throughout the simulations (**Figure 3A**). For the (*S*)-pocket in *Ap*PDC_E469G, the center-of-mass of the C_α_-atoms of residue G469 and the C7’-atom of ThDP were chosen to describe the center of the (*S*)-pocket cavity throughout the simulations (**Figure 3B**). The lowest distance of the respective molecule or molecule subunit was determined and used for analyses. A distance below 5 Å was classified as a bound state. For the cations, we additionally discriminated between interacting polar and apolar moieties of the IL cations by performing the analysis separately for the atoms of the alkyl- or PEG-chains. Only interactions accounting for at least 10 frames were considered for analysis.

**Figure 3:**
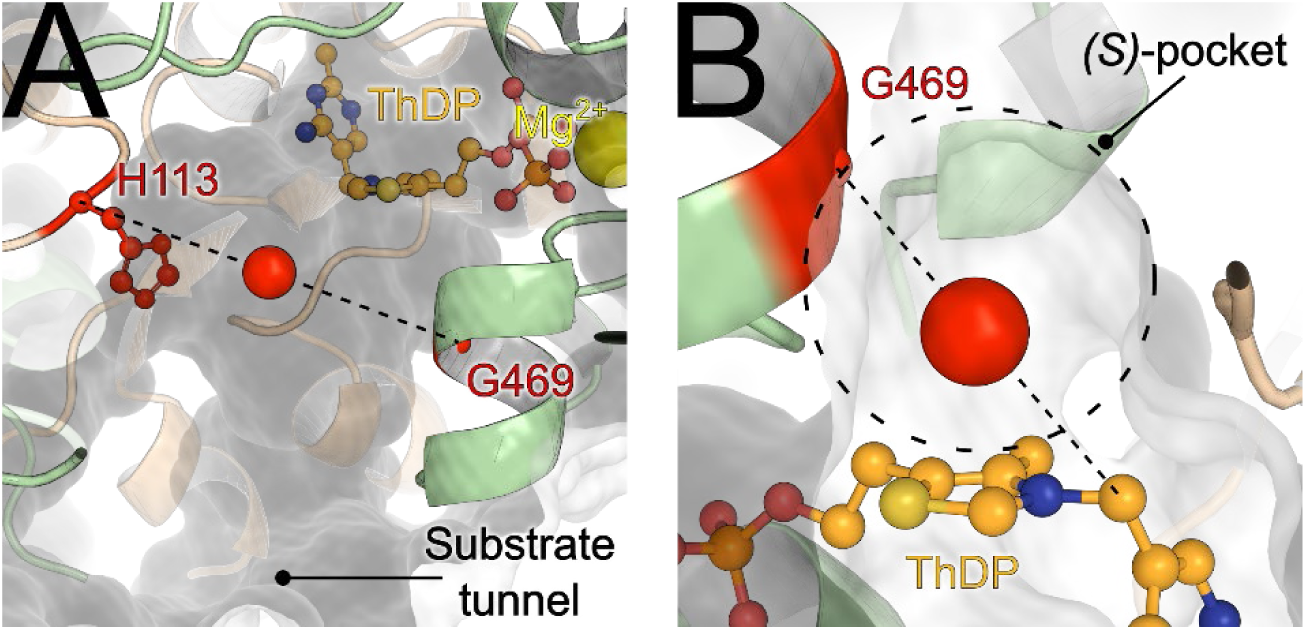
Location of the active site cavity of *Ap*PDC_E469G/*Ap*PDC_WT or the (*S*)-pocket in *Ap*PDC_E469G used in the analyses of IL presence within the active site of *Ap*PDCs. (A) The red sphere indicates the center of mass of C_α_ atoms of H113 and G469/E469, representing the center of the active site cavity throughout our simulations. (B) The red sphere indicates the center-of-mass of the C_α_ atoms of G469 and the C7’ atom of ThDP, representing the center of the (*S*)-pocket throughout our simulations of *Ap*PDC_E469G.

### 2.11 Quantitative assessment of the IL-3 cation binding on enantioselectivity

The spatial product, which describes the relative orientation of benzaldehyde to the ThDP-bound acetaldehyde, was computed via **Eq. 1**

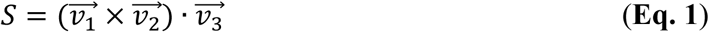

where 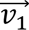, 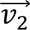, and 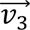 denote the vectors from the carbonyl carbon of benzaldehyde to the adjacent oxygen atom, to the C4-atom of benzaldehyde, and the C2α-carbanion/enamine [14,77] resulting from the initial binding of acetaldehyde to ThDP, respectively (see **Figure 4**). Values > 0 or < 0 indicate a substrate configuration leading to (*R*)- or (*S*)-selective product formation, respectively.

**Figure 4:**
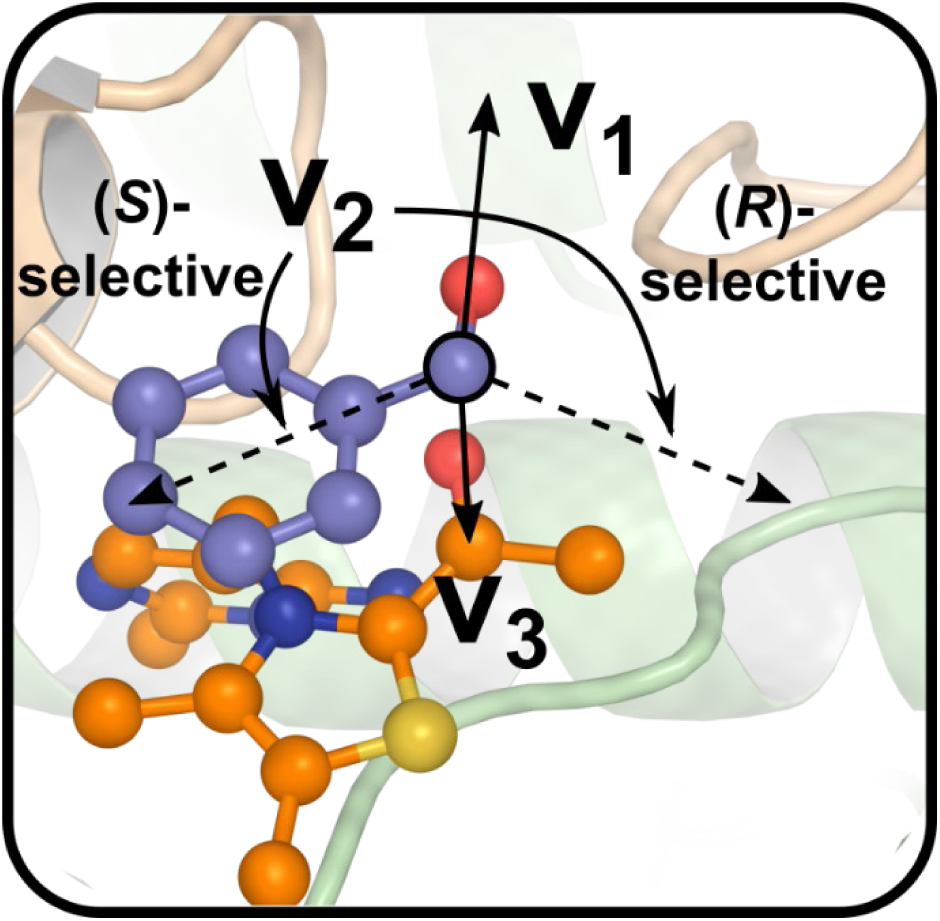
S*i*-face approach of benzaldehyde (blue) to the ThDP-bound acetaldehyde (orange) resulting in the formation of (*S*)-PAC.

### 2.12 Tunnel analyses using the CAVER 3.0 software

To test whether product egress is still possible despite the presence of the IL cation within the active site, we applied the CAVER 3.0 software [74]. Starting points for the computations were defined based on the “surrounding residues” function of CAVER. For this, the residues I468, G469, as well as the ThDP molecule were used. All residues except water were included in the tunnel computation, except for the computation of tunnel dynamics in the process of benzaldehyde vacating the active site, where BA was excluded, too. The default values were used for the probe radius (0.9 Å), shell radius (3.0 Å), shell depth (4.0 Å), and clustering threshold (3.5) for the general identification of all tunnels and visualization of the internal (water) tunnel connecting the two active sites. For visualization and computation of tunnel dynamics during BA leaving, slightly higher values for the clustering threshold (4.5), shell radius (6.0 Å), and shell depth (8.0 Å) were used. This aided in the visualization of the tunnels by the inclusion of the near-surface section of the tunnel and the grouping of almost identical tunnels. While this lengthens the computed tunnel towards the enzyme surface compared to the default values, it does not affect the other computed properties, such as the tunnel (bottleneck) radius. Computations were performed over every 10^th^ frame of the simulations for the general tunnel analyses or every frame in the case of the BA leaving trajectory and the single crystal structure, respectively. The absolute number of tunnel occurrences for the different systems was normalized to the number of frames used in the analysis.

### 2.13 Spatial distribution of ligands, substrates, and solvent molecules

To visualize the positions of ligands, substrates, and solvent molecules within the active site of *Ap*PDC and around the enzyme surface, we investigated the spatial distribution of molecules using the “grid”-functionality of the CPPTRAJ [78] program of the AMBER20 [59] simulation suite. A grid resolution of 1 Å was used in all computations. The densities were normalized according to the number of frames used in the computation with all frames being used in the computation. For the evaluation of molecules within the *Ap*PDC active sites, σ-values were chosen such as to best represent the mobility and volume of the molecules, respectively, for the different molecules ThDP (0.05), Mg^2+^ (0.02), BA (0.05), AA (0.01), or the IL-3 cation (0.05). For the analyses of solvent dynamics around the *Ap*PDC surface, the σ-values were calculated based on the number of heavy atoms within the ions. Here, a reference value of 0.0025 per atom was used and scaled accordingly, resulting in ion-specific σ-values for the IL-2 cation (78 heavy atoms; σ = 0.1950) and anion (1 heavy atom; σ = 0.0025) and the IL-3 cation (69 heavy atoms; σ = 0.1725) and anion (7 heavy atoms; σ = 0.0175). For the analyses of the spatial distribution of the IL-3 sub-structures, the core substructure was defined as the central N-atom including the two closest adjacent carbon atoms, including the carbonyl function’s oxygen, resulting in a σ-value of 0.0250 (10 heavy atoms) for the core structure and σ-values of 0.0450, 0.550, and 0.0475 for the alkyl-chain (18 heavy atoms) and the two PEG-chains (22 or 19 heavy atoms), respectively.

## 3. Results and Discussion

### 3.1 Preamble

A disadvantage of ILs is that each IL preparation may have a different composition due to impurities from the manufacturing process. Such substances may lead to misinterpretation of the observed effects. Therefore, all experiments were performed with the same preparation of ILs. In the case of the most intensively studied IL in this work, Ammoeng 102 (IL-3), the batch used was examined in detail by elemental analysis (**SI**, **Figures S7 and S8**). Small amounts of ethanol [0.02% (w/v)] and a second unidentified compound [0.06% (w/v)] were found. However, the concentrations of these impurities were very low and far below the concentrations that were shown to affect enzyme behavior in our previous study [11]. Assuming that this is true for all ILs studied, we believe it is reasonable to correlate the observed shifts with the respective ILs.

Our investigations resulted in a large data set, which is presented in full in the **SI**. In the interest of a focused and comprehensible interpretation of the effects, we limit ourselves here to the interpretation of the effects of only two enzymes. *Pf*BAL and *Ap*PDC_E469G were chosen because they represent the enzymes with the lowest and highest observed effects, respectively (**Figure 5**). The plots for the other four enzymes, as well as the plots shown below on a larger scale, can be found in the **SI**. General trends and the most pronounced effects on individual enzymes are discussed below for all biocatalysts tested.

**Figure 5:**
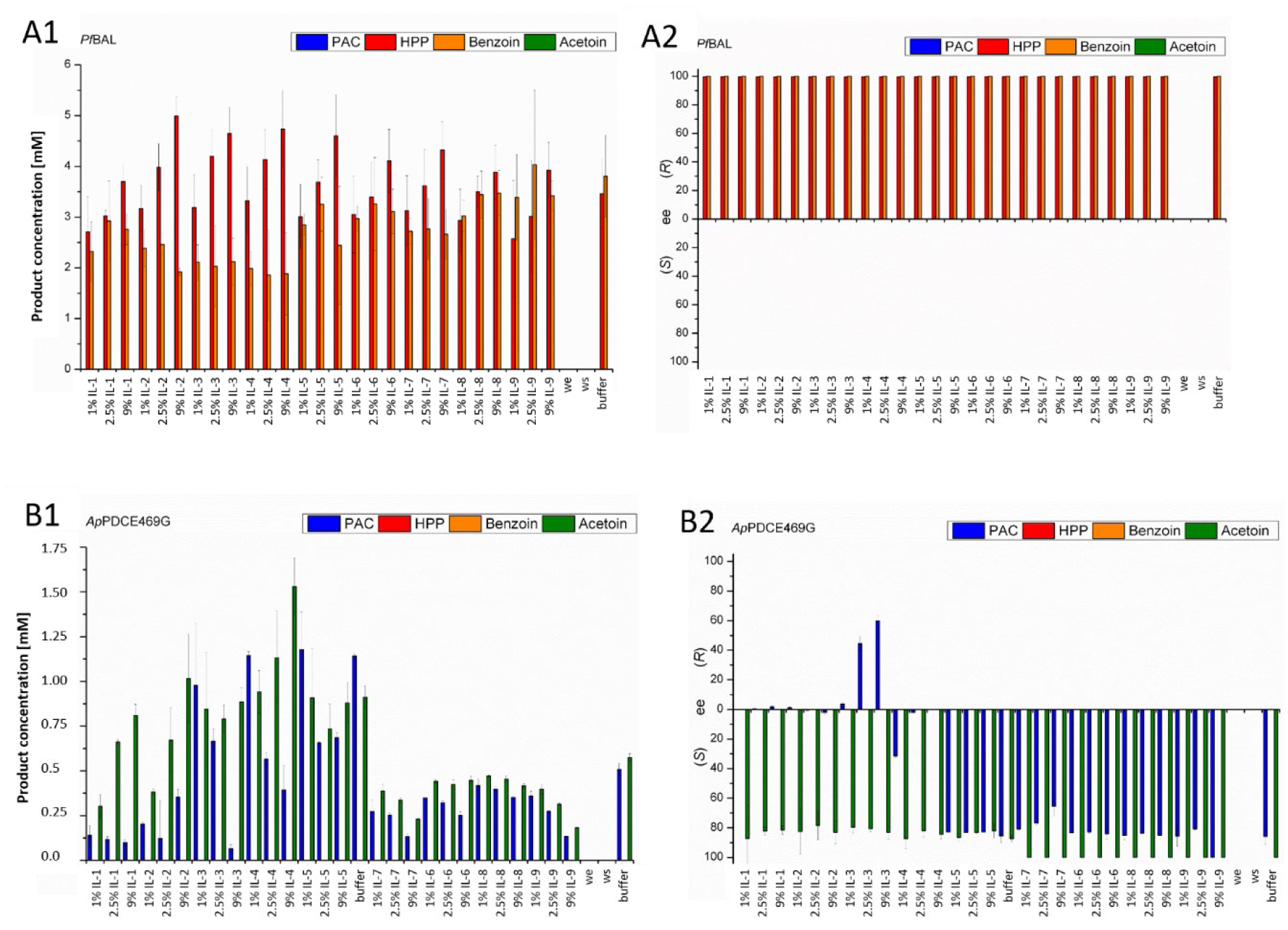
Influence of ILs on the product concentrations of the possible 2-hydroxy ketones obtained after carboligation of acetaldehyde and benzaldehyde (**A1** and **B1**) and on the enantiomeric excess of so-obtained products (**A2** and **B2**). A represents the effect on the benzaldehyde lyase from *Pseudomonas fluorescence* (*Pf*BAL) and B the effects on the pyruvate decarboxylase variant *Ap*PDC_E469G from *Acetobacter pasteurianus*. Reaction conditions for **A**: TEA-buffer (50 mM, 2.5 mM MgSO_4_, 0.1 mM ThDP), pH 8.0, 0.02 mg/mL purified, lyophilized *Pf*BAL, 18 mM acetaldehyde, 18 mM benzaldehyde. Reaction conditions for **B**: same as for **A**, but the buffer pH was 7.5 and the enzyme concentration with 0.1 mg/mL purified, lyophilized *Ap*PDC_E469G higher due to generally lower activity (if the conversion was still too low for product determination 0.4 mg/mL *Ap*PDC_E469G was added). All bars represent the arithmetic mean of three independent replicates. Since the error bars are difficult to see in this comprehensive overview, all plots can be found in the **SI** on a larger scale (**Figures S1-S6**). we = same experiment but without enzyme, ws = same experiment but without substrate; buffer = same experiment but without addition of IL.

### 3.1. Chemoselectivity

#### 3.1.1 Putative influence of ILs on the volume of the active site

We have observed significant effects of ILs on the chemoselectivity of enzymes. In most cases, there is a clear shift from the larger to the smaller 2-hydroxy ketone in the presence of ILs. For example, for *Pf*BAL, which catalyzes the formation of 2-HPP and benzoin in buffered systems, a clear shift toward increased formation of the smaller product 2-HPP with a concomitant decrease in the production of the larger product benzoin can be observed in the presence of ILs (except for a low concentration of IL-9, **Figure 5A1**). For the *Ap*PDC_E469G variant, a shift towards the smaller product, in this case from PAC to acetoin, is particularly pronounced in the presence of Ammoeng ILs. The only exception was *Pp*BFDH281A, where we observed higher concentrations of all products, including the sterically demanding ones, in the presence of 2.5% (w/v) of almost all ILs tested (**SI**, **Figure S3**).

We also found similar trends in the presence of organic solvents and in buffer without additives when the biotransformations were performed at elevated temperature [11]. As temperature increases mobility, a less spacious active site could be a factor for smaller ligation products. The same trend towards smaller ligation products is expected if the ILs, especially the often large cations, can also directly interfere with amino acids in the active site of ThDP-dependent enzymes. Some of the sterically demanding ions are hydrophobic. This allows them to interact with hydrophobic residues located in parts of the active site, such as residues T384, G385, I468, and G469 of the (*S*)-pocket (identified in MD simulations of the *Ap*PDC_E469G variant), which are located in the acceptor binding region. Such preferential hydrophobic interactions have been suggested for several small and particularly hydrophobic organic solvents for the enzymes described here, but have also been observed, for example, in tyrosinase from *Bacillus megaterium*, where a direct interaction of sodium dodecyl sulfate (SDS) with residues at the entrance of the active site was visualized in the crystal structure [79]. The possible direct interaction of the IL cation is discussed in more detail below in relation to the observed shifts in enantioselectivity.

### 3.2 Enantioselectivity

The observed effects on enantioselectivity were highly enzyme-dependent. From almost no effect, as with *Pf*BAL (**Figure 5A2**), to a reduction or even inversion of enantioselectivity, as with *Ap*PDC_E469G (**Figure 5B2**), to improved selectivity, all effects were observed as described below. Since the most pronounced effects were observed with the *Ap*PDC_E469G variant, this enzyme was selected for more detailed studies.

#### 3.2.1 Improvement of poor or moderate enantioselectivity

In several cases, the addition of ILs improved poor or moderate stereoselectivities. For carboligations towards acetoin using ThDP-dependent enzymes, enantioselectivity is predominantly poor due to the small size of the acetaldehyde, which prevents a preferred parallel or antiparallel orientation prior to carboligation [2,80]. For example, in the case of *Ll*KdcA, the enantioselectivity of acetoin formation [*ee* = 24% (*R*) in buffer] was improved up to 40% *ee* (*R*) in the presence of ILs such as Ammoeng 112 [IL-4, 9% (w/v)] (**SI**, **Figure S1C**). None of the organic solvents resulted in such a pronounced improvement in the selectivity of *Ll*KdcA [11]. Similarly, in the case of *Pp*BFD-catalyzed acetoin formation, the addition of ILs such as 2.5% (w/v) Ammoeng 101 (IL-2) increased the *ee* values from 16.8 to 23% (*R*) in buffer to 42% (*R*) (**SI**, **Figure S2C**). The same trend was observed with the *Pp*BFD_H281A variant, where a moderately (*R*)-selective synthesis of acetoin in buffer [*ee* = 34.4% (*R*)] was improved to 76% *ee* (*R*) by the addition of 2.5% (w/v) Ammoeng 100 (IL-1) (SI, **Figure S3C**). This is the highest (*R*)-selectivity for acetoin formation achieved so far by the six ThDP-dependent enzymes tested. For *Pp*BFD_H281A, the selectivity for (*S*)-2-HPP was slightly improved from 55% (*S*) to 62% *ee* (*S*) in the presence of 9% Ecoeng 110 (IL-5), in contrast to IL-2, which reduced the ee to 10% (*S*), and IL-1, which caused the formation of a small excess of (*R*)-2-HPP (*ee* = 18%) (**SI**, **Figure S3C**).

In the case of *Ap*PDC_WT, the presence of 9% (w/v) Ammoeng 100 (IL-1) increased the selectivity for (*S*)-acetoin from 30% (*S*) to 36% *ee* (*S*) (**SI**, **Figure S5C**). The IL-dependent increase in (*S*)-selectivity was generally lower than for the (*R*)-selective carboligations, consistent with similar effects observed with organic co-solvents [11]. This was interpreted as an occupation of the (*S*)-pocket by organic solvents of appropriate size and hydrophobicity. The interaction of the often bulky cations and anions of ILs with the active sites of ThDP-dependent enzymes was studied in detail experimentally and by MD simulation (see below).

#### 3.2.2 Decrease of enantioselectivity

Despite the positive effect of some ILs (e.g. Ammoeng 112, IL-4) on the enantioselectivity, especially the (*R*)-selective acetoin formation, other ammonium ILs (e.g. IL-8 with *Ll*KdcA) drastically reduced the enantioselectivity. For *Ll*KdcA, the selectivity shifts introduced by addition of ILs were much less pronounced than for organic cosolvents [11]. In contrast, in *Ap*PDC_E469G-catalyzed reactions, the selectivity for PAC production was shifted from the preferred formation of the (*S*)-product in buffer (87% *ee*) to the predominant formation of the (*R*)-product (60% *ee*) in the presence of 9% (w/v) Ammoeng 102 (IL-3) (**Figure 5B2** & **SI**, **Figure S6C**).

In the case of *Pp*BFD_H281A-catalyzed biotransformations, pronounced influences were found with respect to the products 2-HPP [in buffer *ee* = 47% (*S*)] and acetoin [in buffer *ee* = 34% (*R*), SI, **Figure S3C**]. For example, for acetoin, the selectivity decreased to 17% *ee* (*R*) in the presence of Ammoeng 100 [9% (w/v), IL-1] and to 12% *ee* (*R*) with Ecoeng 110 [9% (w/v), IL-5]. For *Pp*BFD-catalyzed reactions, a decrease in (*R*)-selective acetoin production [in buffer *ee* = 16.8 - 23% (*R*)] was observed. The presence of 1% (w/v) Ammoeng 112 (IL-4) decreased the *ee* to a value of *ee* = 6% (*R*) (**SI**, **Figure S2C**).

#### 3.2.4 No influence on highly enantioselective biotransformations in buffer > 99% *ee*

As already observed in our related studies with organic cosolvents [11], reactions that are highly stereoselective in buffer remain so upon addition of ILs (probably due to an optimal fit of both reaction partners in only one arrangement in the active site of the respective enzyme). This is true for the *Ll*KdcA-catalyzed formation of (*R*)-PAC and (*R*)-2-HPP as well as for the stereoselective formation of (*R*)-benzoin and (*R*)-2-HPP by *Pf*BAL (**Figure 5A2** & **SI**, **Figure S1**).

Besides the achiral ILs (IL-1 to IL-9), we also tested several imidazolium-based chiral ILs (IL-10 to IL-14) (**Figure 2**) for effects on the enantioselectivity of the six ThDP-dependent enzymes. In all cases, however, no significant effect on enantioselectivity was observed and the effect of these additives remained far behind the effect of organic solvents and the tested achiral ILs.

#### 3.2.5 No significant improvement in (*S*)-selectivity

As previously reported [11], the use of organic cosolvents in aqueous buffer did not significantly improve the initial (*S*)-selectivity observed in ThDP-dependent enzymes. We hypothesized, especially for the (*S*)-selective variant *Ap*PDC_E469G, that organic solvents of appropriate size selectively block the (*S*)-pocket, thereby preventing the antiparallel arrangement of the two aldehydes prior to carboligation. Also, in the presence of ILs, there was no significant improvement in the (*S*)-selectivity observed in buffer (besides the minor enhancement of (*S*)-acetoin formation with *Ap*PDC described above). We assumed that the proposed selective blockade of the (*S*)-pocket would also be applicable to the system discussed here. To validate this, the enantioselectivity of PAC formation catalyzed by *Ap*PDC_E469G was investigated in more detail. It was found that only ammonium-based Ammoeng ILs affected the selectivity of *Ap*PDC_E469G-catalyzed reactions (**Figure 6**).

**Figure 6:**
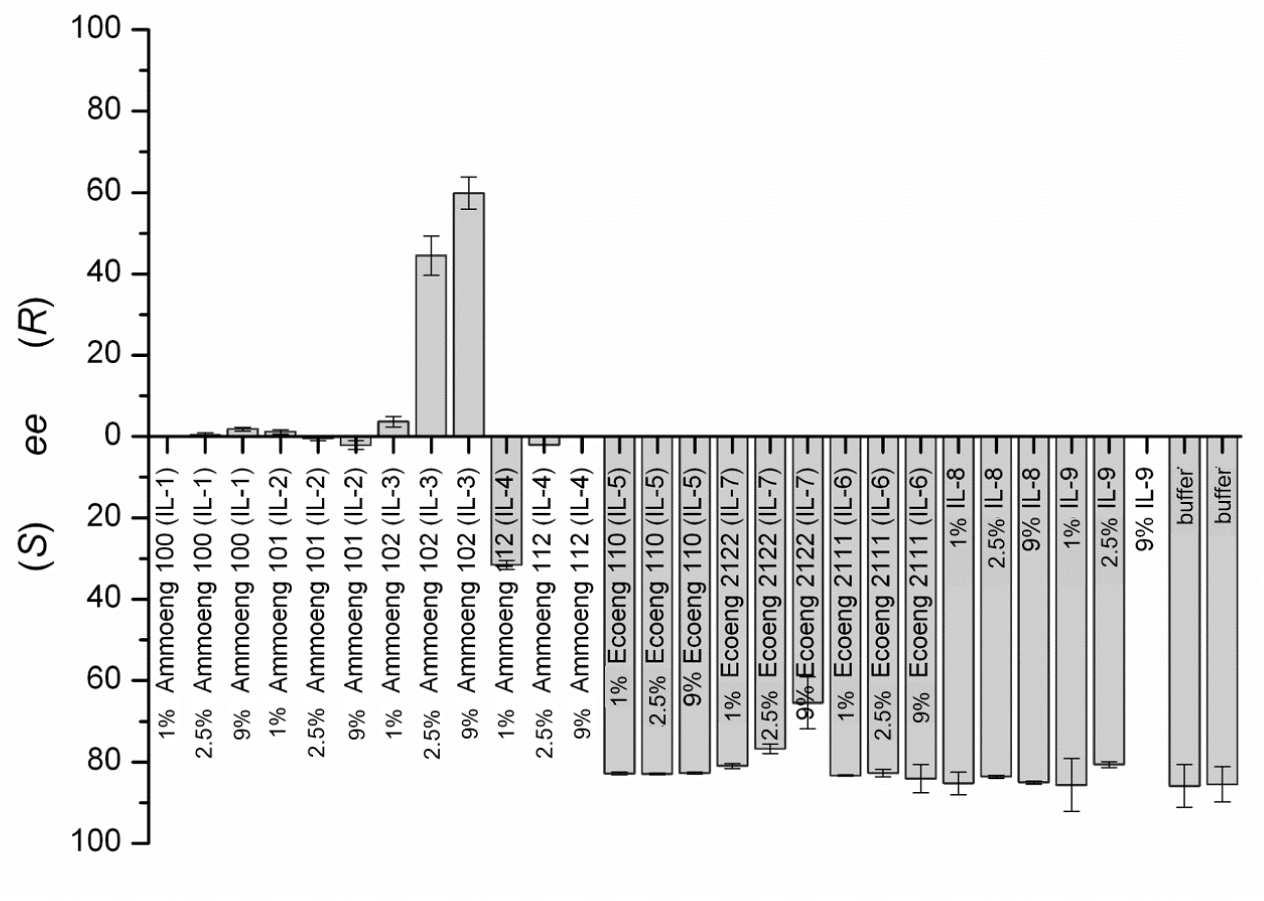
Enantioselectivity for the production of PAC in *Ap*PDC_E469G-catalyzed biotransformations in the presence of IL-1 to IL-9. Reaction conditions: triethanolamine (TEA) buffer (50 mM, 2.5 mM MgSO_4_, 0.1 mM ThDP), pH 7.5, 20°C, 24 h, 0.1 mg/mL *Ap*PDC_E469G, substrates: 18 mM acetaldehyde, 18 mM benzaldehyde. All presented values are mean values from three independent measurements.

In the pronounced case, the enantioselectivity of *Ap*PDC_E469G was shifted from an *ee* of 86% (*S*) to an *ee* of 60% (*R*) in the presence of 9% (w/v) Ammoeng 102 (IL-3). The shift was dependent on the concentration of Ammoeng 102 (IL-3) (**Table 1**). In addition to shifts in enantioselectivity, we also observed a reduction in overall activity. **Table 1** shows the influence of Ammoeng 102 (IL-3) to clarify whether both enantiomers are formed at lower concentrations or whether the concentration of one enantiomer is selectively reduced.

**Table 1.** *Ap*PDC_E469G catalyzed formation of PAC in the presence of different concentrations of Ammoeng 102 (IL-3). Concentrations and *ee* values after 24 h reaction time are shown. (For experimental information see **Figure 4**.)

|  |  |  |  |  |  |  |
| --- | --- | --- | --- | --- | --- | --- |
| **Ammoeng 102 (IL-3)** | **PAC** | ***ee*** | **(*R*)-PAC** |  | **(*S*)-PAC** |  |
| (%, w/v) | mM | % | % | mM | % | mM |
| 0 | 1.1 ($\pm 0.01$) | 86 ($\pm 4.3$) (*S*) | 7 | 0.1 | 93 | 1.0 |
| 1 | 1.0 ($\pm 0.4$) | 4 ($\pm 1.0$) (*R*) | 52 | 0.52 | 48 | 0.48 |
| 2.5 | 0.7 ($\pm 0.1$) | 45 ($\pm 5.0$) (*R*) | 72.5 | 0.51 | 27.5 | 0.19 |
| 9 | 0.07($\pm 0.0$) | 60 ($\pm 4.0$) (*R*) | 80 | 0.06 | 20 | 0.01 |

Although the total concentration of PAC was reduced by 30% by the addition of 2.5% (w/v) Ammoeng 102 (IL-3) compared to the addition of 1% (w/v) Ammoeng 102 (IL-3), the total concentration of the (*R*)-enantiomer remained unchanged (0.52 to 0.51 mM). Thus, in this concentration range, the loss of activity correlates with a selective decrease in the production of the (*S*)-enantiomer and not the (*R*)-enantiomer. The previously published hypothesis that organic solvents of appropriate size cause a selective blockade of the (*S*)-pocket could therefore apply to ionic liquids.

Since the (*S*)-pocket was designed for the side chain of benzaldehyde [7], it was not initially expected that the significantly larger Ammoeng cations could reach the (*S*)-pocket through the substrate channel in the enzyme. Therefore, the effect of the much smaller anions was investigated first. Since benzaldehyde with the aromatic ring can be accommodated in the (*S*)-pocket [11], all anions with a solvent-excluded volume < 94.6 Å^3^ should be able to reach this pocket, at least, if only their volume is considered.

**Table 2.** Solvent-excluded volume (calculated by ChemDraw (PerkinElmer) for Excel Add-In) of anions of ammonium-based ionic liquids in comparison to benzaldehyde as substrate.

|  |  |
| --- | --- |
| Solvent-excluded volume ($\text{\AA}^3$) | |
| Chloride | 29.3 |
| Methyl sulfate | 67.7 |
| Ethyl sulfate | 71.6 |
| Dihydrogen phosphate | 92.6 |
| Benzaldehyde | 94.6 |

To investigate whether the anions alone affect the enantioselectivity of *Ap*PDC_E469G-catalyzed PAC formation, the five anions of the different Ammoeng ILs were added to the respective biotransformations in combination with the same cation (sodium) (**Figure 7**). In addition, an ammonium cation was added to the reaction as ammonium sulfate to investigate whether the type of cation influences the enantioselectivity (**Figure 7**).

**Figure 7:**
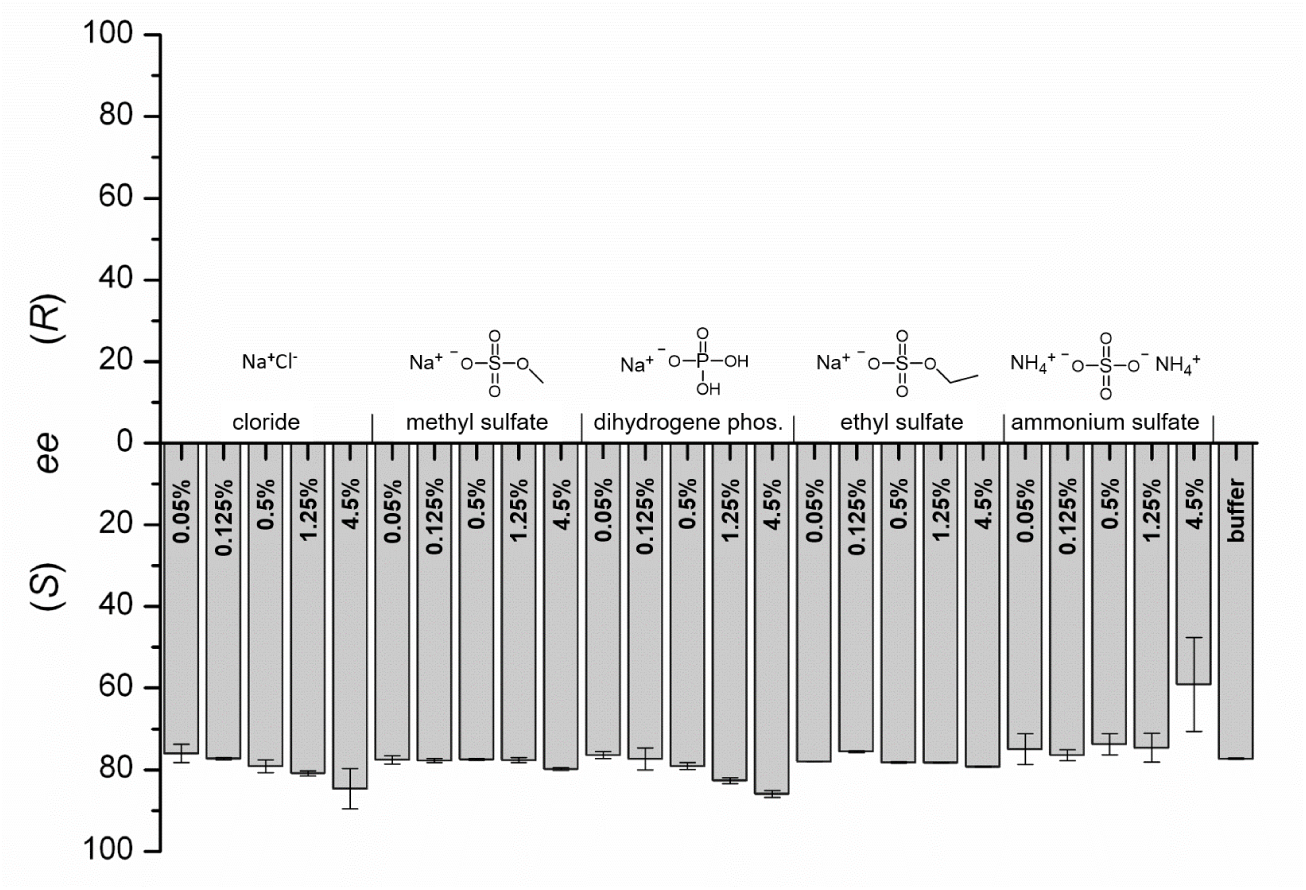
Enantioselectivity of *Ap*PDC_E469G-catalyzed PAC formation in the presence of different sodium salts. Reaction conditions: TEA-buffer (50 mM, 2.5 mM MgSO_4_, 0.1 mM ThDP), pH 7.5, 20°C, 24 h, 0.1 mg/mL *Ap*PDC_E469G; substrates: 18 mM acetaldehyde, 18 mM benzaldehyde. The concentration of the salt is given within the bar (this concentration resembles the molar concentration of the corresponding Ammoeng ionic liquid, compare **SI**). Shown values are mean values from two independent measurements.

When the different sodium salts were added to the *Ap*PDC_E469G-catalyzed carboligation, there was almost no change in the enantioselectivity of PAC production (**Figure 7**). Obviously, the anions are not responsible for the enantioselectivity shift observed with ILs. While contradicting early beliefs that anions are the dominant species responsible for IL effects [28], our observation is consistent with more recent observations that ion-specific effects must be considered [81] and that IL effects often arise from a few specific and typically favorable IL-enzyme interactions [27,30]. In our case, these potential ion-specific effects or favorable IL-enzyme interactions are likely to be strongly influenced by the physicochemical properties of the substrate tunnel, the active site, and specifically the (*S*)-pocket, which are designed to guide and accommodate the neutral substrate benzaldehyde, as in the case of *Ap*PDC_E469G (**Figure 6** and **Figure 7**).

#### 3.2.6 Binding of the Ammoeng 102 cation via hydrophobic interactions to the (*S*)-pocket shifts enantioselectivity in *Ap*PDC_E469G but not in *Ap*PDC_WT

To elucidate the molecular mechanism of IL-induced enantioselectivity shifts, we investigated the influence of 0.08 M Ammoeng 102 (IL-3) versus pure water on the structural dynamics of *Ap*PDC_WT and *Ap*PDC_E469G using all-atom molecular dynamics (MD) simulations of in a total length of 320 µs. Ammoeng 102 was chosen because of its strongest effect on the enantioselectivity of *Ap*PDC_E469G. For *Ap*PDC_E469G, we additionally performed MD simulations of a total of 96 µs length of the structurally similar Ammoeng 101 (IL-2) at the same concentration to identify effects on enantioselectivity due to different IL structures; IL-2, similar to the other ILs of the Ammoeng series except IL-3, leads only to a racemic mixture of PAC. Both ILs were found to interact frequently and at different positions with surface residues of *Ap*PDC_WT and *Ap*PDC_E469G (see **Figure S17** and **Text S6** for analyses of the spatial ion distribution around *Ap*PDC). However, the global enzyme structures of *Ap*PDC_WT and *Ap*PDC_E469G remained invariant in both water and ILs over the 8 µs timescale of the MD simulations (see **Figure S11** and **Text S1**), indicating that these interactions with *Ap*PDC surface residues and the (overall unaffected) global structural changes are not responsible for the shift in enantioselectivity (see **Figure 7**).

Instead, we observed that both cations and anions of Ammoeng IL-2 and IL-3 entered the active sites of *Ap*PDC_WT and *Ap*PDC_E469G via the substrate tunnel (**Figure 8A**; see SI for alternative entry routes). A detailed description of the frequencies of these visits for the different IL-ions is given in **Text S2**. As an example, **Figure 8B** shows the evolution of the distance to the center of the active site cavity for selected ions over the simulation time with the insets showing representative binding modes observed throughout the simulations. Both the IL-2 and IL-3 cations interacted with their hydrophobic and hydrophilic moieties. However, the hydrophobic interactions of IL-2 required a sterically hindered and potentially unfavorable deeper penetration of the cation core into the substrate tunnel to reach the active site due to the shorter alkyl chain length (see **Figure 8B**), whereas this is not required for the interactions of IL-3, suggesting a possible role of this structural feature in the lesser effect on enantioselectivity for IL-2. Notably, our MD simulation setup is sensitive enough to probe for such differences between the tested ILs. For anions, we typically observed much shorter visits to the active site cavity (median of 508 ns and 115 ns for ethyl sulfate and chloride, respectively) compared to cations (median of 1134 ns and 936 ns for the IL-3 and IL-2 cation, respectively), often followed by a complete exit from the active site (see **Text S2** in the **SI**).

**Figure 8:**
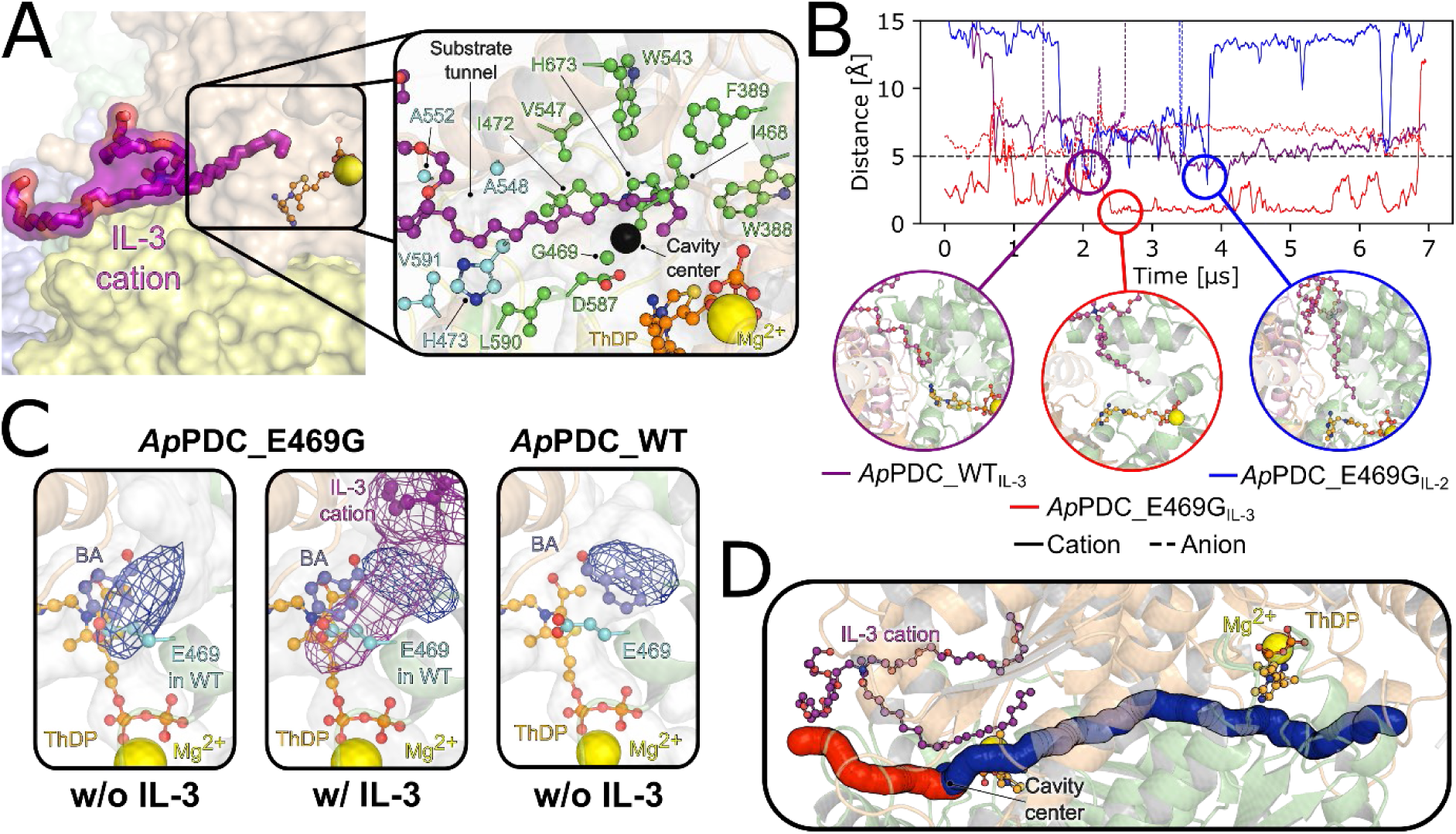
IL ions alter the substrate binding modes by direct interactions within the active site of *Ap*PDC. (A) Binding mode of the IL-3 cation (purple). The ion surface is shown for solvent-exposed atoms, i.e., a lack of surface indicates atoms within the *Ap*PDC substrate tunnel. Structures of ThDP and Mg^2+^ are shown to indicate the location of the catalytic site. The inset shows a close-up view of residues of the tunnel entrance (cyan) or active site residues (green) interacting with the IL-3 cation. The approximate center-of-mass of the active site cavity, used for subsequent distance calculations in panel B, is indicated by a black sphere. (B) Distances over time of selected IL cations (solid lines) and anions (dashed lines) to the active site cavity for *Ap*PDC_WT in IL-3 (purple), *Ap*PDC_E469G in IL-3 (red), and *Ap*PDC_E469G in IL-2 (blue) with close-up views of representative binding modes. (C) Spatial distribution analyses of solvent and substrate molecules. The ball and stick representations of ThDP, Mg^2+^, benzaldehyde (BA), and the IL-3 cation show the starting positions, while regions with a high density of BA or IL-3 cations throughout the MD simulations are shown as blue or purple meshes, respectively. The position of the (*S*)-pocket in *Ap*PDC_E469G is indicated by cyan sticks of the *Ap*PDC_WT residue aligned to the *Ap*PDC_E469G structure. All distributions were normalized according to the number of frames. *σ*-values defining the intensity cutoff of the represented data of 0.05 for BA and IL-3 cations were used. (D) Accessibility of the active site for *Ap*PDC_E469G when an IL-3 cation is already present in the substrate tunnel. The average and maximum tunnel diameters of alternative access tunnels to the active site are statistically indifferent or larger than the native substrate tunnel in the crystal structure or in simulations of *Ap*PDC in water, allowing a substrate or product to pass through [Table 1 in Ref. 11]. An alternative substrate tunnel (red) and the tunnel connecting the two active sites of the dimer of *Ap*PDC_E469G (blue) are shown as examples.

In summary, these results indicate that IL cations are predominantly responsible for the observed shifts in enantioselectivity by influencing the binding and dynamics of substrates through direct interactions. This is consistent with the experiments (**Figures 5 and 6**), and suggests a mechanism similar to that proposed for organic solvents [11]. If these observations and the IL structures are combined with the experiments, these results also suggest that (i) the hydrophilic PEG chain may be mainly involved in the formation of the racemic mixture of PAC in the carboligation using *Ap*PDC_E469G in the presence of IL-2, as this is a common structural feature of all ILs of the Ammoeng series, while (ii) the extended alkyl chain of the IL-3 cation may play a crucial role in the inversion of enantioselectivity in the presence of IL-3.

Based on the above results, we investigated how the presence of the IL-3 cation alkyl chain in the active site affects the substrate binding of *Ap*PDC. Therefore, we set up additional MD simulations of in total 128 µs length including the substrates benzaldehyde (BA) and acetaldehyde (AA) in configurations positioned according to the respective binding modes in *Ap*PDC_E469G or *Ap*PDC_WT resulting in (*S*)- or (*R*)-configurated products [14,82]. We simulated these systems with and without an IL-3 cation alkyl chain placed in the substrate tunnels of both *Ap*PDC variants. In *Ap*PDC_E469G, the IL-3 cation adopted a binding mode in which the alkyl chain occupies the (*S*)-pocket and BA is displaced toward the active site cavity responsible for (*R*)-selective product formation, whereas benzaldehyde typically remained in the configuration leading to the (*S*)-product in the absence of the IL-3 cation (see **Figure 8C**). Our results also showed that this binding mode is persistent, as the IL-3 cation remained bound in all simulations of the second set of simulations starting from this position. However, the large active site cavity of *Ap*PDC_E469G allows the IL-3 cation to sample a large configurational space, i.e., different locations and conformations, such that only one out of twelve replicas assumed this state within 750 ns in the 3^rd^ set of simulations. Taken together, these results indicate that, although the binding of the IL-3 cation is a rare event, the binding mode is persistent over longer durations when it occurs. In *Ap*PDC_WT, the presence of the IL-3 cation did not have a clear effect on the configuration of benzaldehyde leading to the (*R*)-product, because benzaldehyde and the IL-3 cation mostly competed for the available space in the active site cavity. Taken together, these observations suggest a potential mechanism for a enantio-selectivity shift in *Ap*PDC_E469G in which the extended alkyl chain of an IL-3 cation, but not the alkyl chain of an IL-2 cation, effectively displaces benzaldehyde from the (*S*)-pocket due to a likely stronger binding of the IL-3 cation to the (*S*)-pocket compared to benzaldehyde.

To establish a correlation between the simulation and the experimental data, we simulated the approach of benzaldehyde to ThDP using controlled MD simulations for both *Ap*PDC variants with and without an IL-3 cation, such that either the *Re*-face attack in the (*S*)-pocket (*Ap*PDC_E469G) or the *Si-*face attack (*Ap*PDC_WT) was blocked, and evaluated the effect on enantioselectivity by computing the spatial product of vectors describing the relative orientation of benzaldehyde to ThDP using **Eq. 1**. Our results are in qualitative agreement with experimentally determined effects on enantioselectivity in that *Ap*PDC_WT did not experience shifts in enantioselectivity in the presence of the IL-3 cation, whereas the presence of the cation reversed the enantioselectivity in *Ap*PDC_E469G (**Table 3**), as reflected by the large and statistically significant change in the spatial product from (*S*)-PAC to (*R*)-PAC in the latter case only.

**Table 3.** Spatial product of vectors describing the benzaldehyde orientation relative to acetaldehyde-bound ThDP. Values are shown as mean ± standard error of the mean (*n* = 100); values > 0 or < 0 indicate an configuration leading to the (*R*)- or (*S*)-selective product, respectively. The *p*-value is determined via a two-sided Student’s *t*-test. In addition, the excess stereoisomer found in experiment is given (see also **Figures 5 and 6**).

| | without IL-3 cation | stereoisomer in excess (experiment) | with IL-3 cation | stereoisomer in excess (experiment) | $p$-value |
| --- | --- | --- | --- | --- | --- |
| *ApPDC_WT* | $1.79 \pm 0.54$ | (*R*) | $1.10 \pm 0.53$ | (*R*) | 0.36 |
| *ApPDC_E469G* | $-1.47 \pm 0.43$ | (*S*) | $0.90 \pm 0.46$ | (*R*) | $< 0.01$ |
| $p$-value | $< 0.01$ | | 0.77 | | 0.21 |

To test whether substrate access and product exit are possible even in the presence of an IL-3 cation in the active site, i.e., when the main substrate tunnel is blocked, we analyzed the accessibility of the active site of all individual MD snapshots using the CAVER software [74]. Alternative tunnels originating from the active site cavity following different routes to the surface than the main (blocked) substrate tunnel were identified in almost half of all MD frames (see **Figure 8D**). The average and maximum bottleneck radii calculated across the different replicas, where the bottleneck radius describes the narrowest point of a given tunnel, were not statistically different in *Ap*PDC_E469G with bound IL-3 cation (*Ap*PDC_E469G_+IL_) from those observed in MD simulations of *Ap*PDC_WT in water or *Ap*PDC_E469G without IL-3 cation (*Ap*PDCE469G_-IL_), as well as the *Ap*PDC_WT crystal structure (**Table 4**). Note that the formation of alternative tunnels did not require the binding of an IL-3 cation, but can also occur spontaneously, although a promoting role of IL-ions cannot be excluded (see **SI**, **Figure S13**). These results indicate that even when the main substrate tunnel is blocked, substrate egress is possible via the formation of (probably temporary) alternative substrate tunnels.

**Table 4.**
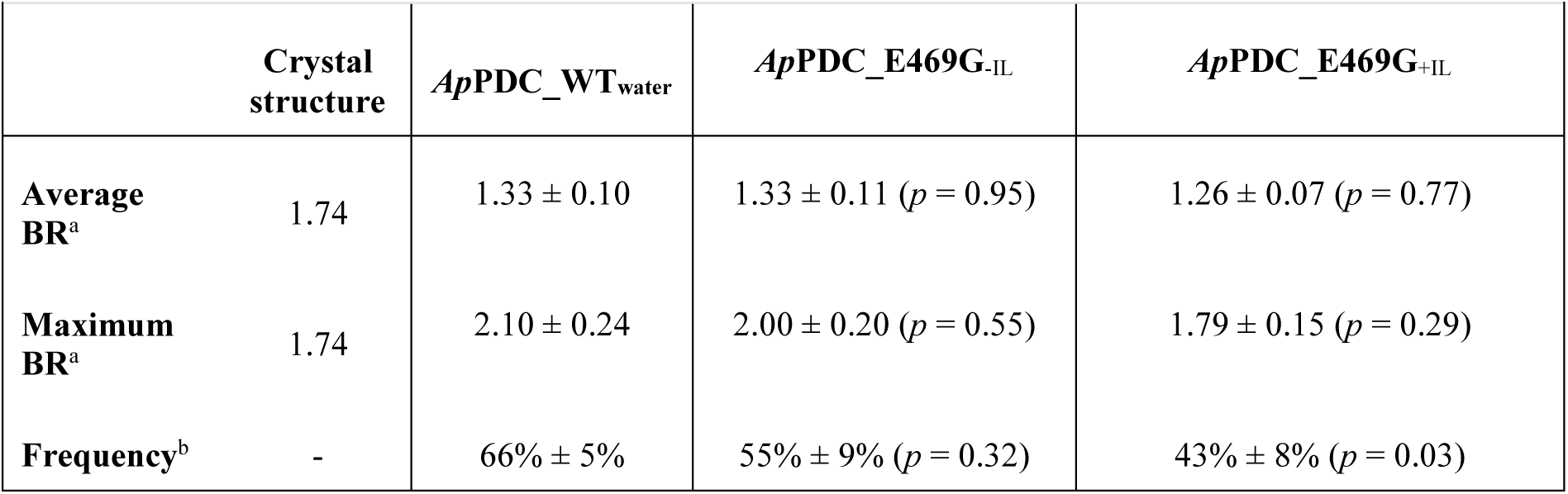
Structural parameters of highest-priority tunnels in *Ap*PDC. Values are shown as mean ± standard error of the mean (*n* = 8-12). *p*-value to *Ap*PDC_WT in water (determined via a two-sided Student’s *t*-test) is shown in brackets. ^a^ Bottleneck radius (BR) in Å. ^b^ In %.

## 4. Conclusion & Outlook

This study demonstrates the profound influence of the addition of ILs on the activity, chemo- and enantioselectivity of ThDP-dependent carboligations. Regarding the molecular effects of ILs on the chemo- and enantioselectivity of ThDP-dependent enzymes, the effects already discussed for enzymatic carboligations in the presence of organic cosolvents [11] seem to apply also in the case of IL addition.

While decreases in activity and changes in chemo- and enantioselectivity were observed when several ThDP-dependent enzymes were incubated with ILs as additives in aqueous reaction media, improvements were also observed in all three categories. For activity, a more than 2.5-fold increase in activity for the formation of PAC and acetoin was observed when *Ap*PDC_E469G was used in the presence of 9% (w/v) Ammoeng 112. In terms of chemoselectivity, the addition of 1% to 9% (w/v) Ammoeng 100 resulted in improved chemoselectivity for several enzymes, with the most notable improvement observed for *Pp*BFD. Finally, the most pronounced enantioselectivity shifts were observed with *Ap*PDC_E469G, although the most substantial improvements in terms of enantiomeric excess were achieved with *Pp*BFD_H281A for the synthesis of (*R*)-acetoin with improvements in *ee* from 34% (*R*) up to 76%.

The results of MD simulations indicated that the extended hydrophobic and hydrophilic moieties of the cations of Ammoeng 102 and Ammoeng 101 could enter the active site of *Ap*PDCs via the substrate tunnel. This provides a plausible mechanism for the IL-induced enantioselectivity shifts in carboligation, as IL cation binding affected benzaldehyde binding to the (*S*)-pocket in *Ap*PDC_E469G, but not in *Ap*PDC_WT, which lacks such a pocket. Additional simulations and simulated annealing experiments further emphasized the pivotal role of the composition of the interacting cation moiety in this process. The Ammoeng 102 cation was observed to induce a complete inversion of enantioselectivity by establishing preferential hydrophobic interactions with the (*S*)-pocket of *Ap*PDC_E469G. By contrast, such hydrophobic interactions of the alkyl chain were found to be sterically hindered in the case of the Ammoeng 101 cation. This indicates that the hydrophilic PEG chains of both cations (present in all cations of the Ammoeng series) play a secondary role in benzaldehyde binding to the (*S*)-pocket, resulting in the formation of racemic PAC in the carboligation employing *Ap*PDC_E469G with Ammoeng 101.

It can be concluded that solvent engineering is an additional tool for modifying enzyme selectivity and, consequently, for engineering the product range of biotransformations.

## Abbreviations

2-HPP: 2-Hydroxypropiophenone, 2-Hydroxy-1-phenyl-1-propanone;
PAC: phenylacetylcarbinol, 1-Hydroxy-1-phenyl-2-propanone

## Author Contributions

TG: Methodology, Investigation, Validation, Formal analysis, Writing-original draft; THE: Methodology, Investigation, Validation, Formal analysis, Visualization; Writing-original draft; ASK: Investigation, Validation, Writing-review&editing; EvL: Visualization, Writing-review&editing; CEP: Resources; Writing-review&editing; IL: Ressources; VG-F: Writing-review&editing; JP: Writing-review&editing; MP: Conceptualization; Writing-review&editing; HG: Conceptualization, Ressources; Writing-review&editing, Project administration; Funding application; DR: Dörte Rother: Conceptualization, Ressources; Writing-review&editing; Project administration; Funding application

## Supplementary Data

Detailed plots visualizing the influence of IL addition on activity, chemo- and enantioselectivity for the combination of all tested enzymes and all ILs including well-visible error bars as well as further details of the MD simulations can be found online at XXX.

## Acknowledgments

Tina Gerhards thanks the German Research Foundation DFG for financial support within the Research Training Group 1166 “BioNoCo”, and Carolin Paul thanks the EU FP7 “BIOTRAINS” grant number 238531 for funding. We thank the Central Institute for Analytics (ZEA-3) at Forschungszentrum Jülich GmbH for purity analysis of Ammoeng 102. We are grateful for computational support and infrastructure provided by the “Zentrum für Informations-und Medientechnologie” (ZIM) at the Heinrich Heine University Düsseldorf and the computing time provided by the John von Neumann Institute for Computing (NIC) on the supercomputer JUWELS at Jülich Supercomputing Centre (JSC) (user ID: VSK33, protil).

## Data and Software Availability Statement

The data underlying this study are available in the published article and its Supporting Information. The molecular simulations data are available here: https://researchdata.hhu.de/handle/entry/187 at https://researchdata.hhu.de.

The AMBER suite of biomolecular simulation programs is available here: https://ambermd.org/.

## References

[1] Hailes HC, Rother D, Müller M, Westphal R, Ward JM, Pleiss J, et al. Engineering stereoselectivity of ThDP-dependent enzymes. FEBS J 2013;280:6374–94. 10.1111/febs.12496.

[2] Pohl M, Dresen C, Beigi M, Müller M. Enzymatic Acyloin and Benzoin Condensations. In: Drauz K, Gröger H, May O, editors. Enzym. Catal. Org. Synth., vol. II, Weinheim: Wiley-VCH; 2012, p. 919–46. 10.1002/9783527639861.ch22.

[3] Dünkelmann P, Kolter-Jung D, Nitsche A, Demir AS, Siegert P, Lingen B, et al. Development of a donor-acceptor concept for enzymatic cross-coupling reactions of aldehydes: the first asymmetric cross-benzoin condensation. J Am Chem Soc 2002;124:12084–5. 10.1021/ja0271476.

[4] Gocke D, Walter L, Gauchenova E, Kolter G, Knoll M, Berthold CL, et al. Rational protein design of ThDP-dependent enzymes: engineering stereoselectivity. ChemBioChem 2008;9:406–12. 10.1002/cbic.200700598.

[5] Knoll M, Müller M, Pleiss J, Pohl M. Factors mediating activity, selectivity, and substrate specificity for the thiamin diphosphate-dependent enzymes benzaldehyde lyase and benzoylformate decarboxylase. ChemBioChem 2006;7:1928–34. 10.1002/cbic.200600277.

[6] Pohl M, Sprenger GA, Müller M. A new perspective on thiamine catalysis. Curr Opin Biotechnol 2004;15:335–42. 10.1016/j.copbio.2004.06.002.

[7] Rother neé Gocke D, Kolter G, Gerhards T, Berthold CL, Gauchenova E, Knoll M, et al. S-Selective Mixed Carboligation by Structure-Based Design of the Pyruvate Decarboxylase from Acetobacter pasteurianus. ChemCatChem 2011;3:1587–96. 10.1002/cctc.201100054.

[8] Sehl T, Bock S, Marx L, Maugeri Z, Walter L, Westphal R, et al. Asymmetric synthesis of (S)-phenylacetylcarbinol – closing a gap in C–C bond formation. Green Chem 2017;19:380–4. 10.1039/C6GC01803C.

[9] Westphal R, Vogel C, Schmitz C, Pleiss J, Müller M, Pohl M, et al. A Tailor-Made Chimeric Thiamine Diphosphate Dependent Enzyme for the Direct Asymmetric Synthesis of (*S*)-Benzoins. Angew Chemie Int Ed 2014;53:9376–9. 10.1002/anie.201405069.

[10] Müller M, Gocke D, Pohl M. Thiamin diphosphate in biological chemistry: exploitation of diverse thiamin diphosphate-dependent enzymes for asymmetric chemoenzymatic synthesis. FEBS J 2009;276:2894–940. 10.1111/j.1742-4658.2009.07017.x.

[11] Gerhards T, Mackfeld U, Bocola M, von Lieres E, Wiechert W, Pohl M, et al. Influence of Organic Solvents on Enzymatic Asymmetric Carboligations. Adv Synth Catal 2012;354:2805–20. 10.1002/adsc.201200284.

[12] Demir ASS, Dünnwald T, Iding H, Pohl M, Müller M. Asymmetric benzoin reaction catalyzed by benzoylformate decarboxylase. Tetrahedron: Asymmetry 1999;10:4769–74. 10.1016/S0957-4166(99)00516-9.

[13] González B, Vicuña R. Benzaldehyde lyase, a novel thiamine PPi-requiring enzyme, from *Pseudomonas fluorescens* biovar I. J Bacteriol 1989;171:2401–5. 10.1128/jb.171.5.2401-2405.1989.

[14] Iding H, Dünnwald T, Greiner L, Liese A, Müller M, Siegert P, et al. Benzoylformate decarboxylase from *Pseudomonas putida* as stable catalyst for the synthesis of chiral 2-hydroxy ketones. Chem - A Eur J 2000;6:1483–95. 10.1002/(SICI)1521-3765(20000417)6:8<1483::AID-CHEM1483>3.0.CO;2-S.

[15] Polovnikova ES, McLeish MJ, Sergienko EA, Burgner JT, Anderson NL, Bera AK, et al. Structural and kinetic analysis of catalysis by a thiamin diphosphate-dependent enzyme, benzoylformate decarboxylase. Biochemistry 2003;42:1820–30. 10.1021/bi026490k.

[16] Gocke D, Nguyen CL, Pohl M, Stillger T, Walter L, Müller M. Branched-Chain Keto Acid Decarboxylase from *Lactococcus lactis* (KdcA), a Valuable Thiamine Diphosphate-Dependent Enzyme for Asymmetric C-C Bond Formation. Adv Synth Catal 2007;349:1425–35. 10.1002/adsc.200700057.

[17] Smit BA, van Hylckama Vlieg JET, Engels WJM, Meijer L, Wouters JTM, Smit G. Identification, Cloning, and Characterization of a Lactococcus lactis Branched-Chain α-Keto Acid Decarboxylase Involved in Flavor Formation. Appl Environ Microbiol 2005;71:303–11. 10.1128/AEM.71.1.303-311.2005.

[18] Gocke D, Graf T, Brosi H, Frindi-Wosch I, Walter L, Müller M, et al. Comparative characterisation of thiamin diphosphate-dependent decarboxylases. J Mol Catal B Enzym 2009;61:30–5. 10.1016/j.molcatb.2009.03.019.

[19] Cvjetko Bubalo M, Radošević K, Radojčić Redovniković I, Halambek J, Gaurina Srček V. A brief overview of the potential environmental hazards of ionic liquids. Ecotoxicol Environ Saf 2014;99:1–12. 10.1016/j.ecoenv.2013.10.019.

[20] Dupont J, Leal BC, Lozano P, Monteiro AL, Migowski P, Scholten JD. Ionic Liquids in Metal, Photo-, Electro-, and (Bio) Catalysis. Chem Rev 2024;124:5227–420. 10.1021/acs.chemrev.3c00379.

[21] Tiano M, Clark R, Bourgeois L, Costa Gomes M. The dialkylcarbonate route to ionic liquids: purer, safer, greener? Green Chem 2023;25:2541–58. 10.1039/d2gc04065d.

[22] Wells AS, Coombe VT. On the freshwater ecotoxicity and biodegradation properties of some common ionic liquids. Org Process Res Dev 2006;10:794–8. 10.1021/op060048i.

[23] Sheldon RA. Biocatalysis in ionic liquids: State-of-the-union. Green Chem 2021;23:8406–27. 10.1039/d1gc03145g.

[24] Lozano P, editor. Biocatalysis in Green Solvents. Academic Press, Inc.; 2022. 10.1016/C2021-0-00004-3.

[25] Xu P, Liang S, Zong MH, Lou WY. Ionic liquids for regulating biocatalytic process: Achievements and perspectives. Biotechnol Adv 2021;51:107702. 10.1016/j.biotechadv.2021.107702.

[26] Park S, Kazlauskas RJ. Biocatalysis in ionic liquids – advantages beyond green technology. Curr Opin Biotechnol 2003;14:432–7. 10.1016/S0958-1669(03)00100-9.

[27] Pramanik S, Dhoke G V., Jaeger KE, Schwaneberg U, Davari MD. How to Engineer Ionic Liquids Resistant Enzymes: Insights from Combined Molecular Dynamics and Directed Evolution Study. ACS Sustain Chem Eng 2019;7:11293–302. 10.1021/acssuschemeng.9b00752.

[28] Nordwald EM, Kaar JL. Stabilization of enzymes in ionic liquids via modification of enzyme charge. Biotechnol Bioeng 2013;110:2352–60. 10.1002/bit.24910.

[29] Zhao J, Frauenkron-Machedjou VJ, Fulton A, Zhu L, Davari MD, Jaeger KE, et al. Unraveling the effects of amino acid substitutions enhancing lipase resistance to an ionic liquid: A molecular dynamics study. Phys Chem Chem Phys 2018;20:9600–9. 10.1039/c7cp08470f.

[30] El Harrar T, Frieg B, Davari MD, Jaeger KE, Schwaneberg U, Gohlke H. Aqueous ionic liquids redistribute local enzyme stability via long-range perturbation pathways. Comput Struct Biotechnol J 2021;19:4248–64. 10.1016/j.csbj.2021.07.001.

[31] El Harrar T, Davari MD, Jaeger KE, Schwaneberg U, Gohlke H. Critical assessment of structure-based approaches to improve protein resistance in aqueous ionic liquids by enzyme-wide saturation mutagenesis. Comput Struct Biotechnol J 2022;20:399–409. 10.1016/j.csbj.2021.12.018.

[32] Schindl A, Hagen ML, Cooley I, Jäger CM, Warden AC, Zelzer M, et al. Ion-combination specific effects driving the enzymatic activity of halophilic alcohol dehydrogenase 2 from Haloferax volcanii in aqueous ionic liquid solvent mixtures. RSC Sustain 2024;2:2559–80. 10.1039/d3su00412k.

[33] Piccoli V, Martínez L. Competitive Effects of Anions on Protein Solvation by Aqueous Ionic Liquids. J Phys Chem B 2024;128:7792–802. 10.1021/acs.jpcb.4c03735.

[34] El Harrar T, Gohlke H. Cumulative Millisecond-Long Sampling for a Comprehensive Energetic Evaluation of Aqueous Ionic Liquid Effects on Amino Acid Interactions. J Chem Inf Model 2023;63:281–98. 10.1021/acs.jcim.2c01123.

[35] Fischer F, Mutschler J, Zufferey D. Enzyme catalysis with small ionic liquid quantities. J Ind Microbiol Biotechnol 2011;38:477–87. 10.1007/s10295-010-0908-1.

[36] Weingärtner H, Cabrele C, Herrmann C. How ionic liquids can help to stabilize native proteins. Phys Chem Chem Phys 2012;14:415–26. 10.1039/C1CP21947B.

[37] Yang Z, Yue Y-J, Huang W-C, Zhuang X-M, Chen Z-T, Xing M. Importance of the Ionic Nature of Ionic Liquids in Affecting Enzyme Performance. J Biochem 2009;145:355–64. 10.1093/jb/mvn173.

[38] Pilissão C, Nascimento MDG. Effects of organic solvents and ionic liquids on the aminolysis of (RS)-methyl mandelate catalyzed by lipases. Tetrahedron: Asymmetry 2006;17:428–33. 10.1016/j.tetasy.2006.02.001.

[39] Schöfer SH, Kaftzik N, Kragl U, Wasserscheid P. Enzyme catalysis in ionic liquids: lipase catalysed kinetic resolution of 1-phenylethanol with improved enantioselectivity. Chem Commun 2001:425–6. 10.1039/b009389k.

[40] Nara SJ, Mohile SS, Harjani JR, Naik PU, Salunkhe MM. Influence of ionic liquids on the rates and regioselectivity of lipase-mediated biotransformations on 3,4,6-tri-O-acetyl-d-glucal. J Mol Catal B Enzym 2004;28:39–43. 10.1016/j.molcatb.2004.01.010.

[41] Park S, Kazlauskas RJ. Improved Preparation and Use of Room-Temperature Ionic Liquids in Lipase-Catalyzed Enantio- and Regioselective Acylations. J Org Chem 2001;66:8395–401. 10.1021/jo015761e.

[42] Frauenkron-Machedjou VJ, Fulton A, Zhu L, Anker C, Bocola M, Jaeger K-E, et al. Towards Understanding Directed Evolution: More than Half of All Amino Acid Positions Contribute to Ionic Liquid Resistance of *Bacillus subtilis* Lipase A. ChemBioChem 2015;16:937–45. 10.1002/cbic.201402682.

[43] Nutschel C, Fulton A, Zimmermann O, Schwaneberg U, Jaeger KE, Gohlke H. Systematically Scrutinizing the Impact of Substitution Sites on Thermostability and Detergent Tolerance for *Bacillus* subtilis Lipase A. J Chem Inf Model 2020;60:1568–84. 10.1021/acs.jcim.9b00954.

[44] Kokova M, Zavrel M, Tittmann K, Spiess AC, Pohl M. Investigation of the carboligase activity of thiamine diphosphate-dependent enzymes using kinetic modeling and NMR spectroscopy. J Mol Catal B Enzym 2009;61:73–9. 10.1016/j.molcatb.2009.02.021.

[45] Janzen E, Müller M, Kolter-Jung D, Kneen MM, McLeish MJ, Pohl M. Characterization of benzaldehyde lyase from *Pseudomonas fluorescens*: A versatile enzyme for asymmetric C-C bond formation. Bioorg Chem 2006;34:345–61. 10.1016/j.bioorg.2006.09.002.

[46] Gerhards T. Einfluss unkonventioneller Medien auf die Selektivität ThDP-abhängiger Enzyme. doctoral thesis, Heinrich-Heine University Düsseldorf, 2012. https://docserv.uni-duesseldorf.de/servlets/DocumentServlet?id=24413.

[47] Magnuson DK, Bodley JW, Evans DF. The activity and stability of alkaline phosphatase in solutions of water and the fused salt ethylammonium nitrate. J Solution Chem 1984;13:583–7. 10.1007/BF00647226.

[48] Dreyer S, Kragl U. Ionic liquids for aqueous two-phase extraction and stabilization of enzymes. Biotechnol Bioeng 2008;99:1416–24. 10.1002/bit.21720.

[49] Holbrey JD, Turner MB, Reichert WM, Rogers RD. New ionic liquids containing an appended hydroxyl functionality from the atom-efficient, one-pot reaction of 1-methylimidazole and acid with propylene oxide. Green Chem 2003;5:731–6. 10.1039/b311717k.

[50] Ríos-Lombardía N, Busto E, Gotor-Fernández V, Gotor V, Porcar R, García-Verdugo E, et al. From salts to ionic liquids by systematic structural modifications: A rational approach towards the efficient modular synthesis of enantiopure imidazolium salts. Chem - A Eur J 2010;16:836–47. 10.1002/chem.200901623.

[51] Paul CE, Gotor-Fernández V, Lavandera I, Montejo-Bernardo J, García-Granda S, Gotor V. Chemoenzymatic preparation of optically active 3-(1H-imidazol-1-yl) cyclohexanol-based ionic liquids: Application in organocatalysis and toxicity studies. RSC Adv 2012;2:6455–63. 10.1039/c2ra20876h.

[52] Schrödinger release 2018-4: Maestro. LLC, New York: Schrödinger; n.d.

[53] Paulikat M, Wechsler C, Tittmann K, Mata RA. Theoretical Studies of the Electronic Absorption Spectra of Thiamin Diphosphate in Pyruvate Decarboxylase. Biochemistry 2017;56:1854–64. 10.1021/acs.biochem.6b00984.

[54] Šali A, Blundell TL. Comparative Protein Modelling by Satisfaction of Spatial Restraints. J Mol Biol 1993;234:779–815. 10.1006/jmbi.1993.1626.

[55] Shelley JC, Cholleti A, Frye LL, Greenwood JR, Timlin MR, Uchimaya M. Epik: A software program for pKa prediction and protonation state generation for drug-like molecules. J Comput Aided Mol Des 2007;21:681–91. 10.1007/s10822-007-9133-z.

[56] Schrödinger suite 2018-4 protein preparation wizard. LLC, New York: Schrödinger; n.d.

[57] Schafmeister CEAF, Ross WS, Romanowski V. No Title. San Francisco: University of California, San Fransciso; 1995.

[58] Tian C, Kasavajhala K, Belfon KAA, Raguette L, Huang H, Migues AN, et al. Ff19SB: Amino-Acid-Specific Protein Backbone Parameters Trained against Quantum Mechanics Energy Surfaces in Solution. J Chem Theory Comput 2020;16:528–52. 10.1021/acs.jctc.9b00591.

[59] Case DA, Belfon K, Ben-Shalom IY, Brozell SR, Cerutti DS, Cheatham III. TE, et al. Amber 2020. University of California, San Fransciso; 2020.

[60] Frisch MJ, Trucks GW, Schlegel HB, Scuseria GE, Robb MA, Cheeseman JR, et al. Gaussian 16 Rev. A. 03. Wallingford CT: Gaussian, Inc.; 2016.

[61] Roothaan CCJ. New developments in molecular orbital theory. Rev Mod Phys 1951;23:69–89. 10.1103/RevModPhys.23.69.

[62] Bayly CI, Cieplak P, Cornell WD, Kollman PA. A well-behaved electrostatic potential based method using charge restraints for deriving atomic charges: The RESP model. J Phys Chem 1993;97:10269–80. 10.1021/j100142a004.

[63] Minisini B, Chavand S, Barthelery R, Tsobnang F. Calculations of the charge distribution in dodecyltrimethylammonium: A quantum chemical investigation. J Mol Model 2010;16:1085–92. 10.1007/s00894-009-0620-0.

[64] Hoffmann MM, Too MD, Paddock NA, Horstmann R, Kloth S, Vogel M, et al. On the Behavior of the Ethylene Glycol Components of Polydisperse Polyethylene Glycol PEG200. J Phys Chem B 2023;127:1178–96. 10.1021/acs.jpcb.2c06773.

[65] Ensing B, Tiwari A, Tros M, Hunger J, Domingos SR, Pérez C, et al. On the origin of the extremely different solubilities of polyethers in water. Nat Commun 2019;10:2893. 10.1038/s41467-019-10783-z.

[66] He X, Man VH, Yang W, Lee T-S, Wang J. A fast and high-quality charge model for the next generation general AMBER force field. J Chem Phys 2020;153:114502. 10.1063/5.0019056.

[67] Martínez L, Andrade R, Birgin EG, Martínez JM. PACKMOL: A package for building initial configurations for molecular dynamics simulations. J Comput Chem 2009;30:2157–64. 10.1002/jcc.21224.

[68] Izadi S, Anandakrishnan R, Onufriev A V. Building Water Models: A Different Approach. J Phys Chem Lett 2014;5:3863–71. 10.1021/jz501780a.

[69] Roe DR, Brooks BR. A protocol for preparing explicitly solvated systems for stable molecular dynamics simulations. J Chem Phys 2020;153. 10.1063/5.0013849.

[70] Hopkins CW, Le Grand S, Walker RC, Roitberg AE. Long-time-step molecular dynamics through hydrogen mass repartitioning. J Chem Theory Comput 2015;11:1864–74. 10.1021/ct5010406.

[71] Salomon-Ferrer R, Götz AW, Poole D, Le Grand S, Walker RC. Routine Microsecond Molecular Dynamics Simulations with AMBER on GPUs. 2. Explicit Solvent Particle Mesh Ewald. J Chem Theory Comput 2013;9:3878–88. 10.1021/ct400314y.

[72] Darden T, York D, Pedersen L. Particle mesh Ewald: An N⋅log(N) method for Ewald sums in large systems. J Chem Phys 1993;98:10089–92. 10.1063/1.464397.

[73] Ryckaert JP, Ciccotti G, Berendsen HJC. Numerical integration of the cartesian equations of motion of a system with constraints: molecular dynamics of n-alkanes. J Comput Phys 1977;23:327–41. 10.1016/0021-9991(77)90098-5.

[74] Chovancova E, Pavelka A, Benes P, Strnad O, Brezovsky J, Kozlikova B, et al. CAVER 3.0: A Tool for the Analysis of Transport Pathways in Dynamic Protein Structures. PLoS Comput Biol 2012;8:23–30. 10.1371/journal.pcbi.1002708.

[75] Virtanen P, Gommers R, Oliphant TE, Haberland M, Reddy T, Cournapeau D, et al. SciPy 1.0: fundamental algorithms for scientific computing in Python. Nat Methods 2020;17:261–72. 10.1038/s41592-019-0686-2.

[76] McKinney W. Data Structures for Statistical Computing in Python. Proc. 9th PYTHON Sci. conf. (SCIPY 2010), 2010, p. 56–61. 10.25080/Majora-92bf1922-00a.

[77] Jordan F, Nemeria NS. Progress in the experimental observation of thiamin diphosphate-bound intermediates on enzymes and mechanistic information derived from these observations. Bioorg Chem 2014;57:251–62. 10.1016/j.bioorg.2014.08.002.

[78] Roe DR, Cheatham TE. PTRAJ and CPPTRAJ: Software for Processing and Analysis of Molecular Dynamics Trajectory Data. J Chem Theory Comput 2013;9:3084–95. 10.1021/ct400341p.

[79] Goldfeder M, Egozy M, Shuster Ben-Yosef V, Adir N, Fishman A. Changes in tyrosinase specificity by ionic liquids and sodium dodecyl sulfate. Appl Microbiol Biotechnol 2013;97:1953–61. 10.1007/s00253-012-4050-z.

[80] Domínguez de María P, Pohl M, Gocke D, Gröger H, Trauthwein H, Stillger T, et al. Asymmetric Synthesis of Aliphatic 2-Hydroxy Ketones by Enzymatic Carboligation of Aldehydes. European J Org Chem 2007;2007:2940–4. 10.1002/ejoc.200600876.

[81] Kim HS, Ha SH, Sethaphong L, Koo Y-M, Yingling YG. The relationship between enhanced enzyme activity and structural dynamics in ionic liquids: a combined computational and experimental study. Phys Chem Chem Phys 2014;16:2944. 10.1039/c3cp52516c.

[82] Berthold CL, Gocke D, Wood MD, Leeper FJ, Pohl M, Schneider G. Structure of the branched-chain keto acid decarboxylase (KdcA) from *Lactococcus lactis* provides insights into the structural basis for the chemoselective and enantioselective carboligation reaction. Acta Crystallogr Sect D Biol Crystallogr 2007;63:1217–24. 10.1107/S0907444907050433.

