## Supporting Information for "Influence of ionic liquids on enzymatic asymmetric carboligations"

#### Supplementary figures

##### Figure S1: Chemo- and stereoselectivity of *Ll*KdcA.

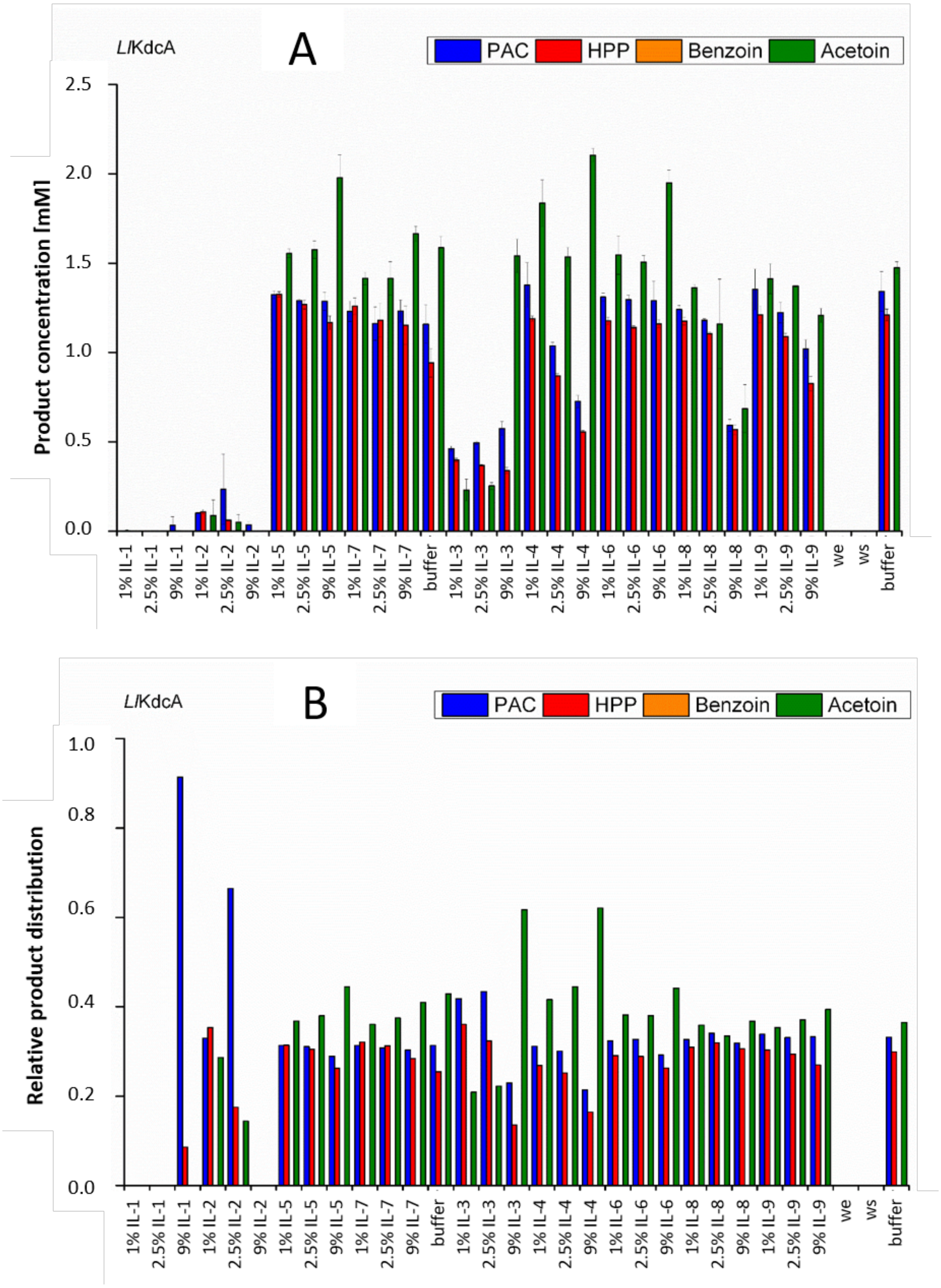

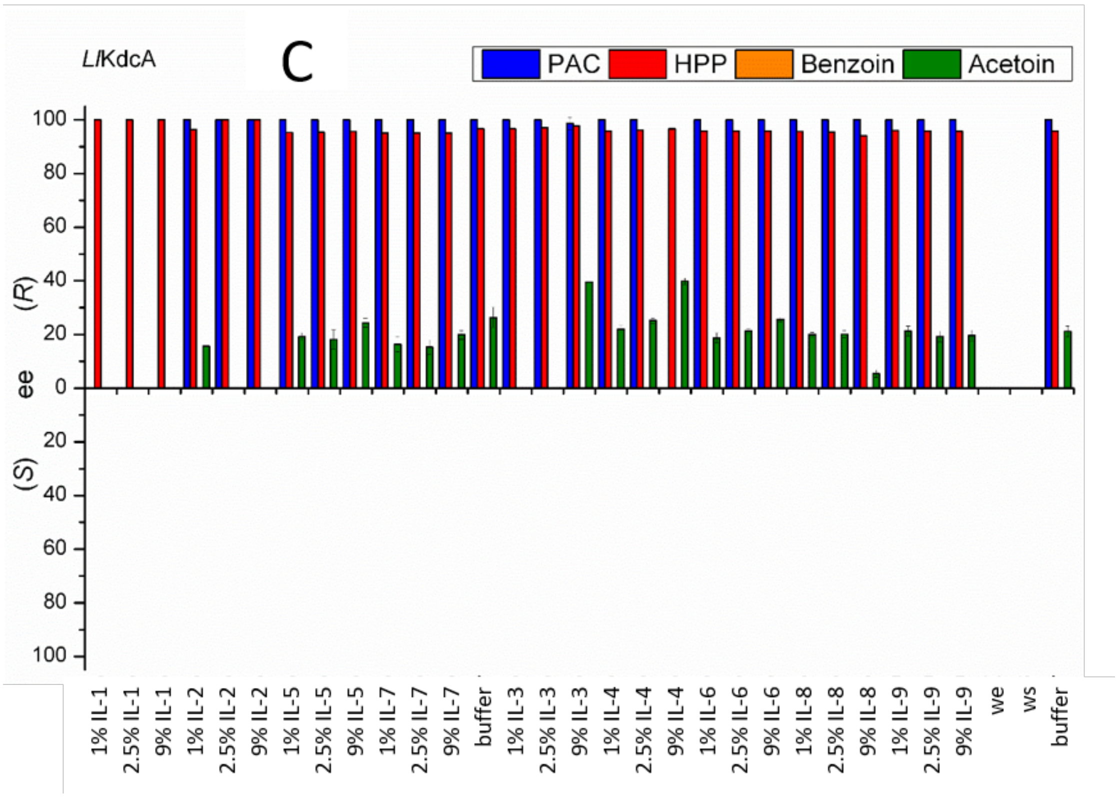


**Figure S1. Influence of IL addition on the chemo- and stereoselectivity of *Ll*KdcA-catalyzed carboligations of benzaldehyde and acetaldehyde. A:** Product concentration. **B**: Relative product distribution. **C**: Enantiomeric excess of obtained products. Reaction conditions: TEA-buffer pH 7.5 (50 mM, 2.5 mM $MgSO_4$, 0.1 mM ThDP), 0.1 mg/mL LlKdcA (if conversion was too little for product determination 0.4 mg/mL *Ll*KdcA was added), 180 mM acetaldehyde, 18 mM benzaldehyde. All bars represent the arithmetic average determined from three independent reactions. we = same reaction but without enzyme, ws = same reaction but without substrate; buffer = same reaction but without addition of ionic liquid.

Figure S2: Chemo- and stereoselectivity of *Pp*BFD.

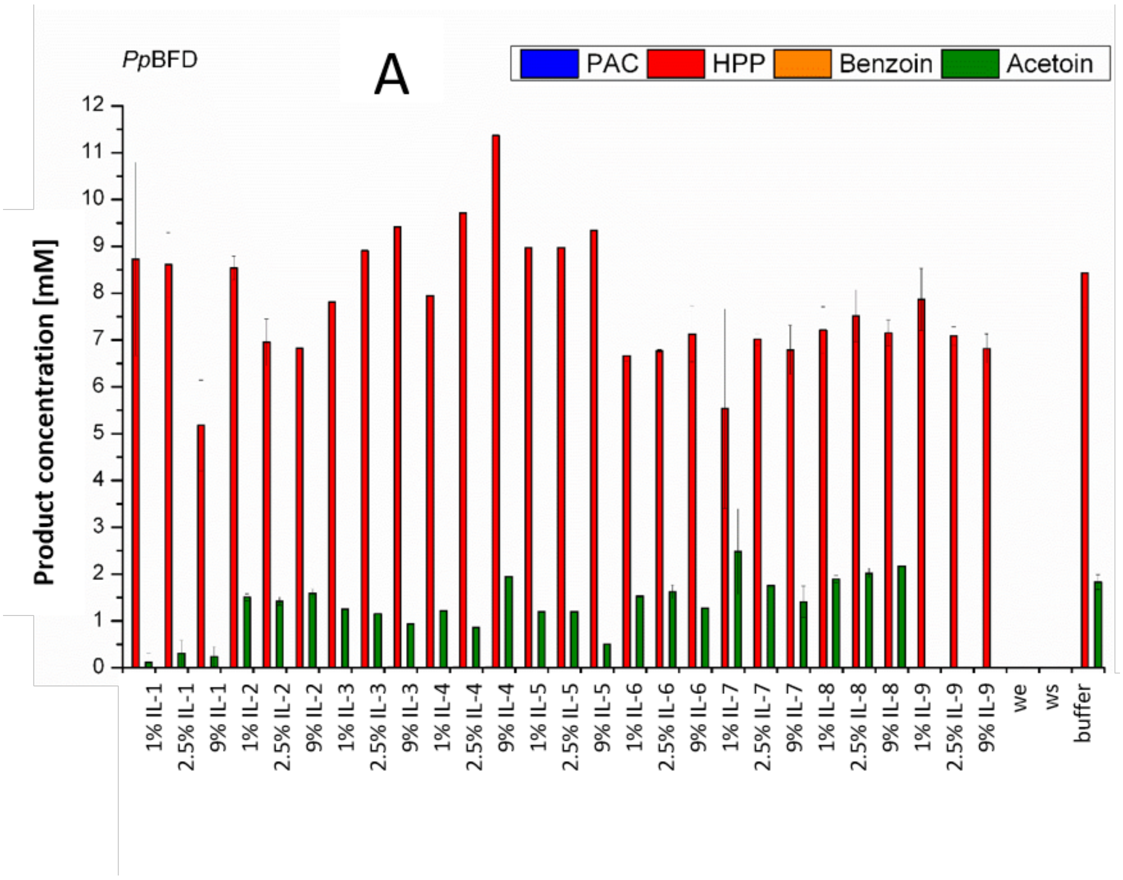


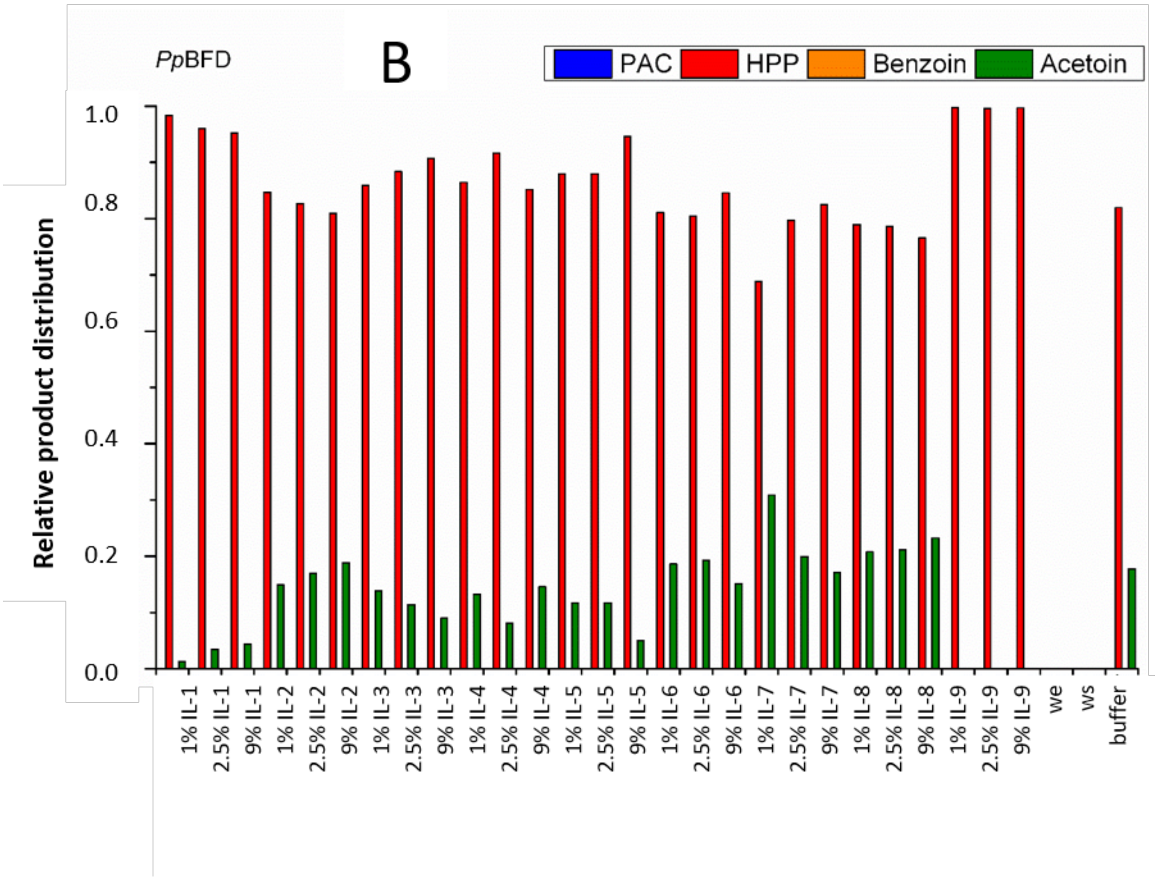

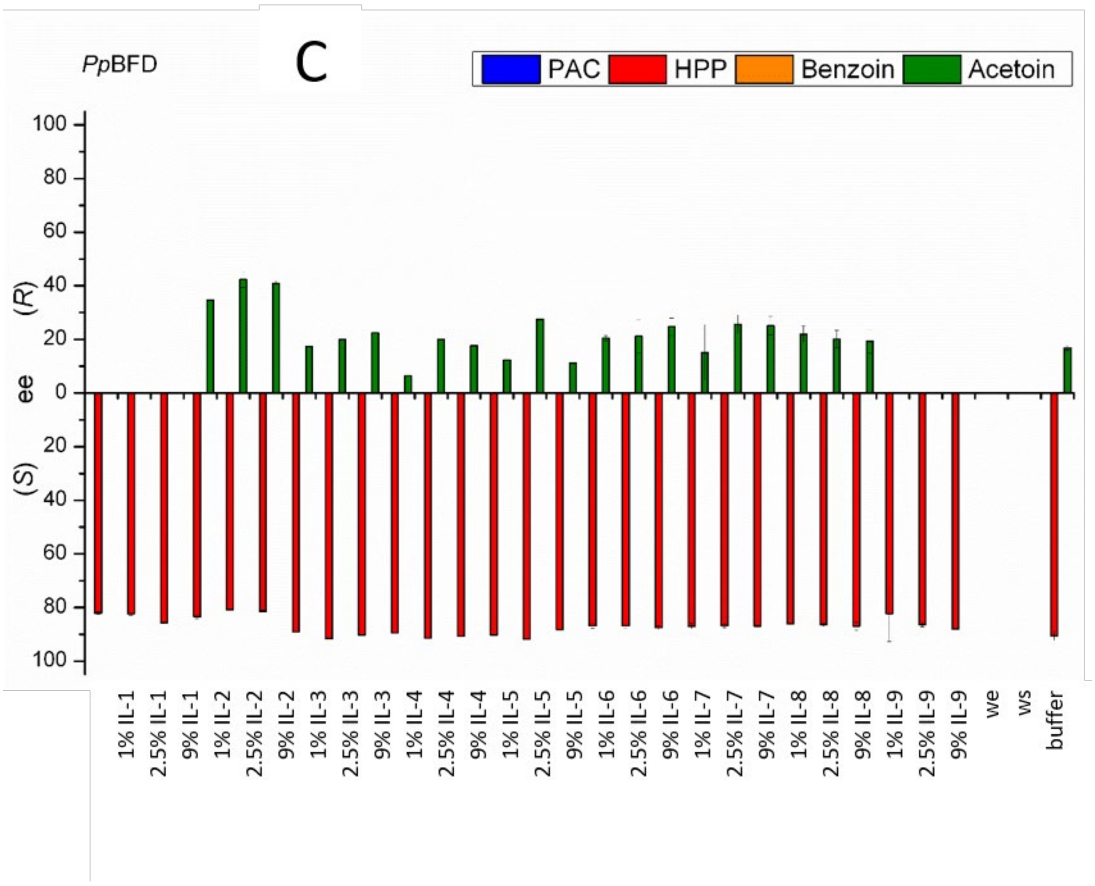


**Figure S2. Influence of IL addition on the chemo- and stereoselectivity of *Pp*BFD-catalyzed carboligations of benzaldehyde and acetaldehyde. A:** Product concentration. **B**: Relative product distribution. **C**: Enantiomeric excess of gained products. Reaction conditions: TEA-buffer pH 7.5 (50 mM, 2.5 mM $MgSO_4$, 0.1 mM ThDP), 0.1 mg/mL *Pp*BFD, 180 mM acetaldehyde, 18 mM benzaldehyde. All bars represent the arithmetic average determined from three independent reactions. we = same reaction but without enzyme, ws = same reaction but without substrate; buffer = same reaction but without addition of ionic liquid.

Figure S3: Chemo- and stereoselectivity of *Pp*BFD_H281A.

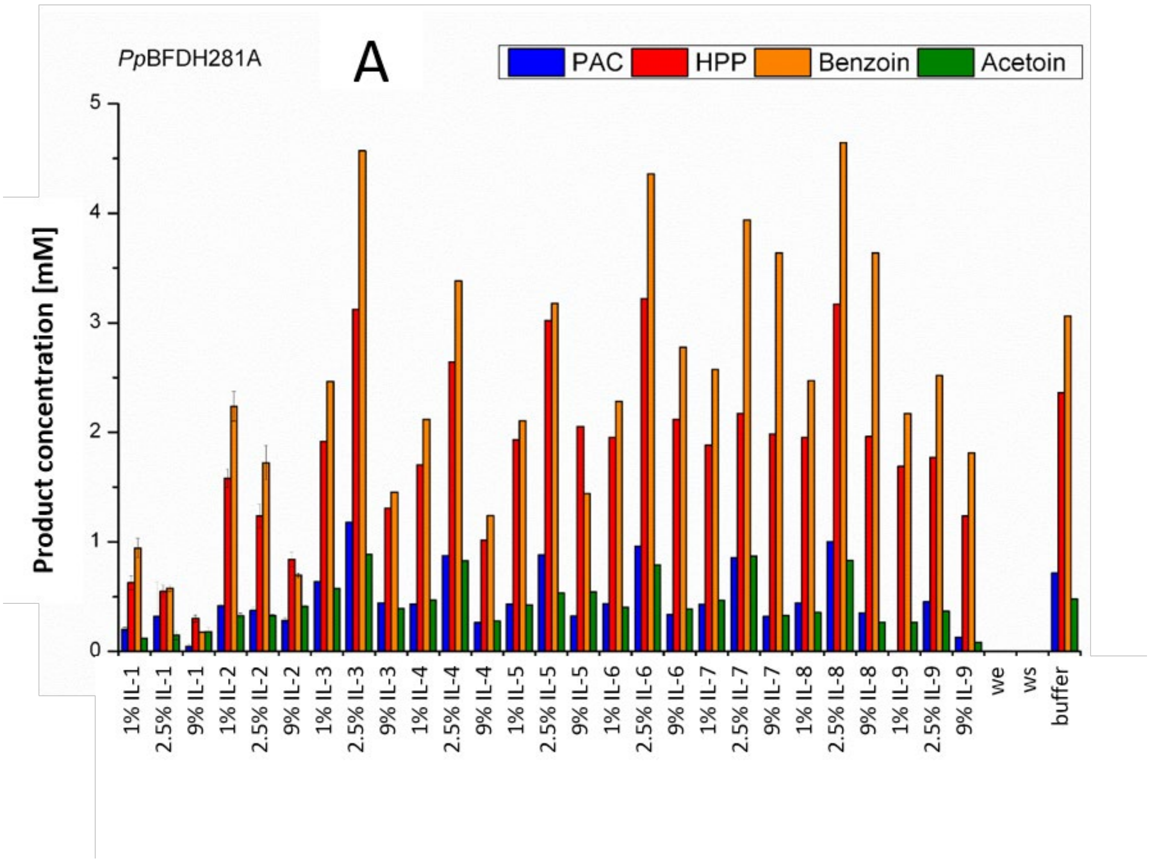


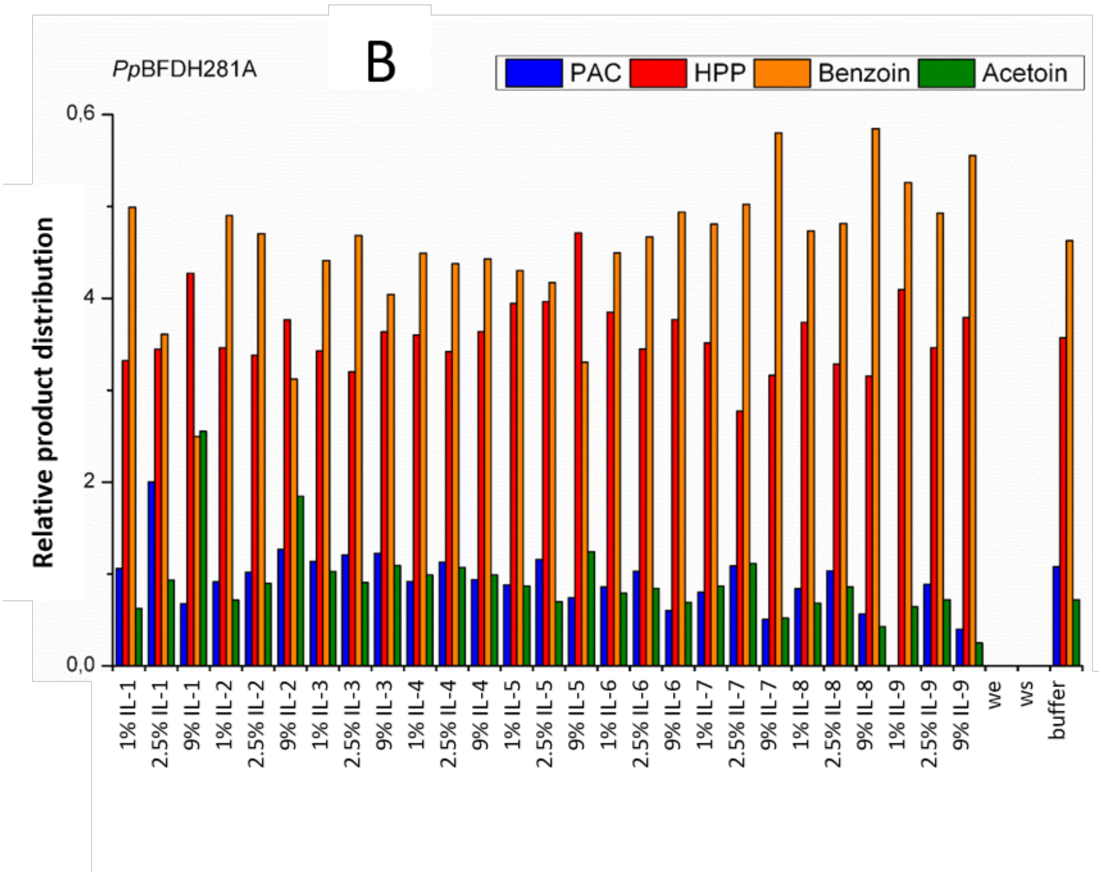

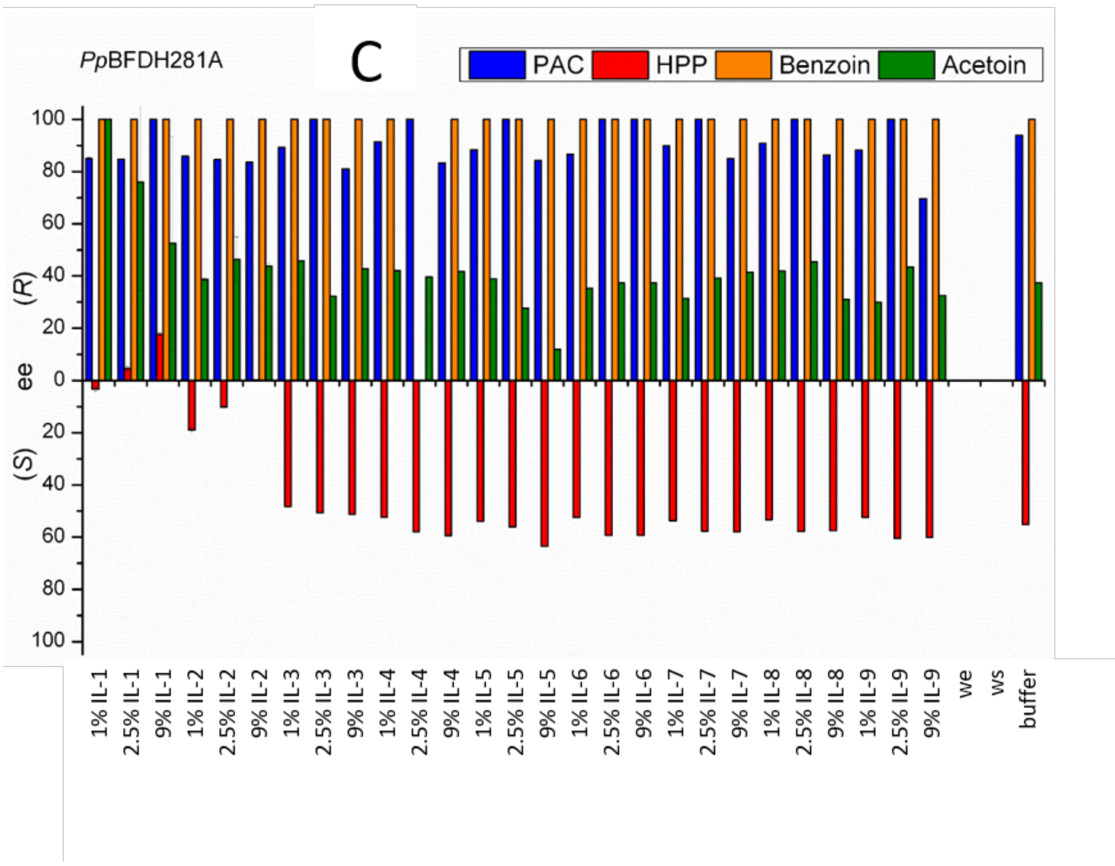


**Figure S3. Influence of IL addition on the chemo- and stereoselectivity of *Pp*BFD_H281A-catalyzed carboligations of benzaldehyde and acetaldehyde. A:** Product concentration. **B**: Relative product distribution. **C**: Enantiomeric excess of gained products. Reaction conditions: TEA-buffer pH 7.5 (50 mM, 2.5 mM $MgSO_4$, 0.1 mM ThDP), 0.1 mg/mL *Pp*BFD_H281A, 180 mM acetaldehyde, 18 mM benzaldehyde. All bars represent the arithmetic average determined from three independent reactions. we = same reaction but without enzyme, ws = same reaction but without substrate; buffer = same reaction but without addition of ionic liquid.

#### Figure S4: Chemo- and stereoselectivity of *Pf*BAL

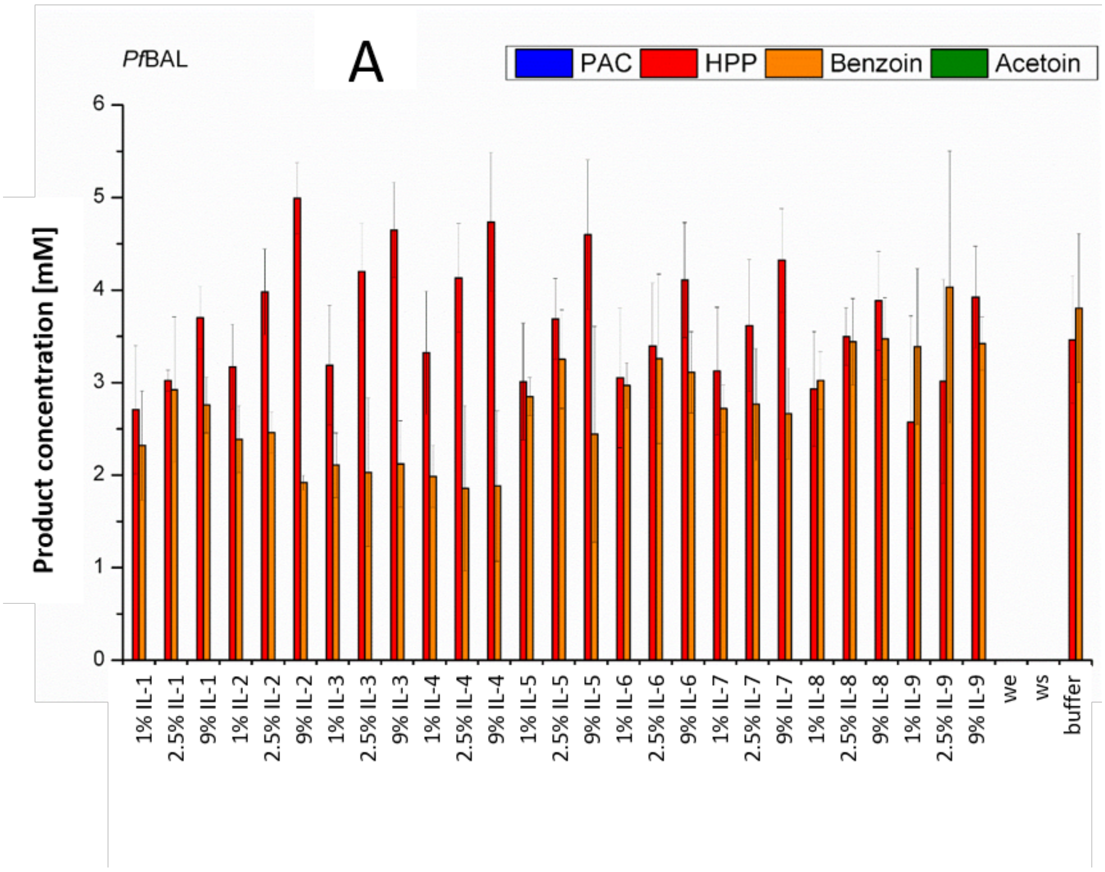


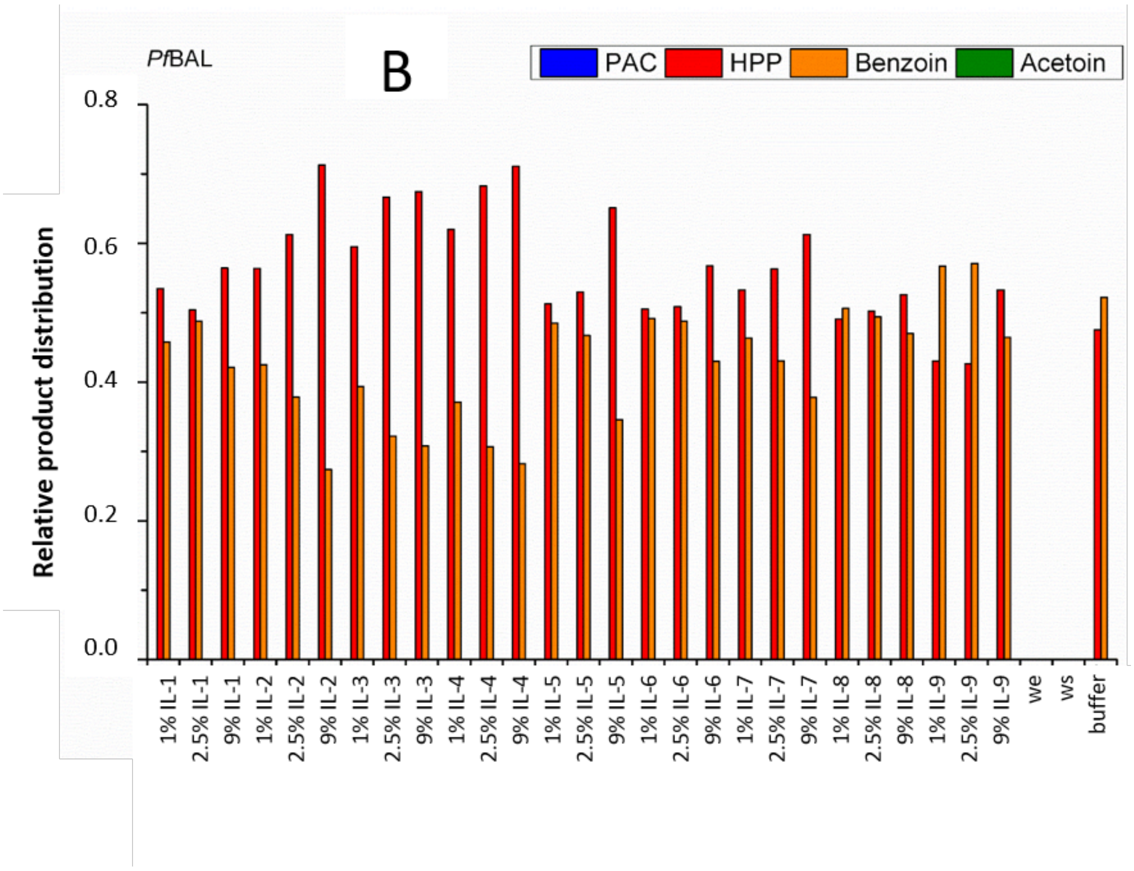

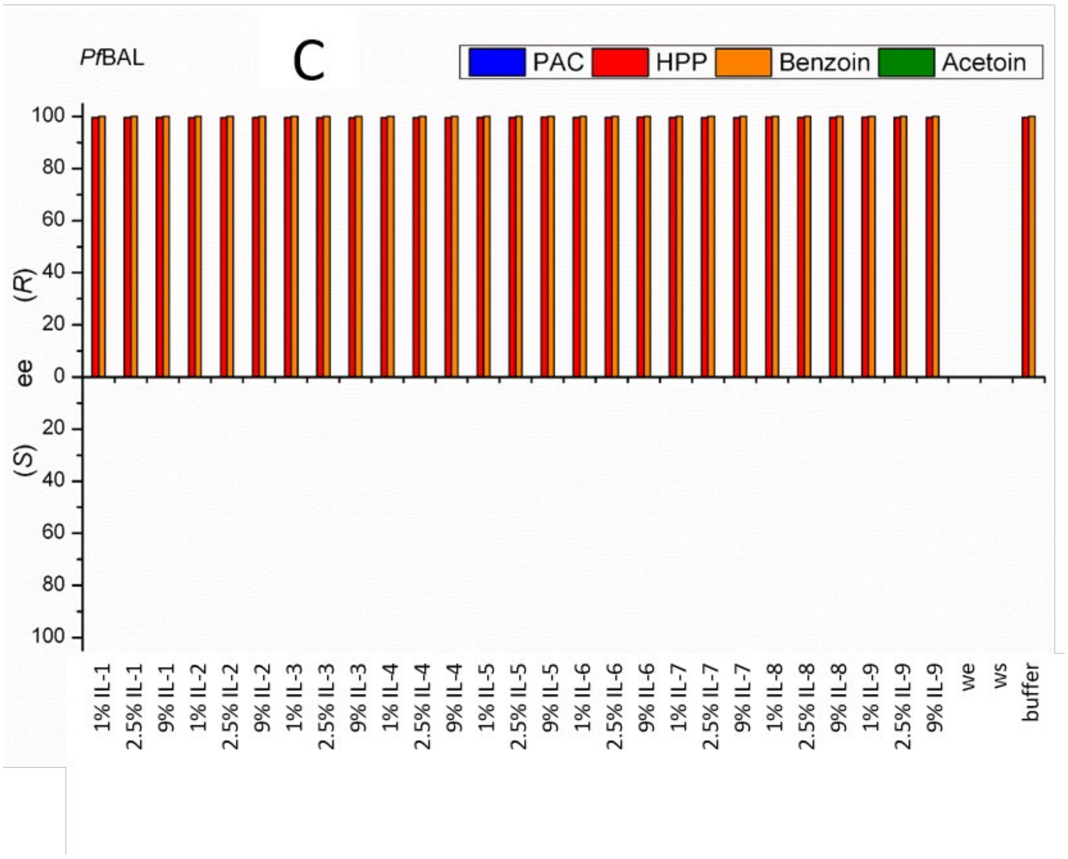


**Figure S4. Influence of IL addition on the chemo- and stereoselectivity of *Pf*BAL-catalyzed carboligations of benzaldehyde and acetaldehyde. A:** Product concentration. **B**: Relative product distribution. **C**: Enantiomeric excess of gained products. Reaction conditions: TEA-buffer pH 8.0 (50 mM, 2.5 mM $MgSO_4$, 0.1 mM ThDP), 0.02 mg/mL *Pf*BAL, 18 mM acetaldehyde, 18 mM benzaldehyde. All bars represent the arithmetic average determined from three independent reactions. we = same reaction but without enzyme, ws = same reaction but without substrate; buffer = same reaction but without addition of ionic liquid.

#### Figure S5: Chemo- and stereoselectivity of *Ap*PDC_WT

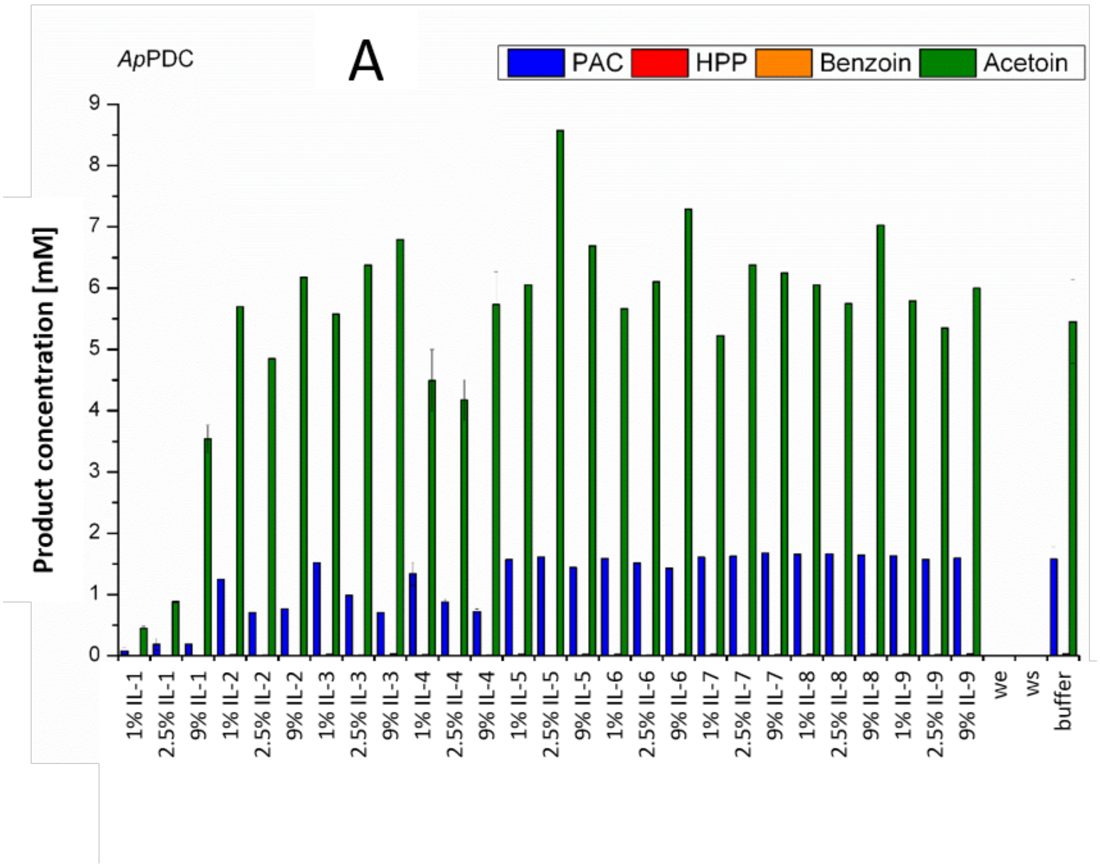


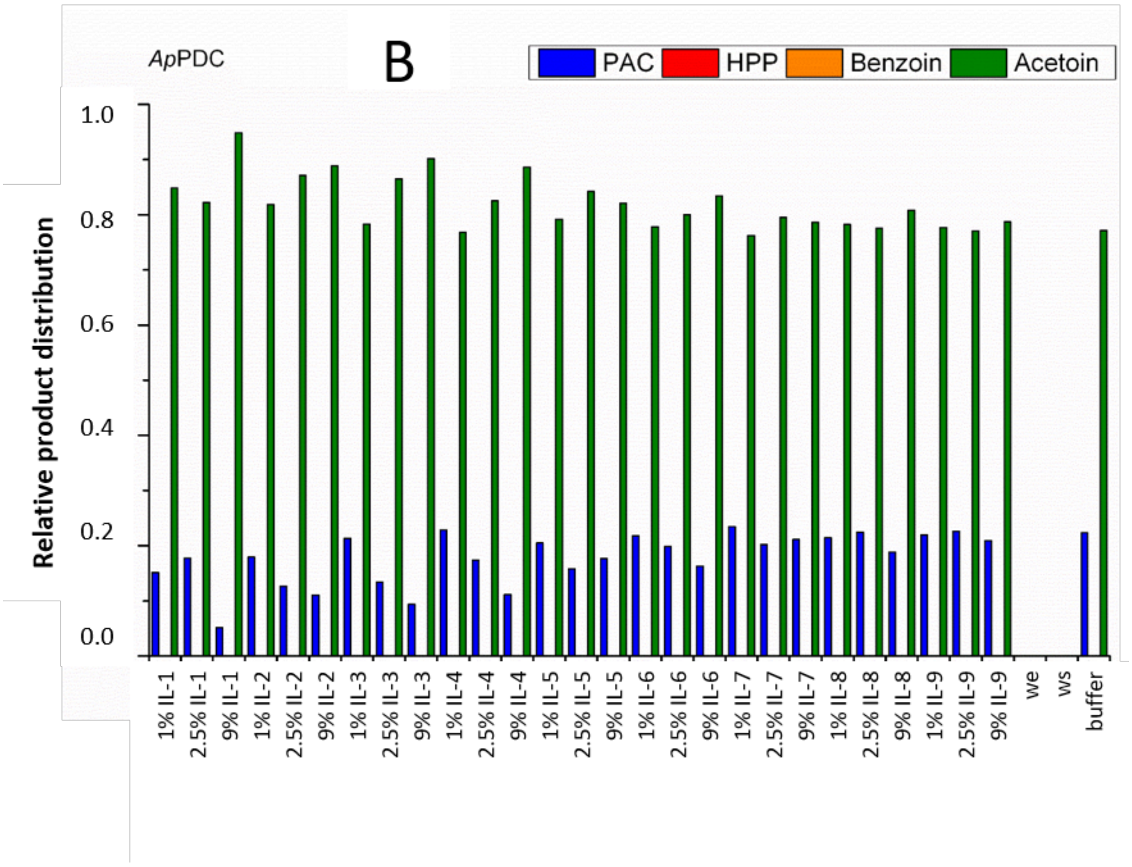

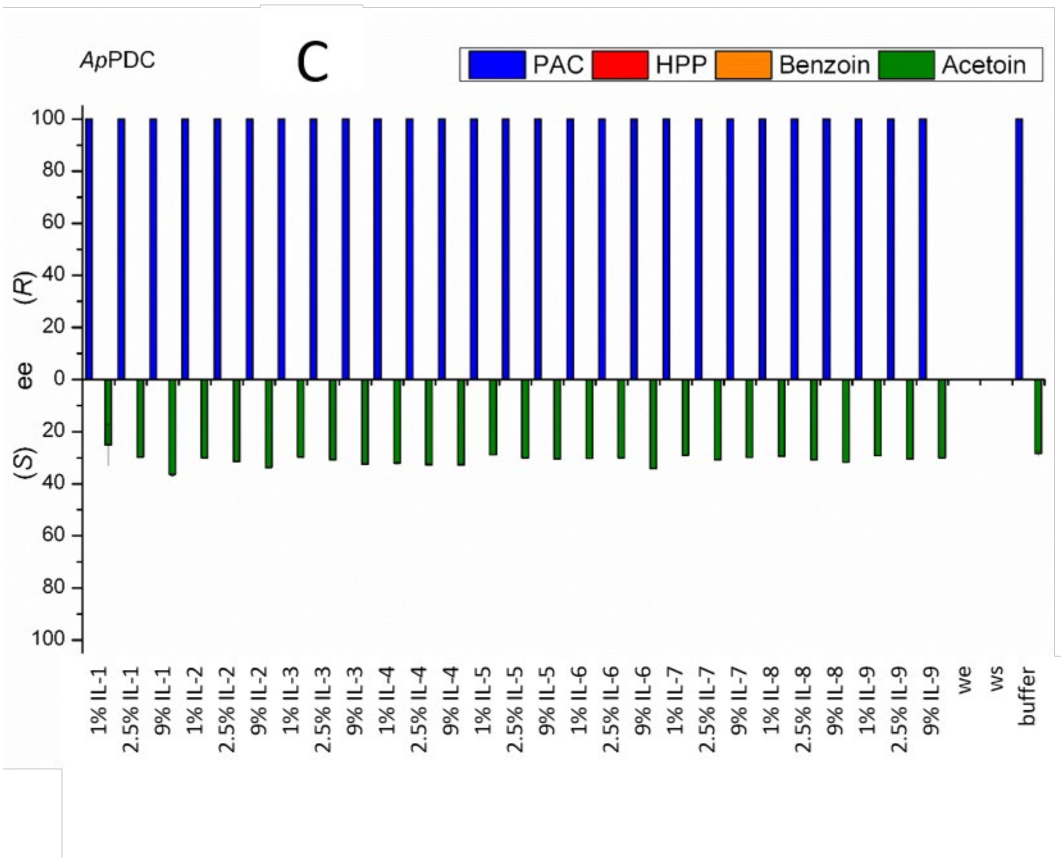


**Figure S5. Influence of IL addition on the chemo- and stereoselectivity of *Ap*PDC_WT-catalyzed carboligations of benzaldehyde and acetaldehyde. A:** Product concentration. **B**: Relative product distribution. **C**: Enantiomeric excess of gained products. Reaction conditions: TEA-buffer pH 7.5 (50 mM, 2.5 mM $MgSO_4$, 0.1 mM ThDP), 0.1 mg/mL *Ap*PDC_WT, 18 mM acetaldehyde, 18 mM benzaldehyde. All bars represent the arithmetic average determined from three independent reactions. we = same reaction but without enzyme, ws = same reaction but without substrate; buffer = same reaction but without addition of ionic liquid.

Figure S6: Chemo- and stereoselectivity of *Ap*PDC_E469G.

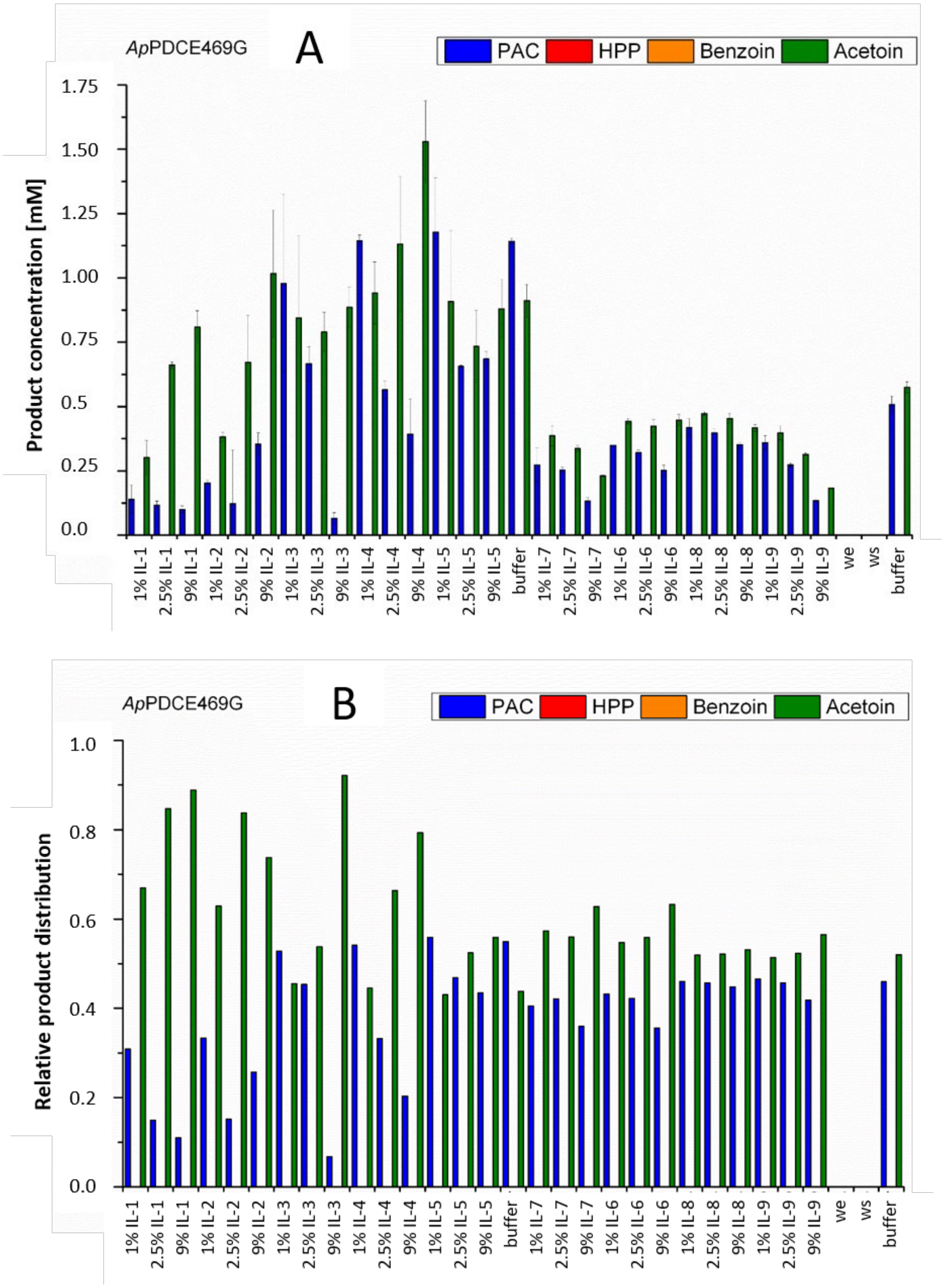

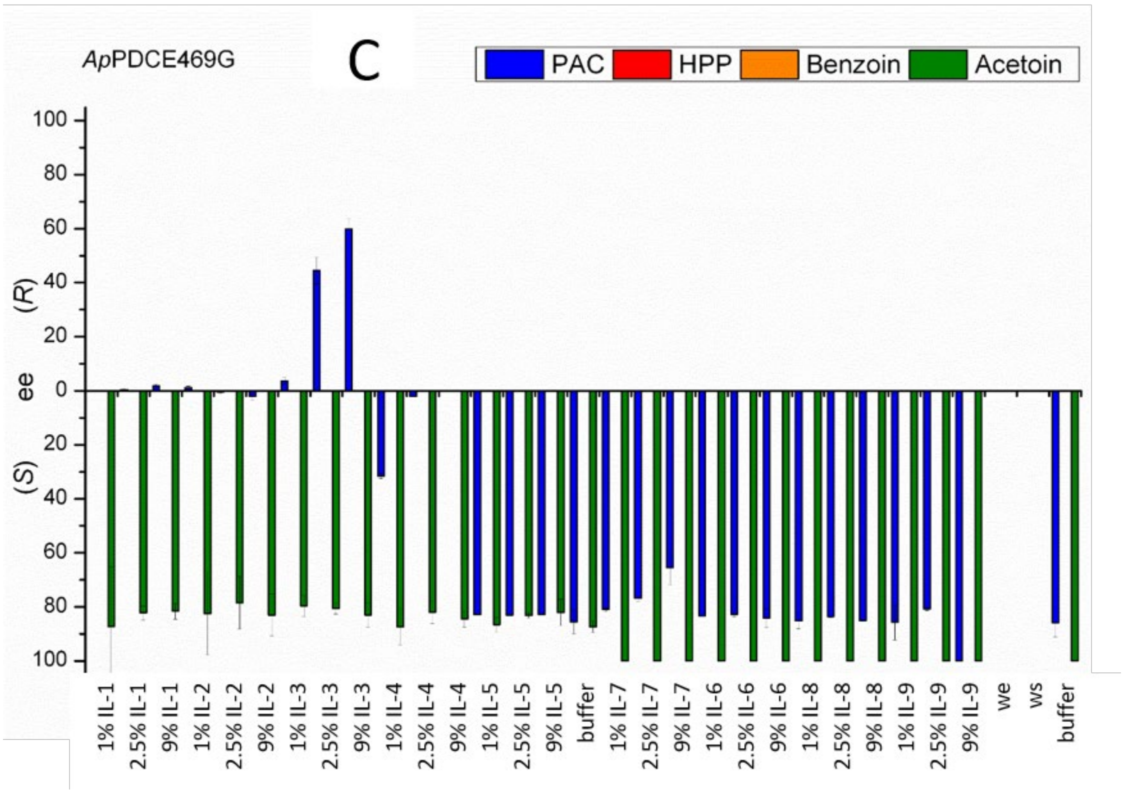


**Figure S6. Influence of IL addition on the chemo- and stereoselectivity of *Ap*PDC_E469G-catalyzed carboligations of benzaldehyde and acetaldehyde. A:** Product concentration. **B**: Relative product distribution. **C**: Enantiomeric excess of gained products. Reaction conditions: TEA-buffer pH 7.5 (50 mM, 2.5 mM $MgSO_4$, 0.1 mM ThDP), 0.1 mg/mL *Ap*PDC_E469G (if conversion was too little for product determination 0.4 mg/mL *Ap*PDC_E469G was added), 18 mM acetaldehyde, 18 mM benzaldehyde. All bars represent the arithmetic average determined from three independent reactions. we = same reaction but without enzyme, ws = same reaction but without substrate; buffer = same reaction but without addition of ionic liquid.

### Figure S7: Headspace-GC-MS of the IL Ammoeng 102 (IL-3).

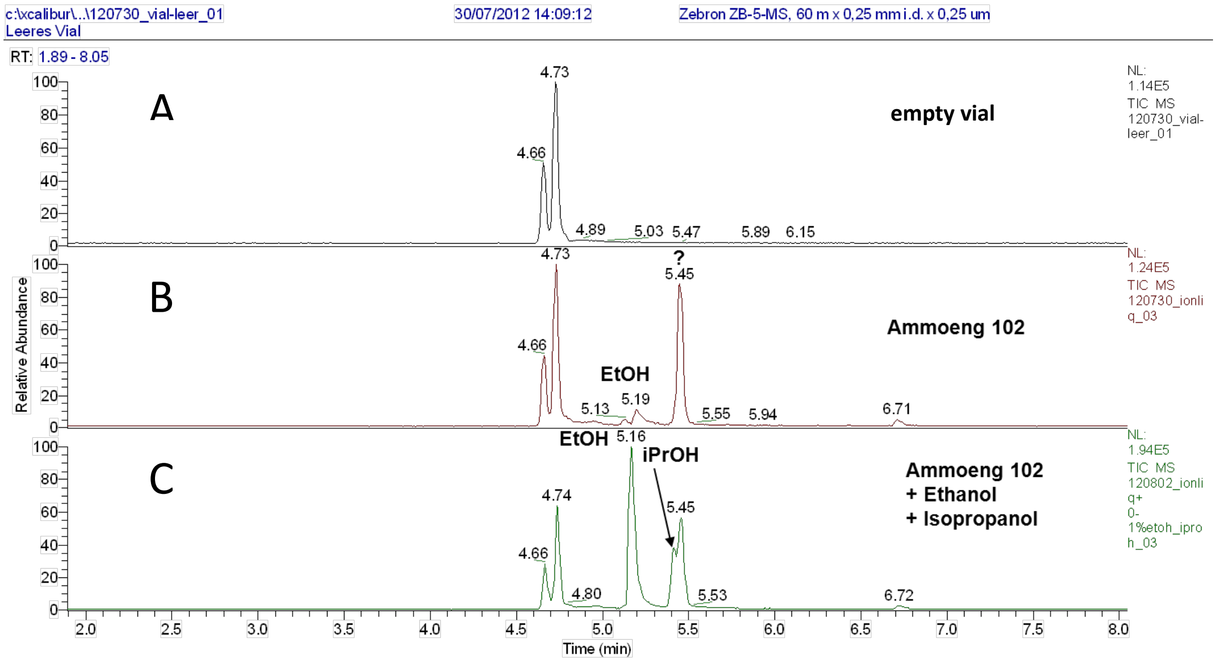


**Figure S7: Headspace-GC-MS of the IL Ammoeng 102 (IL-3). A**: Empty vial. **B**: Ammoeng 102 (IL-3). **C**: Ammoeng 102 (IL-3) with ethanol and isopropanol co-injected. Analysis of IL-3 revealed two additional signals at 5.19 min and 5.45 min (B). By co-injection of the solvents ethanol and isopropanol (C), ethanol could be assigned as the impurity. The identity of the second component could not be clarified. Estimated quantities: EtOH: 0.02% (w/v) and unknown compound: 0.06% (w/v). This measurement was carried out by the Central Department for Chemical Analysis (ZCH) of the Forschungszentrum Jülich GmbH.

### Figure S8: FIA-FTICR-MS of IL-3 Ammoeng 102.

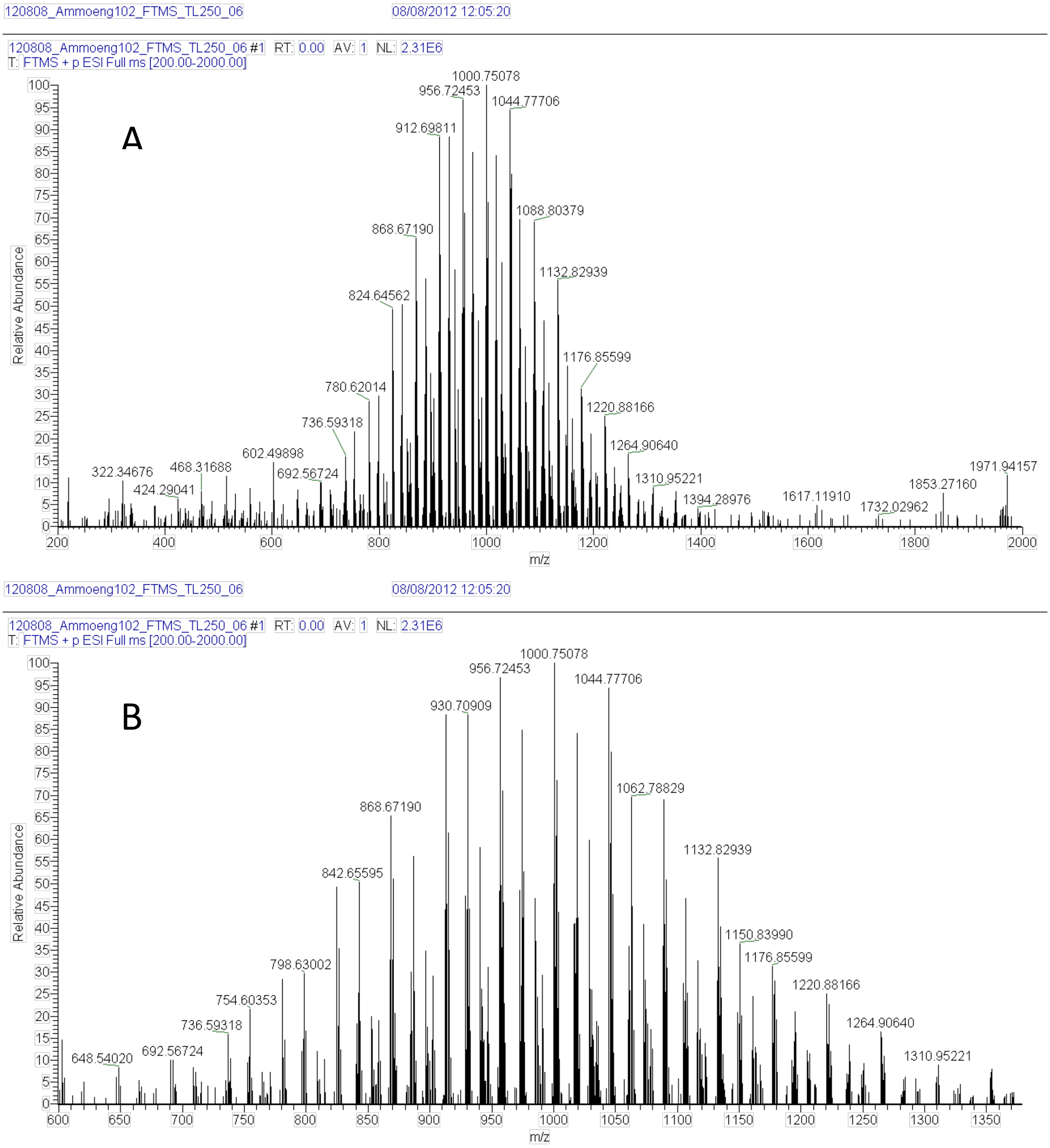


**Figure S8: Flow-Injection-Analysis-Fourier-Transformation-Ion-Cyclotron Resonance Mass Spectrometry (FIA-FTICR-MS) of IL-3 Ammoeng 102. A**. Signals in mass range 200 - 2000 m/Z. **B**. Signals in mass range 600 - 1400 m/z. The spectrum shows different Ammoeng 102 species with polyethylene glycol chains of different lengths, as well as variants with alkyl chains of different lengths (**for details see Text S5**). No evidence of polyethylene glycol as an impurity was found. This measurement was carried out by the Central Department for Chemical Analysis (ZCH) of the Forschungszentrum Jülich GmbH.

#### Figure S9: Calibration of the achiral HPLC.

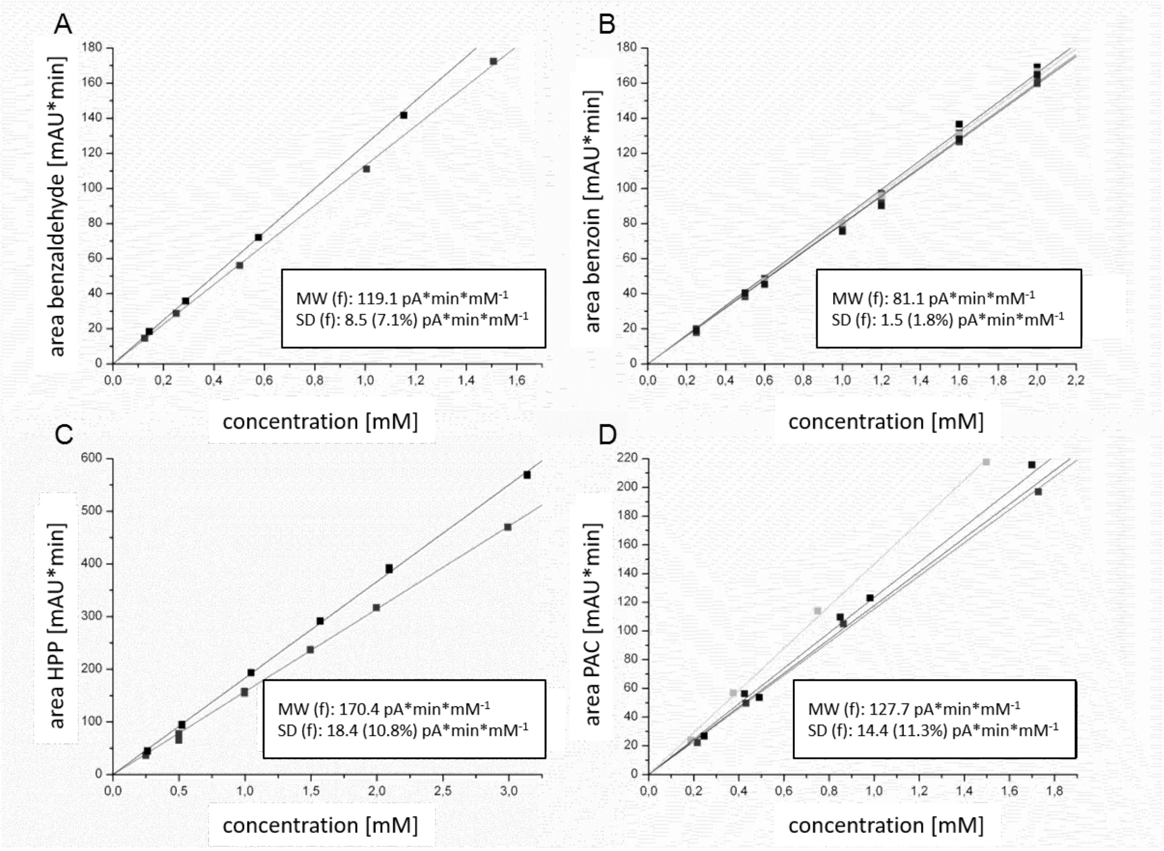


**Figure S9: Calibration of the achiral HPLC for benzaldehyde (A), benzoin (B), HPP (C), and PAC (D).** MW = mean value, f = conversion factor (value correction due to internal standard consideration), SD = absolute root mean square deviation (relative root mean square deviation in %).

**HPLC conditions:**

Column: LiChrospher® 100 RP-8 5 µm (Merck, Darmstadt, Germany)

Eluent: 25% (v/v) Acetonitrile(aq), 60% (v/v) Acetonitrile(aq)

Program:
0 – 12 min, 25% Acetonitrile(aq)
12 – 13 min, gradient: 25% - 60% Acetonitrile(aq)
13 – 20 min, 60% Acetonitrile(aq)
20 – 23 min, gradient: 60% - 25% Acetonitrile(aq)
23 – 25 min, 25% Acetonitrile(aq)

Flow: 1 mL/min

Injection volume: 20 µL

Detection:
λ = 200 nm for PAC, HPP, Benzaldehyde, Benzoin
λ = 250 nm for HPP, Benzaldehyde, Benzoin

Internal standard: 2-Methoxybenzaldehyde

The specified conversion factors (slopes of the calibration lines) refer to a value corrected using the internal standard 2-methoxybenzaldehyde (calculation of the corrected value: area (substance)/area (2-MBA) * mean value of the areas (2-MBA)). This means that a series of measurements is first corrected internally using the mean value of the standard area and then converted into the respective concentrations using the conversion factor f the respective concentrations using the conversion factor f (calculation: corrected area value/conversion factor). In this way, the injection error is corrected from measurement series to series of measurements and the values are adjusted.

#### Figure S10: Calibration of the chiral GC with acetoin.

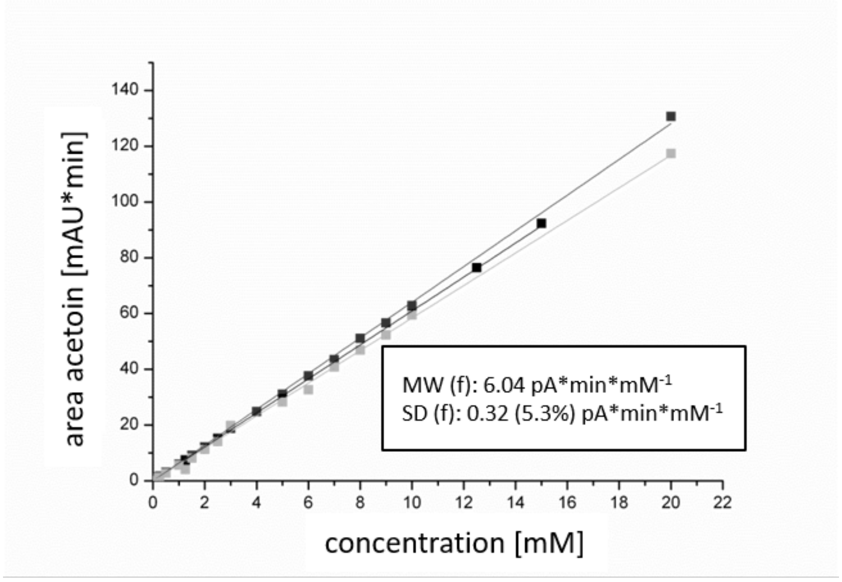


**Figure S10: Calibration of the chiral GC with acetoin.** MW = mean value, f = conversion factor (value correction due to internal standard consideration), SD = absolute root mean square deviation (relative root mean square deviation in %).

**Chiral GC conditions:**

| | |
|---|---|
| Column: | FS-Cyclodex beta-I/P, 50 m x 0.32 mm |
| Carrier gas: | H2 |
| Program: | 0 – 5, 3 min 75 °C |
| | 10 °C/min at 180 °C (3 min hold) |
| Injection volume: | 1 µL (split: 50) |
| Retention times: | (*R*)-Acetoin 5.3 min |
| | (*S*)-Acetoin 5.6 min |
| | Decane 7.5 min |

The acetoin enantiomers were analyzed by chiral GC. For sample preparation, 200 µL of the carboligation mixture was mixed vigorously with 200 µL ethyl acetate (incl. 0.1 µL/mL decane as internal standard). This mixture was then centrifuged for phase separation. Subsequently, 180 µL was sampled from the reaction vial and transferred to a GC vial. 1 µL of this mixture was injected and measured.

The identification and quantification of the individual enantiomers were possible based on prior calibration (**Figure S10**). The chiral GC system was calibrated with acetoin three times independently of each other until the standard deviation of the calibrations from each other was less than 15%.

The specified conversion factor (slope of the calibration line) refers to a value corrected via the internal standard (calculation of the corrected value: area (acetoin)/area (decane) * mean value of the areas (decane)). The mean value of the standard areas was determined from the other values of the measurements in of the measurements in this run to minimize the error of the GC injection.

### Figure S11: Structural dynamics of investigated *Ap*PDC systems.

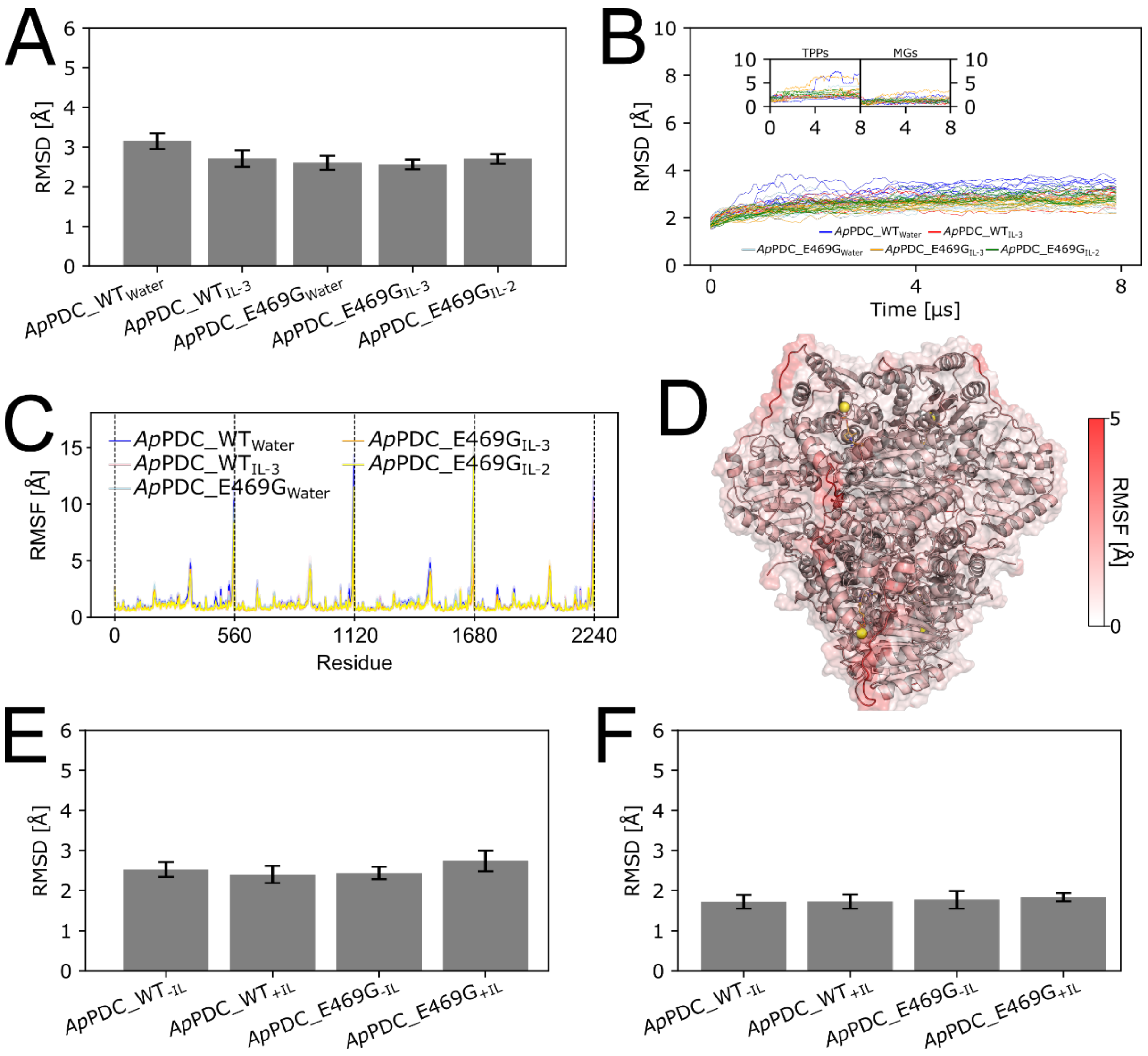


**Figure S11:** Structural dynamics of investigated *Ap*PDC systems in water and ILs. (**A**) RMSD analyses of backbone atoms of *Ap*PDC_WT and *Ap*PDC_E469G in water and in the presence of IL-2 and IL-3 show no significant differences between the systems. (**B**) Evolution of the RMSD over time for all investigated systems. The RMSD of protein backbone atoms is shown for all replicas of each solvent, whereas the RMSD of ThDP non-hydrogen atoms and $Mg^{2+}$ ions, respectively, are only shown for the first replica. (**C**) Analyses of the residue-wise backbone RMSF for all systems indicate that all systems exhibit the same few flexible regions, irrespective of the solvent. (**D**) Mean residue-wise RMSF exemplarily mapped on the structure of *Ap*PDC_WT for the simulation in water (color scale from 0 Å (white) to 5 Å (red)). For clarity, the RMSF values were capped at 5 Å. Structures of ThDP and $Mg^{2+}$ (spheres) indicate the location of active sites. (**E**) RMSD analyses of protein backbone atoms for the 2$^{nd}$ set of unbiased MD simulations of *Ap*PDC_WT and *Ap*PDC_E469G with (+IL) and without (-IL) an IL-3 cation placed in the active sites. (**F**) RMSD of protein backbone atoms for the 3$^{rd}$ set of unbiased MD simulations of *Ap*PDC_WT and *Ap*PDC_E469G with (+IL) and without (-IL) IL-3 cation placed in the active sites. Note for E) and F) that the protein structures of the 2$^{nd}$ and 3$^{rd}$ set are identical and only differ in the state in which ThDP and the substrates are modeled (see Section **2. Materials, methods and computational section in the main manuscript**).

#### Figure S12: IL-2 and IL-3 cations form clusters in solution.

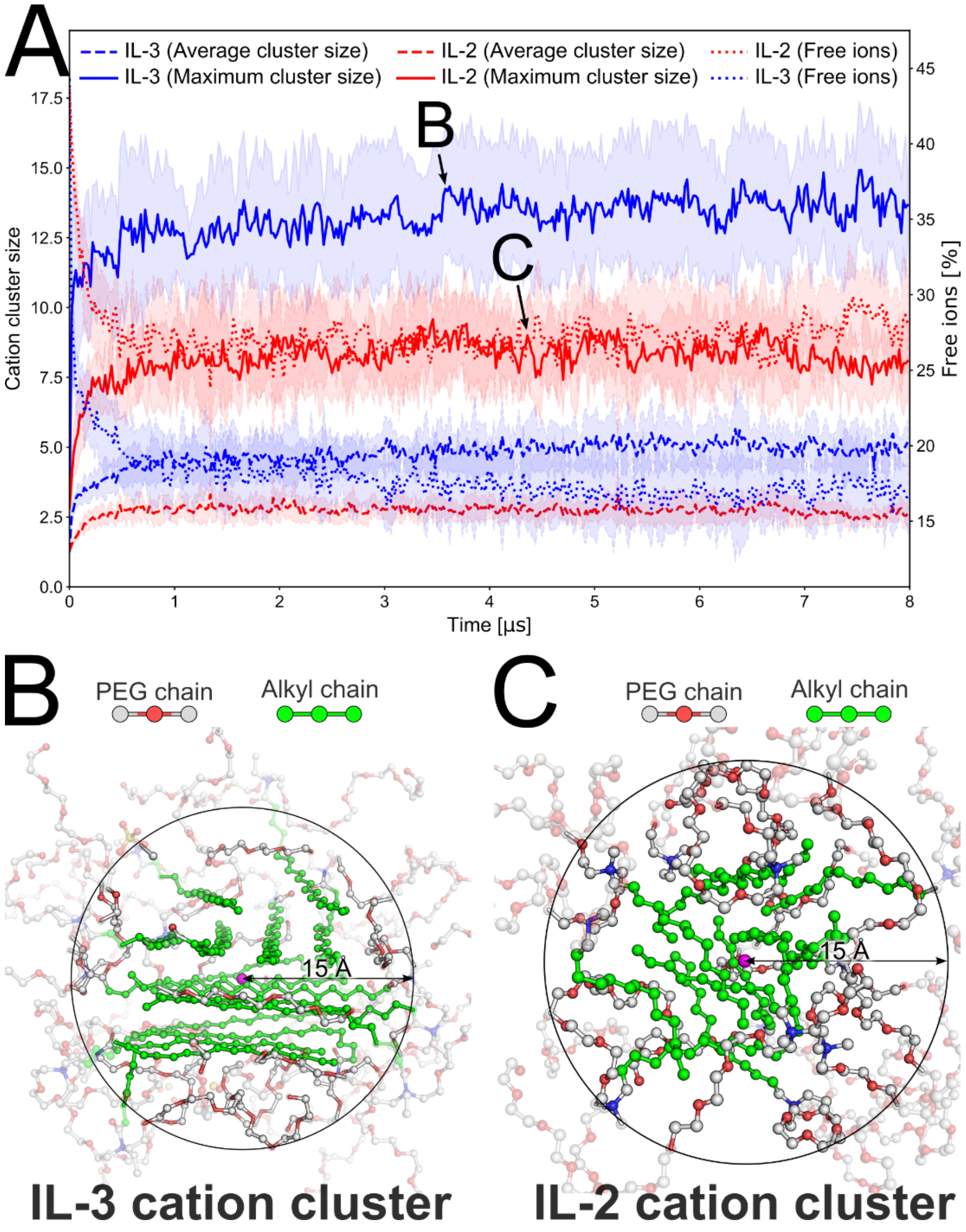


**Figure S12:** IL-2 and IL-3 cations form clusters in solution due to hydrophobic interactions between the alkyl chains. (**A**) Degree of cluster formation for IL-2 and IL-3 cations based on interactions between hydrophobic moieties. Solid and dashed lines depict the maximum and average number of cations in clusters (≥ 2 ions) forming interactions (distance ≤ 5 Å) between two alkyl chain carbons of IL-2 cations (red) and IL-3 cations (blue). Dotted lines depict the percentage of free ions not involved in cation clusters. (**B**) Representative snapshot of an IL-3 cation cluster observed during the MD simulations incorporating 15 cations forming hydrophobic interactions. Alkyl chains of the cations are colored green, hydrophilic moieties are colored white. For clarity, atoms with a distance > 15 Å from the center of the cluster (purple sphere) are drawn transparent. (**C**) Representative snapshot of an IL-2 cation cluster observed during MD simulations incorporating 10 cations forming hydrophobic interactions. Alkyl chains of the cations are colored green, hydrophilic moieties are colored white. For clarity, atoms with a distance larger than 15 Å from the center of the cluster (purple sphere) are drawn transparent.

#### Figure S13: Tunnel dynamics of *Ap*PDC_E469G.

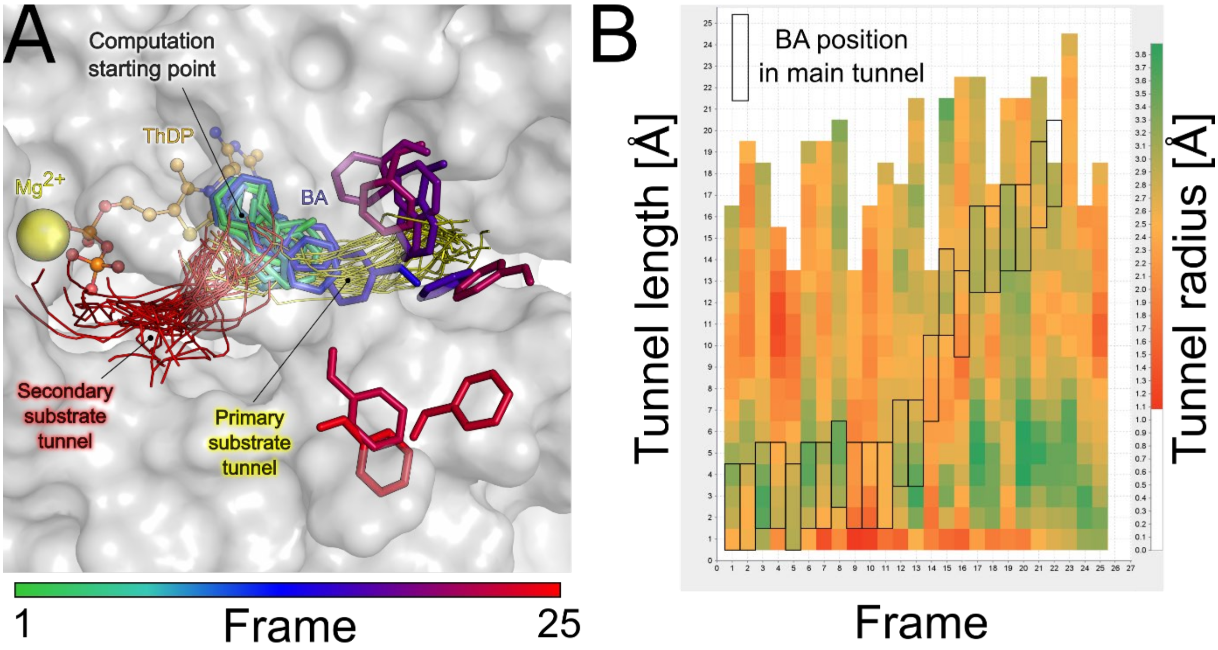


**Figure S13:** Tunnels and tunnel dynamics of *Ap*PDC_E469G in water. (**A**) Process of benzaldehyde (BA) vacating the active site of *Ap*PDC_E469G via the main substrate channel (yellow lines) alongside an existing secondary tunnel (red lines) over the course of 25 frames (50 ns). BA is colored according to the time from green to blue to red for the beginning, middle, and end of the frames, respectively. The tunnels are represented as centerlines for all snapshots. (**B**) Dynamics of the main substrate tunnel of *Ap*PDC_E469G. Data is shown for the tunnel length on the y-axis against the frame number on the x-axis, with data colored according to the tunnel radius from red (narrow) to green (wide). The approximate position of BA along the tunnel over time is indicated via black boxes.

#### Analysis of the molecular structure of Ammoeng 102 (IL-3).

IL-3 (Ammoeng 102 or IoLiLyte T2EG) is an ammonium-based ionic liquid with a quaternary ammonium cation and a sulfate anion. The molecular structure of the cation consists of a linear aliphatic chain (tallow), two polyethylene glycol (PEG) chains, and an ethyl group. The aliphatic tallow chain is derived from tallow fat of vegetable or animal origin, with predominantly saturated or partially unsaturated C16–C18 chains.

The molecular structure is verified by the $^{1}$H- and $^{13}$C-NMR spectra (Figures S14/S15) and the IR spectrum (Figure S16). In the $^{1}$H-NMR spectrum, the signal at δ = 1.18 ppm can be assigned to a majority of the aliphatic tallow chain. The signal at δ = 3.56 ppm is characteristic of the two PEG chains. In the $^{13}$C-NMR, the signals between δ = 25–30 ppm can be assigned mainly to the hydrophobic tallow chain and the signals at δ = 70 ppm to the PEG chains. The IR spectrum shows characteristic signals at 2922 cm$^{-1}$ and 2855 cm$^{-1}$ for symmetric and asymmetric vibrations of the $CH_2$ groups, as well as a strong signal at 1102 cm$^{-1}$ for the C–O–C motif of the PEG chains. The spectra exclude the possibility that the tallow chain is connected to the quaternary nitrogen *via* an amide bond. For an amide bond, a characteristic signal for the carbonyl carbon (C=O) at δ = 175–190 ppm is missing in the $^{13}$C-NMR spectrum, as well as the C=O vibration for amides (Amide I vibration) between 1600–1700 cm$^{-1}$ in the IR spectrum.

#### Figure S14: $^{1}$H-NMR spectra of IL-3.

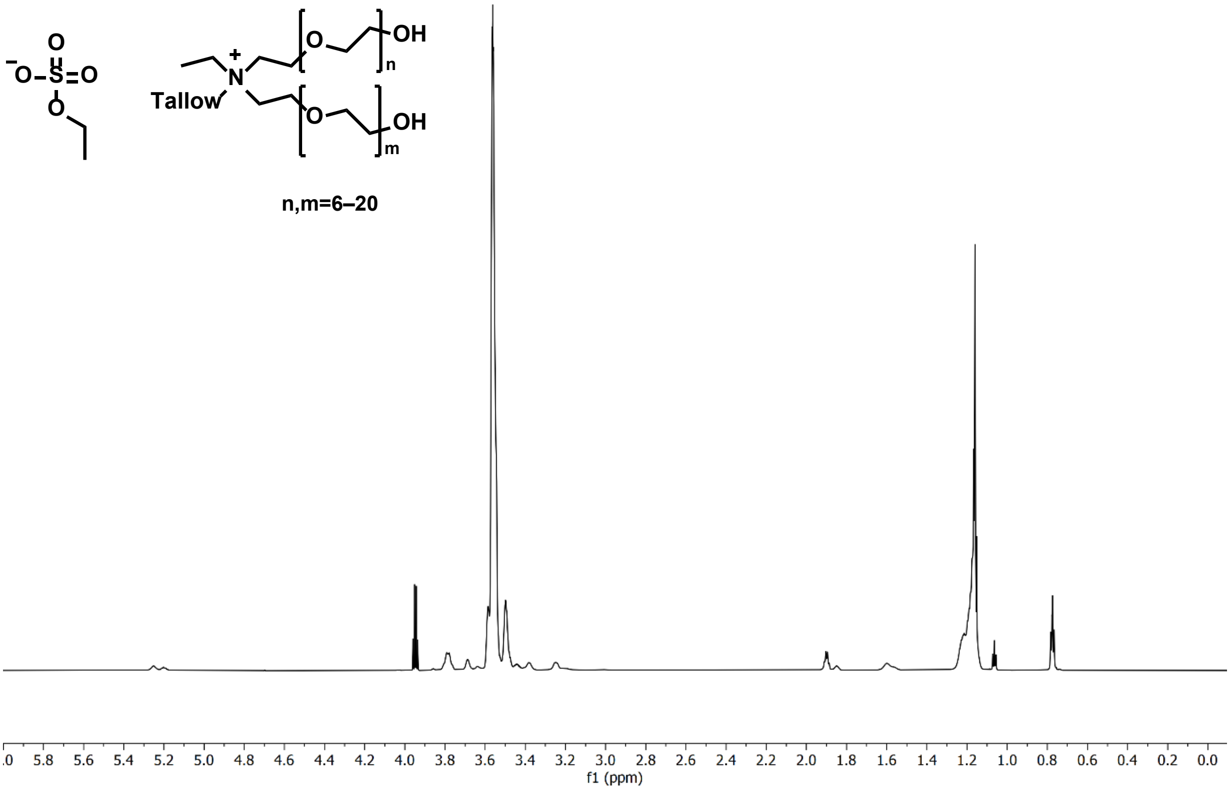


**Figure S14:** $^{1}$H-NMR spectra of IL-3 (IoLiLyte T2EG) in $D_2O$ (900 MHz) at 293 K. $^{1}$H-NMR spectra were recorded at 293.1 K on a Bruker Avance Neo 900 nuclear magnetic resonance spectrometer (900/223 MHz) in $D_2O$ with water suppression. Chemical shifts are given in ppm.

### Figure S15: $^{13}$C-NMR spectra of IL-3

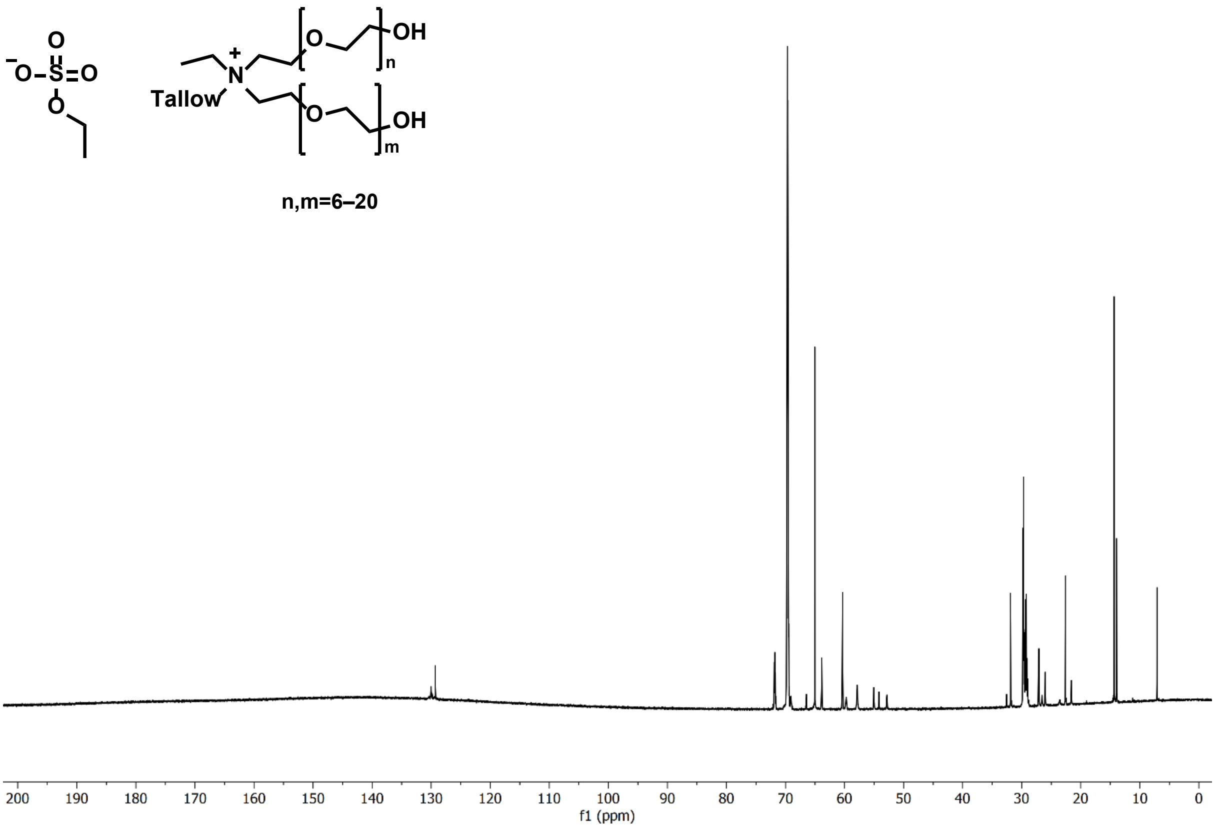


**Figure S15**: $^{1}$H- and $^{13}$C-NMR spectra of IL-3 (IoLiLyte T2EG) were recorded at 293.1 K on a Bruker Avance Neo 900 nuclear magnetic resonance spectrometer (900/223 MHz) in $D_2O$ with the $^{1}$H-NMR spectrum being recorded with water suppression. Chemical shifts are given in ppm.

### Figure S16: IR spectra of IL-3.

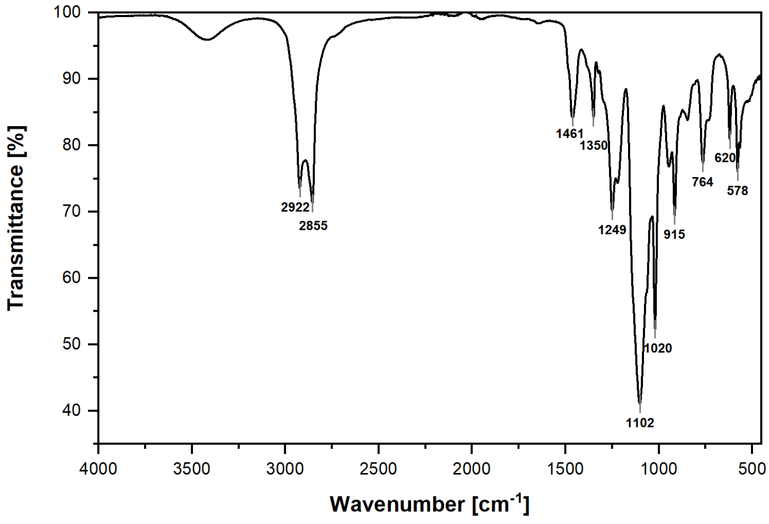


| Peak count | Wavenumber [cm$^{-1}$] | Transmittance [%] |
|---|---|---|
| 1 | 2922.15 | 73.56 |
| 2 | 2854.86 | 71.30 |
| 3 | 1460.50 | 84.28 |
| 4 | 1349.51 | 84.23 |
| 5 | 1248.75 | 70.20 |
| 6 | 1101.70 | 41.08 |
| 7 | 1019.95 | 52.33 |
| 8 | 914.63 | 69.42 |
| 9 | 763.90 | 77.36 |
| 10 | 620.02 | 80.84 |
| 11 | 577.58 | 76.00 |

**Figure S16:** IR spectrum of IL-3. The IR spectrum was recorded at 294 K on a Perkin Elmer Spectrum Two FTIR spectrometer equipped with a $LiTaO_3$ detector and an Attenuated Total Reflection (ATR) module. The sample was measured as a pure substance as a thin film.

#### Figure S17: Solvent dynamics of *Ap*PDC in IL-3 and IL-2.

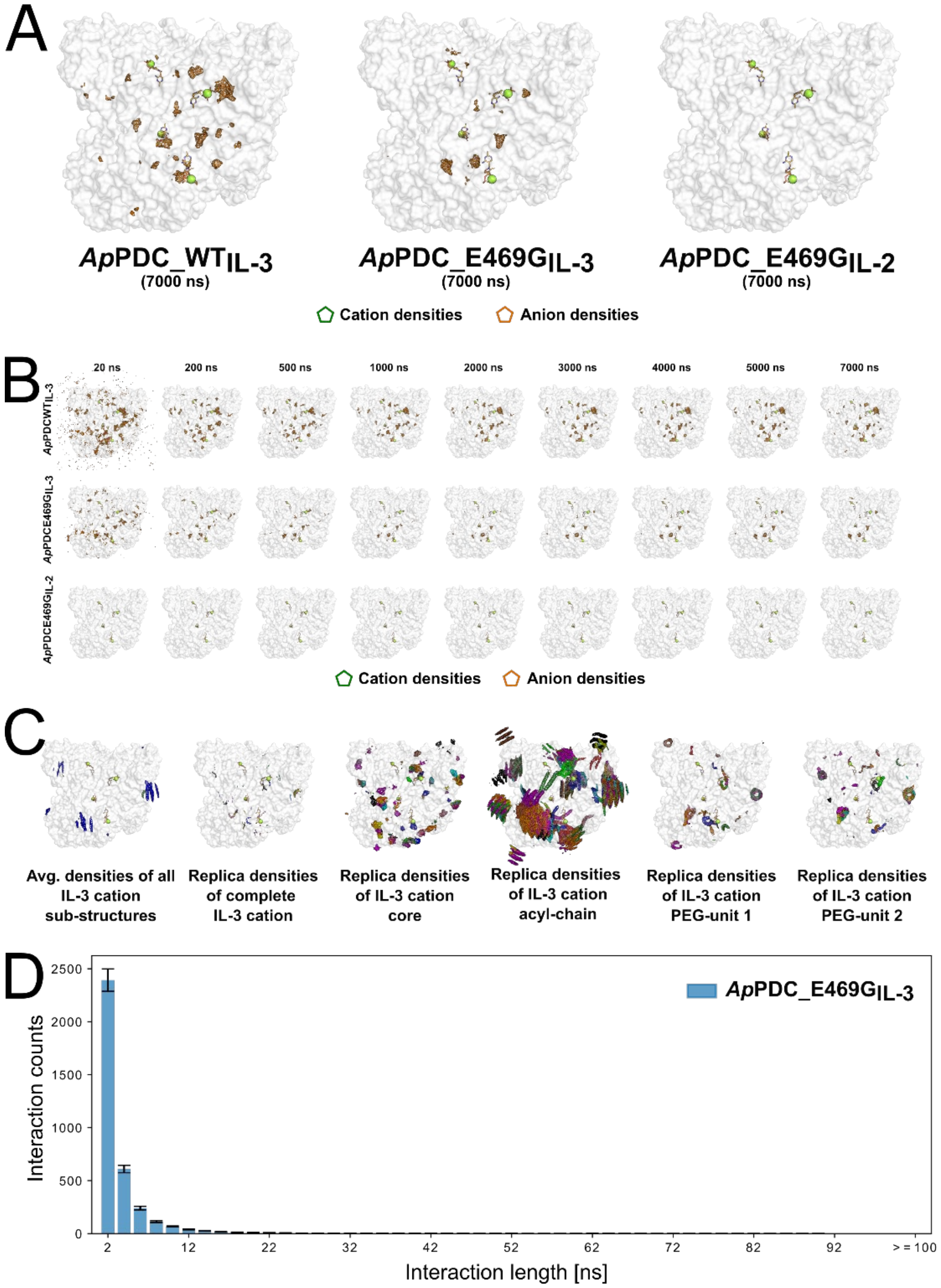


**Figure S17:** Solvent dynamics of *Ap*PDC in the presence of ILs. (**A**) Spatial distribution of solvent molecules around *Ap*PDC_WT and *Ap*PDC_E469G in IL-3 and IL-2 considering the complete simulation time. Regions with a high density of IL cations and anions throughout the MD simulations are shown as green or orange meshes, respectively (cation densities are often too low to be visualized). Structures of ThDP and $Mg^{2+}$ are shown as sticks and spheres. All distributions were normalized according to the number of frames, and σ-values defining the intensity cutoff of the represented data adjusted to the number of heavy atoms were used. (**B**) Evolution of the spatial distribution of solvent molecules throughout the MD simulations, considering 20 ns, 200 ns, 500 ns, 1000 ns, 2000 ns, 3000 ns, 4000 ns, 5000 ns, and 7000 ns, respectively. Regions with a high density of IL cations and anions throughout the MD simulations are shown as green or orange meshes, respectively (cation densities are often too low to be visualized). Structures of ThDP and $Mg^{2+}$ are shown as sticks and spheres. All distributions were normalized according to the number of frames, and σ-values defining the intensity cutoff of the represented data adjusted to the number of heavy atoms were used. (**C**) Spatial distribution of IL-3 cations around *Ap*PDC_E469G resolved on the sub-structure level. Regions with a high density of IL atoms throughout the MD simulations are shown as colored meshes, colored according to substructure (panel 1) or replica (panels 2-6). Structures of ThDP and $Mg^{2+}$ are shown as sticks and spheres. All distributions were normalized according to the number of frames, and σ-values defining the intensity cutoff of the represented data adjusted to the number of heavy atoms were used. (**D**) Interactions of the carbonyl function of IL-3 cation with *Ap*PDC_E469G surface residues according to a distance-based criterion. The total number of interactions is shown for interactions lasting for 1 to 200 ns, respectively. Results are shown as mean ± standard error of the mean (n = 12).

### Figure S18: Structural formula of the IL-3 cation.

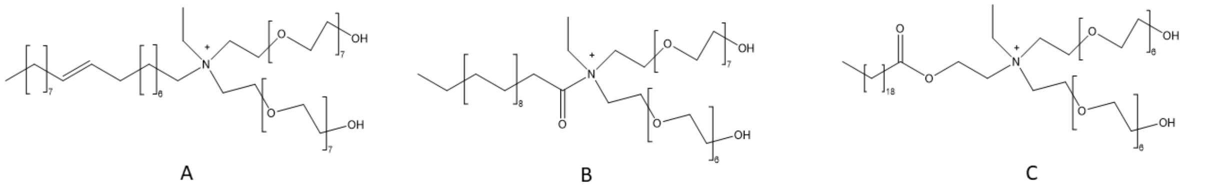


Chemical Formula: $C_{52}H_{106}NO_{16}$
Exact Mass: 1000,7512
Molecular Weight: 1001,4110
m/z: 1000.7512 (100.0%), 1001.7545 (56.2%), 1002.7579 (15.5%), 1002.7554 (3.3%), 1003.7612 (2.8%), 1003.7588 (1.8%), 1001.7574 (1.2%)
Elemental Analysis: C, 62.37; H, 10.67; N, 1.40; O, 25.56

**Figure S18:** Three possible structural formulas matching the m/z ratio of 1000.75078 observed in the FIA-FTICR-MS (see **Figure S8**) (created with ChemDraw Professional 19.1).

### Supplementary results

#### Text S1: Structural dynamics of *Ap*PDC_WT and *Ap*PDC_E469G.

Analyses of the global backbone root-mean-square deviations (RMSD) relative to the crystal structure (PDB ID: 2VBI) [1] revealed that *Ap*PDC_WT and *Ap*PDC_E469G remained structurally largely invariant in water and aqueous IL-2 and IL-3 solutions (RMSD < 3.2 Å) during MD simulations of 8 µs length (**Figure S11A,B)**. Likewise, TPPs and $Mg^{2+}$ ions remained bound to the enzymes (**Figure S11B**). The analyses of the residue-wise root-mean-square fluctuations (RMSF), a measure for atomic mobility, revealed no significant differences between the enzyme variants in the different solvents (**Figure S11C**). Besides a long solvent-exposed loop comprising residues 344-359 distant from the active site as well as the N- and C-terminal regions of the monomers, the remainder of the enzyme showed RMSF values < 2 Å (**Figure S11D)**. The flexible C-terminal helix is part of the substrate tunnel entry, and mobility there might be required for substrate access, as suggested for other enzymes, including the pyruvate decarboxylase from *Zymobacter palmae* [2,3]. These results indicate that *Ap*PDC_WT and *Ap*PDC_E469G remain globally structurally invariant in all solvents and that global changes in the structural dynamics cannot be responsible for the shift in stereoselectivity. For the second and third sets of unbiased MD simulations involving systems of *Ap*PDC_WT and *Ap*PDC_E469G, we observed similar behavior with no significant differences regarding the RMSD or RMSF (**Figure S11E,F**).

#### Text S2: Binding distribution of different functional groups.

Over all replicas and active sites, we observed in total 91 incidents (11 cations/80 anions), 80 incidents (24 cations/56 anions), and 82 incidents (76 cations/6 anions) of IL ions within 5 Å of the active site cavity for *Ap*PDC_WT in IL-3, *Ap*PDC_E469G in IL-3, and *Ap*PDC_E469G in IL-2, respectively.

The IL-3 cation visited the active site less frequently than the IL-2 cation, which can be explained by a lower effective concentration of free IL-3 due to a stronger aggregation in the solvent (see **Text S3**). Both cations interacted with the enzymes with their hydrophobic and hydrophilic moieties. However, mainly hydrophobic enzyme residues were involved in recruiting and guiding the IL cation chains through the tunnel and in subsequent interactions within the active site (see inset of **Figure 8A** of the main manuscript). Both cations showed roughly similar interaction frequencies of the hydrophilic and hydrophobic moieties to the active site cavities of *Ap*PDC. This might indicate a higher affinity of the hydrophobic side chain to the substrate tunnel, as the effective concentration of hydrophilic chains is higher compared to the hydrophobic ones. In detail, the hydrophobic moiety accounted for 5/11 (45.5%), 9/24 (37.5%), and 31/76 (40.8%) of the visits for *Ap*PDC_WT in IL-3, *Ap*PDC_E469G in IL-3, and *Ap*PDC_E469G in IL-2, respectively. Consequently, interactions involving the two hydrophilic PEG-moieties of the IL cations account for 6/11 (54.5%), 15/24 (62.5%), and 45/76 (59.2%) of the visits for *Ap*PDC_WT in IL-3, *Ap*PDC_E469G in IL-3, and *Ap*PDC_E469G in IL-2, respectively. 5/11, 12/24, and 52/76 cations unbound again during the MD simulations (defined as reaching a distance > 15 Å to the active site cavity after the binding event occurred, corresponding to vacating the substrate tunnel). Although we did not observe (multiple) rebinding of a cation and, thus, did not reach a binding equilibrium, the unbinding events indicate that the cations are not kinetically trapped in the active sites. Of these interactions, 3 and 6 reached a distance < 5 Å to the center of the (*S*)-pocket in IL-3 and IL-2, respectively, with 1/3 (2/3) or 4/6 (2/6) interactions involving the hydrophobic (hydrophilic) moiety. Notably, hydrophobic interactions of IL-2 with the active site seem to require a sterically hindered deeper penetration of the cation center towards the entry of the substrate access tunnel due to the shorter alkyl chain length, whereas this is not necessary for IL-3 interactions, indicating that this structural difference is a potential origin for the larger effect on stereoselectivity of IL-3 [4,5]. Complete penetration of the cation core into the active site cavity of *Ap*PDC was not observed for either cation.

As to anions, we also observed substantial differences in interaction frequencies and durations. The ethyl sulfate anion of IL-3 visited the active site more frequently but shorter than the cations, often followed by leaving the active site completely. In detail, the ethyl sulfate anion of IL-3 reached the active site cavity 80 and 56 times for *Ap*PDC_WT and *Ap*PDC_E469G in IL-3, respectively. Of these interactions, 57 (71%) and 44 (79%) anions unbound again during the MD simulations, most often within 500 ns. Of all interactions for *Ap*PDC_E469G in IL-3, 9 events reached a distance < 5 Å to the center of the (*S*)-pocket after entering the active site cavity, typically involving the anions alkyl moiety. Rarely, anions remained in the active site for larger parts of the MD simulations, particularly, when stabilized by the presence of IL cations. By contrast, the chloride anion of IL-2 showed only six visits overall, which typically lasted for < 100 ns before completely unbinding.

Finally, to investigate the relevance of the putative carbonyl function of the IL-3 cation on the observed stereoselectivity shifts of *Ap*PDC_E469G in IL-3, we investigated the binding of the carbonyl function to *Ap*PDC_E469G by computing interactions of the carbonyl oxygen atom with surface residues (see **Figure S17D**). Here, to probe for particularly strong and stable interactions, binding events were determined by assessing interaction distances < 4 Å to neighboring heavy atoms. Additionally, we related the observations with events of IL-3 cations entering the active site cavity as outlined above. Generally, the carbonyl function was found to interact mostly weakly and infrequently, and not specifically with surface residues of *Ap*PDC_E469G (see **Figure S17** for a depiction of the spatial distribution of ions around *Ap*PDC_E469G). Penetration of the cation center, involving the carbonyl function, into the active site was not observed. We observed on average 3563.67 ± 193.32 interactions per replica (in total, 42 764 interactions), of which 67.15% or 96.01% had a duration of ≤ 2 ns (1 frame) or ≤ 10 ns (≤ 5 frames), respectively, indicating that these interactions by themselves should not significantly affect the overall structure of both *Ap*PDC variants. In total, only six cations formed interactions longer than 100 ns, with a maximum interaction duration of 504 ns. For these interactions, visual inspection revealed that the cations were stabilized via additional interactions of the IL-3 side chains with *Ap*PDC and typically not by the cation core structure alone. Furthermore, none of these interactions were formed by carbonyl functions of cations that eventually entered the active site cavity. Overall, these observations indicate that the carbonyl function itself does not play a significant role in the stereoselectivity shifts of *Ap*PDC_E469G in IL-3.

#### Text S3: Cation clusters in IL-3 reduce the degree of free ions in solution.

In view of the higher proportion of IL-2 cations binding to the active site compared to IL-3 cations, we investigated the intra- and intermolecular dynamics of these cations in the respective solvents (**Figure S12)**. Interestingly, both cations aggregate due to intermolecular hydrophobic interactions of the alkyl moieties. For the IL-3 cation, structured and layered clusters involving ≥ 15 cations were observed, whereas the IL-2 cation formed less structured and less dense clusters of typically ≤ 10 cations (**Figure S12A)**. Representative snapshots are shown in **Figure S12B/C**. This finding is in line with the general amphiphilic nature of Ammoeng ILs and experimental or computational studies of other ILs with cations comprising long aliphatic side chains [6–8], such as $C_nMIM$ ($n \geq 8$). Particularly for IL-3, a prior study of Ribot et al. [9] reported ordered IL-3 structures in SAXS experiments, albeit this only occurred in higher concentrations above 40 wt%. The cation clusters likely lower the effective concentration of free alkyl chains (dotted lines in **Figure S12A**) that can enter the active site [8], which can explain the higher proportion of binding of IL-2.

#### Text S4: Tunnel analysis.

To evaluate whether the IL-3 cation binding still allows substrates and products to access and exit from the active site, we applied the CAVER software [10] to identify and evaluate tunnels regarding crucial structural parameters, such as the average or maximum bottleneck radii. Note that we report the maximum in contrast to the minimum bottleneck radius, as the tunnels were found to be flexible rather than rigid, where the latter would be the more important metric. In all simulations and solvents, we observed a tunnel connecting the active site of *Ap*PDC_WT and *Ap*PDC_E469G to the exterior of the enzyme, which resembled the main substrate tunnel found in the crystal structure. Additionally, we investigated structures of *Ap*PDC_E469G and *Ap*PDC_WT with and without an IL-3 cation present in the main substrate tunnel, hereafter referred to as *Ap*PDC_E469G$_{+IL}$, *Ap*PDC_WT$_{+IL}$ (+IL = IL-3 cation placed in the active site), *Ap*PDC_E469G$_{-IL}$, and *Ap*PDC_WT$_{-IL}$ (-IL = no IL-3 cation present). All replicas of *Ap*PDC_E469G$_{+IL}$ revealed a secondary - potentially temporarily, yet frequently occurring - tunnel similar to the main substrate tunnel, i.e., adjacent to the C-terminal helix (see **Figure S13**), which connected the active site to the solvent. We observed no statistically significant differences between the secondary tunnels observed in *Ap*PDC_E469G$_{+IL}$, *Ap*PDC_E469G$_{-IL}$, *Ap*PDC_WT$_{-IL}$ and the tunnel found in the crystal structure of *Ap*PDC_WT in terms of average or maximum bottleneck radii, suggesting that substrate access and product egress are still possible. Interestingly, some MD simulations showed a long (water) tunnel connecting the two active sites of an *Ap*PDC dimer (**Figure 8D of the main manuscript**), which was shown previously for other pyruvate decarboxylases [11,12] but not yet for *Ap*PDC.

Furthermore, we investigated a sequence of 25 frames (50 ns) of *Ap*PDC_E469$_{-IL}$, in which benzaldehyde left the active site via the main substrate tunnel (**Figure S13A**), which was present in all frames with average and maximum bottleneck radii of 1.64 ± 0.44 Å and 2.41 Å, respectively. In all snapshots, the secondary substrate tunnel was present with average and maximum bottleneck radii of 1.54 ± 0.33 Å and 2.26 Å, respectively, further indicating the existence of an alternate exit route in case of a blocked main substrate tunnel. By following the structural features of the tunnel over time (**Figure S13B**), i.e., the length and radius, we related the presence of benzaldehyde (BA) (black rectangles in **Figure S13B**) at a given tunnel location with the tunnel radius at that point. In general, the tunnel was most spacious at the tunnel start, i.e., within the active site, and became narrower, with the narrowest point located approximately at the tunnel center. Interestingly, the position of BA within the tunnel during the translocation to the exterior coincided strongly with increased tunnel radii. However, the tunnel was also able to adapt to widened states (see the first tunnel half in frames 19-25) without BA being present, such that it is unclear whether BA can induce a tunnel widening or rather spontaneously follows widened states of the substrate tunnel.

To conclude, our analyses revealed I) the existence of alternate substrate tunnels that might allow the exit of the substrate in case of an occupied main substrate tunnel by IL cations and II) an inherent flexibility of the main and secondary substrate tunnels, potentially accommodating BA molecules during the substrate egress processes.

#### Text S5: Structural assessment of the IL-3 cation structure.

During our studies, we raised doubts regarding the manufacturer-proposed structure (sold as IoLiLyte T2EG) of the IL-3 cation (see **Figure 2 in the main manuscript**) [13]. It is an ammonium-based ionic liquid with a quaternary ammonium cation and a sulfate anion. The molecular structure of the cation consists of a linear aliphatic chain (tallow), two polyethylene glycol (PEG) chains, and an ethyl group. The structure, as given by the manufacturer, suggests a direct bond of the tallow residue to the central quaternary nitrogen. Regarding the tallow component, its chemical composition can vary depending on the animal source but comprises mainly non-, mono-, or di-unsaturated fatty acids of varying carbon chain lengths (mainly C16-C18), sometimes with minor impurities such as cholesterol. The most abundant components, palmitic acid (C16:0), stearic acid (C18:0), and oleic acid (C18:1ω9), typically account for approximately 23-31%, 11-25%, and 24-47% of the mass fraction, respectively, with oleic acid being the largest fraction (**Figure S18A**) [14–16]. These components usually represent ≥ 90% of tallow [14–16]. However, according to other studies [9,17–21]) the cation structure features a carbonyl group adjacent to the positively charged nitrogen atom, which we deemed potentially prone to hydrolysis in aqueous media (**Figure S18B**). An alternative structure was proposed by Schindl et al. [22], in which the aliphatic residue is linked to the central quaternary nitrogen as ethyl ester (**Figure S18C**; note, however, that the number of repeating units differ from those given by Schindl et al. [22] in their Figure 2). Unfortunately, Schindl et al. provided no direct evidence for this proposed structure. All structural formula shown in **Figure S18** match the m/z ratio of 1000.75078 observed in the FIA-FTICR-MS (**Figure S8**). The expected molar mass of the closest Schindl structure following Schindl et al. [22] (1017.39 u; calculated using Avogadro v1.9.9 [23,24]) does not match the main peaks observed in our mass spectrometry experiments, which were around 1000.75 u (**Figure S8**).

To address these concerns, we compiled and analyzed the existing body of literature and information from relevant patents (cited above), and verified the molecular structure integrating data from FIA-FTICR-MS analyses (**Figure S8**), $^{1}$H- and $^{13}$C-NMR spectra (**Figures S14 and S15**), and an IR spectrum (**Figure S16**), challenging previously suggested structures.

The $^{1}$H-HMR experiment yielded a spectrum (**Figure S14**) similar to that reported by Ribot et al. [9]. Here, the signal at δ = 1.18 ppm can be assigned to a majority of the aliphatic tallow chain (**Figure S14)**. The signal at δ = 3.56 ppm is characteristic of the two PEG chains. In the $^{13}$C-NMR, the signals between δ = 25−30 ppm can be assigned mainly to the hydrophobic tallow chain and the signals at δ = 70 ppm to the PEG chains (**Figure S15)**. The IR spectrum shows characteristic signals at 2922 cm$^{-1}$ and 2855 cm$^{-1}$ for symmetric and asymmetric vibrations of the $CH_2$ groups, as well as a strong signal at 1102 cm$^{-1}$ for the C−O−C motif of the PEG chains (**Figure S16)**. The spectra exclude the possibility that the tallow chain is connected to the quaternary nitrogen *via* an amide bond. For an amide bond, a characteristic signal for the carbonyl carbon (C=O) at δ = 175−190 ppm is missing in the $^{13}$C-NMR spectrum, as well as the C=O vibration for amides (Amide I vibration) between 1600−1700 cm$^{-1}$ in the IR spectrum.

This discrepancy suggests that the actual structure of the IL-3 cation may still differ from both the manufacturer's description and the variations proposed in the literature. Notably, the mass spectrometry analyses did not indicate any hydrolytic decomposition into trialkylamine and fatty acid derivatives: the hypothetical IL-3 hydrolysis products (i.e., the trialkylamine with only the ethyl substituent and two poly-PEG chains) could be fitted only up to a molecular weight of 1014.24 u (compared to observed 1000.75 u) or 661.82 u (compared to observed 648.54 u), which then incorporate 20 or 12 PEG units, respectively. This observation, combined with the absence of typical signals for the carbonyl function in the $^{13}C$ and IR spectra, could indicate that there is no carbonyl group present in the cation.

To further elucidate the composition of the PEG and alkyl chains, we analyzed the molecular weight distribution of the PEG units. PEG is typically synthesized by ring-opening polymerization of ethylene oxide, which often results in a polydisperse distribution of molecular weights, ranging from a few hundred u to several hundred thousand u, in increments of +44 u, i.e., single PEG units. While monodisperse PEG fractions are commercially available, the presence of multiple mass changes of +44 u in the mass spectra indicates that a polydisperse PEG product was used in the synthesis of the IL-3 cation.

Based on this data, we identified specific mass change patterns in our mass spectrometric analyses that correlate with substitutions in the tallow and PEG chains:

| **Change in mass (Δu)** | **Substitution pattern** |
|---|---|
| +2 | C18:1 -> C18:0 |
| +14 | $+CH_2$ |
| **+18** | +PEG & C18:1->C16:0 |
| **+26** | C16:0 -> C18:1 |
| +28 | C16:0 -> C18:0 |
| **+44** | +PEG |
| +62 | +2 PEG & C18:1->C16:0 |
| +70 | +PEG & C16:0->C18:1 |

By examining the substitution patterns in **Figure S8**, we gained insights into the structural features of the main mass peak, particularly the composition of the PEG and alkyl chains. The mass changes of +18 u, +26 u, and +44 u for the peaks 912 -> 930 u, 930 -> 956 u, and 956 -> 1000 u suggest specific substitutions: from a C18:1 to a C16:0 fatty acid with an additional PEG unit, a reverse substitution from C16:0 to C18:1, and the addition of another PEG unit, respectively. Consequently, the main structure at the 1000 u peak likely comprises a mono-unsaturated C18:1 alkyl chain. Smaller peaks adjacent to the main peak indicate the presence of cations with the same number of PEG units but different alkyl chains.

Notably, the gradual decrease in mass and mass fraction from 1000 u to 648 u suggests a reduction in PEG units, indicating that the IL-3 cation contains at least 8 PEG units, albeit the molar weight around 1000.75 u (assuming a tallow acyl- or alkyl-group) rather points to a total of 14 PEG units. However, a novel peak at 602 u (compared to 604 u) with a significantly higher mass fraction indicates either a shift in the dominant molecular distribution or the presence of different molecular species. This new pattern of decreasing PEG units appears to break again at the 468 u mass peak, deviating from the expected 470 u peak.

Taking the above information into account, a plausible IL-3 cation structure of the main peak fraction might be the IL-3 cation with tallow representing a *cis*-9-monounsaturated alkyl-group in contrast to an acyl-group, resembling Ammoeng 101 with the cocos alkyl-group and with

14 PEG moieties in total. While the synthesis pathway is unknown, this structure would also likely be stable and result in an m/z ratio of 1000.75. This is more in line with observations in the mass spectrometry spectra, although a final validation remains to be done. Due to the still elusive real structure, we decided to adapt an IL-3 constitution (**Figure S18A**) that is in line with the results from the mass-spectrometry analyses, the distribution of the individual tallow- and PEG-components, and previous studies.

To conclude, despite the new insights and data provided in this study, it must be acknowledged that a definitive structure for the IL-3 cation, particularly the core structure, remains uncertain. The findings presented here contribute to the ongoing discussion, yet the true structure of the ion still awaits full elucidation. Further research, including advanced spectroscopic techniques and theoretical modeling, as well as knowledge of the applied synthesis route, will be required to resolve this structural uncertainty.

#### Text S6: Solvent dynamics at the *Ap*PDC surface.

To investigate the dynamics of solvent molecules around the *Ap*PDC surface and to identify potential interaction sites, we computed the spatial distribution of all solvent molecules. Overall, the analysis of the spatial density reveals interesting insights into the interactions of the solvent molecules and their functional groups with *Ap*PDC.

In detail, density maps of solvent molecules averaged across all replicas of the MD simulations of *Ap*PDC_WT in IL-3, *Ap*PDC_E469G in IL-3, and *Ap*PDC_E469G in IL-2 only revealed interaction sites of the IL-3 sulfate anion at the protein surface, whereas the IL-2 or IL-3 cations and the chloride anion did not show specific interaction sites (see **Figure S17A**). This was also evident when evaluating the evolution of average spatial densities throughout our simulations, where again only sulfate anion densities were observed. Overall, simulation times of 500 ns to 1000 ns were required to achieve converged results (**Figure S17B**), in line with previous results for interactions of the structurally similar trifluoromethane sulfonate with *Bs*LipA [25].

Interestingly, for the IL-3 cation, density analyses of the substructures – with cut-off values scaled down to match the respective approximated substructure volume – for the individual replica, revealed multiple interaction sites of various cation moieties on the protein surface (**Figure S17C**), supporting the existence of different affinities of the substructures for interactions to *Ap*PDC surface residues. The observed substructure densities were particularly strong for the alkyl chain, where clusters of aliphatic side chains observed for IL-3 (see **Text S3** and **Figure S12**) likely result in localized ion clusters with lower mobility, which, in turn, leads to more pronounced spatial densities. Interestingly, when averaged over all replica, almost no densities of the PEG chains as well as the ion core group remain visible despite dominant localizations in the individual replica, indicating disperse and less specific, interactions of these polar moieties with *Ap*PDC (**Figure S17C**).

Overall, our results indicate that IL ions can interact with the *Ap*PDC surface residues at specific interaction sites. The substantial differences in the observed ion densities around the *Ap*PDC surface further hint to differences in the affinities of ion substructures regarding the preferential interactions with enzyme residues, where particularly the aliphatic moiety of Ammoeng 102 showed dominant density accumulations at and around the *Ap*PDC surface when compared to the polar or core regions of Ammoeng 102. In contrast, the chloride anion of Ammoeng 101 showed weak binding to *Ap*PDC surface residues, in line with previous reports binding studies for inorganic IL ions to enzyme surface residues [26].

#### References:


[1] Gocke D, Walter L, Gauchenova E, Kolter G, Knoll M, Berthold CL, et al. Rational protein design of ThDP-dependent enzymes: engineering stereoselectivity. ChemBioChem 2008;9:406–12. https://doi.org/10.1002/cbic.200700598.

[2] Becker D, Bharatam P V., Gohlke H. F/G Region Rigidity is Inversely Correlated to Substrate Promiscuity of Human CYP Isoforms Involved in Metabolism. J Chem Inf Model 2021;61:4023–30. https://doi.org/10.1021/acs.jcim.1c00558.

[3] Buddrus L, Andrews ES V, Leak DJ, Danson MJ, Arcus VL, Crennell SJ. Crystal structure of pyruvate decarboxylase from *Zymobacter palmae*. Acta Crystallogr Sect F Struct Biol Commun 2016;72:700–6. https://doi.org/10.1107/S2053230X16012012.

[4] Dalhaimer P, Blankenship KR. All-Atom Molecular Dynamics Simulations of Polyethylene Glycol (PEG) and LIMP-2 Reveal That PEG Penetrates Deep into the Proposed CD36 Cholesterol-Transport Tunnel. ACS Omega 2022;7:15728–38. https://doi.org/10.1021/acsomega.2c00667.

[5] Munasinghe A, Mathavan A, Mathavan A, Lin P, Colina CM. PEGylation within a confined hydrophobic cavity of a protein. Phys Chem Chem Phys 2019;21:25584–96. https://doi.org/10.1039/c9cp04387j.

[6] Singh T, Kumar A. Aggregation Behavior of Ionic Liquids in Aqueous Solutions : Effect of Alkyl Chain Length , Cations , and Anions. J Phys Chem B 2007;111:7843–51. https://doi.org/10.1021/jp0726889.

[7] Chen S, Zhang S, Liu X, Wang J, Wang J, Dong K, et al. Ionic liquid clusters: Structure, formation mechanism, and effect on the behavior of ionic liquids. Phys Chem Chem Phys 2014;16:5893–906. https://doi.org/10.1039/c3cp53116c.

[8] Liu J, Wang Y, Huo F, He H. Ionic liquids inhibit the dynamic transition from ?-helices to ?-sheets in peptides. Fundam Res 2024;4:777–84. https://doi.org/10.1016/j.fmre.2023.12.013.

[9] Ribot JC, Guerrero-sanchez C, Greaves TL, Kennedy DF, Hoogenboom R, Schubert US. Amphiphilic oligoether-based ionic liquids as functional materials for thermoresponsive ion gels with tunable properties via aqueous gelation. Soft Matter 2012;8:1025–32. https://doi.org/10.1039/c1sm06468a.

[10] Chovancova E, Pavelka A, Benes P, Strnad O, Brezovsky J, Kozlikova B, et al. CAVER 3.0: A Tool for the Analysis of Transport Pathways in Dynamic Protein Structures. PLoS Comput Biol 2012;8:23–30. https://doi.org/10.1371/journal.pcbi.1002708.

[11] Zyl LJ Van, Schubert W, Tuffin MI, Cowan DA. Structure and functional characterization of pyruvate decarboxylase from *Gluconacetobacter diazotrophicus*. BMC Res Notes 2014;14:21.

[12] Pei X, Erixon KM, Luisi BF, Leeper FJ. Structural Insights into the Prereaction State of Pyruvate Decarboxylase. Biochemistry 2010;49:1727–36. https://doi.org/10.1021/bi901864j.

[13] IoliTec GmbH. https://iolitec.de/en/node/487 n.d. https://iolitec.de/en/node/487.

[14] Fat Content and Composition of Animal Products. Proc. a Symp. Washingt. D.C., Dec. 12 -13, 1974, National Research Council (U.S.); 1976, p. 203.

[15] Panneer Selvam DJ, Vadivel K. Performance and emission analysis of DI diesel

engine fuelled with methyl esters of beef tallow and diesel blends. Procedia Eng 2012;38:342–58. https://doi.org/10.1016/j.proeng.2012.06.043.

[16] Luddy FE, Hampson JW, Herb SF, Rothbart HL. Physiochemically designed fat compositions from tallow. US4130572A, 1977.

[17] Kohlmann C, Robertz N, Leuchs S, Dogan Z, Lütz S, Bitzer K, et al. Ionic liquid facilitates biocatalytic conversion of hardly water soluble ketones. J Mol Catal B Enzym 2011;68:147–53. https://doi.org/10.1016/j.molcatb.2010.10.003.

[18] Yang J, Pérez B, Anankanbil S, Li J, Gao R, Guo Z. Enhanced synthesis of alkyl galactopyranoside by thermotoga naphthophila β-galactosidase catalyzed transglycosylation: Kinetic insight of a functionalized ionic liquid-mediated system. ACS Sustain Chem Eng 2017;5:2006–14. https://doi.org/10.1021/acssuschemeng.6b02862.

[19] Guo Z, Chen B, Murillo RL, Tan T, Xu X. Functional dependency of structures of ionic liquids: Do substituents govern the selectivity of enzymatic glycerolysis. Org Biomol Chem 2006;4:2772–6. https://doi.org/10.1039/b606900b.

[20] Thermodynamics JC, Pereiro AB, Rodriguez A. Application of the ionic liquid Ammoeng 102 for aromatic / aliphatic hydrocarbon separation. J Chem Thermodyn 2009;41:951–6. https://doi.org/10.1016/j.jct.2009.03.011.

[21] Guo Z, Kahveci D, Beraat Ö, Xuebing X. Improving enzymatic production of diglycerides by engineering binary ionic liquid medium system. N Biotechnol 2009;26:37–43. https://doi.org/10.1016/j.nbt.2009.04.001.

[22] Schindl A, Hagen ML, Muzammal S, Gunasekera HAD, Croft AK. Proteins in ionic liquids: Reactions, applications, and futures. Front Chem 2019;7:347. https://doi.org/10.3389/fchem.2019.00347.

[23] Avogadro: an open-source molecular builder and visualization tool. Version 1.99 n.d. http://avogadro.cc/.

[24] Hanwell MD, Curtis DE, Lonie DC, Vandermeersch T, Zurek E, Hutchison GR. Avogadro : an advanced semantic chemical editor , visualization , and analysis platform. Cheminformatics 2012;4:17.

[25] El Harrar T, Frieg B, Davari MD, Jaeger KE, Schwaneberg U, Gohlke H. Aqueous ionic liquids redistribute local enzyme stability via long-range perturbation pathways. Comput Struct Biotechnol J 2021;19:4248–64. https://doi.org/10.1016/j.csbj.2021.07.001.

[26] El Harrar T, Gohlke H. Cumulative Millisecond-Long Sampling for a Comprehensive Energetic Evaluation of Aqueous Ionic Liquid Effects on Amino Acid Interactions. J Chem Inf Model 2023;63:281–98. https://doi.org/10.1021/acs.jcim.2c01123.
